# Gene protein sequence evolution can predict the rapid divergence of ovariole numbers in *Drosophila*

**DOI:** 10.1101/2023.09.03.556080

**Authors:** Carrie A. Whittle, Cassandra G. Extavour

## Abstract

Ovaries play key roles in fitness and evolution: they are essential female reproductive structures that develop and house the eggs in sexually reproducing animals. In *Drosophila*, the mature ovary contains multiple tubular egg-producing structures known as ovarioles. Ovarioles arise from somatic cellular structures in the larval ovary called terminal filaments, formed by terminal filament cells and subsequently enclosed by sheath cells. As in many other insects, ovariole number per female varies extensively in *Drosophila*. At present however, there is a striking gap of information on genetic mechanisms and evolutionary forces that shape the well-documented rapid interspecies divergence of ovariole numbers. To address this gap, here we studied genes associated with *D. melanogaster* ovariole number or functions based on recent experimental and transcriptional datasets from larval ovaries, including terminal filaments and sheath cells, and assessed their rates and patterns of molecular evolution in five closely related species of the *melanogaster* subgroup that exhibit species-specific differences in ovariole numbers. From comprehensive analyses of protein sequence evolution (dN/dS), branch-site positive selection, expression specificity (*tau*) and phylogenetic regressions (PGLS), we report evidence of 42 genes that showed signs of playing roles in the genetic basis of interspecies evolutionary change of *Drosophila* ovariole number. These included the signalling genes *upd2* and *Ilp5* and extracellular matrix genes *vkg* and *Col4a1*, whose dN/dS predicted ovariole numbers among species. Together, we propose a model whereby a set of ovariole-involved gene proteins have an enhanced evolvability, including adaptive evolution, facilitating rapid shifts in ovariole number among *Drosophila* species.

**Significance Statement:** Ovaries in *Drosophila*, like in other insects, contain egg producing structures, known as ovarioles. The number of ovarioles per female varies among *Drosophila* species, but little is known about the genes and evolutionary dynamics that may shape interspecies changes in ovariole numbers. Here, used *a priori* experimental and transcriptome data from *D. melanogaster* to identify genes involved in ovariole formation and functions, and studied their molecular evolution among its closely related species within the *melanogaster* subgroup. Using a multi-layered analysis consisting of protein sequence divergence (dN/dS), adaptive evolution, expression breadth, and phylogenetic regressions, we identified 42 genes whose molecular evolution patterns were well linked to ovariole numbers divergence. Further, gene protein sequence divergence was often predictive of species ovariole numbers.

## Introduction

Ovarian development is a process that is poised to play key roles in organismal evolutionary biology, as the female gonads form and house the oocytes and/or eggs that are central to fertility and reproductive success of a species, and thus affect their fitness (Miller, et al. 2014; Macagno, et al. 2015). In insects, the most well-studied model with respect to ovarian development and genetics is the fruit fly *Drosophila melanogaster* (Dansereau and Lasko 2008; Eliazer and Buszczak 2011; Li, et al. 2014; Slaidina, et al. 2020; Lebo and McCall 2021). The mature ovary in *D. melanogaster*, as in other species of insects, is comprised of tubular egg-producing structures known as ovarioles (King, et al. 1968; Dansereau and Lasko 2008; Lebo and McCall 2021), which are a central factor shaping organismal reproductive output (Montague, et al. 1981; Starmer, et al. 2003; Church, et al. 2021). The number of ovarioles contained in the ovaries is highly variable within the genus *Drosophila* (Kambysellis and Heed 1971; Hodin and Riddiford 2000; Starmer, et al. 2003; Markow, et al. 2009; Sarikaya, et al. 2019; Church, et al. 2021). As an example, within the *melanogaster* subgroup, *D. melanogaster* has typically about 19 ovarioles per ovary, while its closely related sister species *D. sechellia* has only about 8 to 9 ovarioles per ovary (Hodin and Riddiford 2000). A broad range of ovariole numbers has been observed across the family *Drosophilidae*, from one to more than 50 per ovary across the genus *Drosophila* (Sarikaya, et al. 2019; Church, et al. 2021). At present, however, we know little about the genetic basis of the evolution of ovariole number within insects (Hodin and Riddiford 2000; Markow, et al. 2009; Sarikaya, et al. 2019).

A central factor that may underlie the rapid interspecies transitions in ovariole numbers in *Drosophila* is the evolvability of ovariole-related protein-coding genes, that is, the propensity of the proteins encoded by these genes to diverge and/or undergo adaptive sequence changes (Wagner and Zhang 2011; Cutter and Bundus 2020). Functional amino acid changes in protein-coding DNA and associated selection pressures (measured as nonsynonymous to synonymous changes, or dN/dS (Yang 1997; Bielawski and Yang 2005; Cutter and Bundus 2020)) can play a significant role in shaping interspecies divergence of developmental processes and other key phenotypes (Hoekstra and Coyne 2007). For instance, dN/dS of specific genes or sets of genes has been correlated with the divergence of sperm length in *Drosophila* (Chebbo, et al. 2021), sperm head size (Luke, et al. 2014) and testis size (Ramm, et al. 2008) in rodents, plumage color in toucans (Corso, et al. 2016), and brain mass in primates (Montgomery, et al. 2011), as well as other species traits (Swanson and Vacquier 2002; Hoekstra and Coyne 2007; Clark, et al. 2009; Cutter and Bundus 2020). Several lines of evidence indicate that ovariole number may also be a phenotype whose interspecies evolution in *Drosophila* is shaped by gene protein sequence changes and associated selection pressures (dN/dS, (Yang and Nielsen 2002; Bielawski and Yang 2005; Yang 2007)). Specifically, ovariole number is highly heritable and polygenic (Coyne, et al. 1991; Wayne and McIntyre 2002; Bergland, et al. 2008; Green and Extavour 2012; Sarikaya and Extavour 2015; Lobell, et al. 2017; Kumar, et al. 2020), and thus genetic mechanisms exist wherein changes in ovariole-related gene protein products could lead to interspecies differences in ovariole numbers. Further, in *Drosophila*, sexual (positive) selection pressures have been commonly observed and mating behaviors are variable among taxa (Kaneshiro and Boake 1987; Singh, et al. 2002; Singh and Singh 2014; Lupold, et al. 2016; Wigby, et al. 2020). These factors have been linked to accelerated interspecies protein sequence evolution in reproduction-related gene proteins and reproductive characteristics (Markow 2002; Swanson, et al. 2004; Jagadeeshan and Singh 2005; Haerty, et al. 2007; Kang, et al. 2016), that may potentially include ovariole numbers. Natural adaptive selection may also influence ovariole number evolution in *Drosophila*. For example, ovariole numbers and/or functions among species have been correlated with local environmental conditions and with oviposition and larval substrates in the *melanogaster* subgroup, as well as in the Hawaiian *Drosophila* (Kambysellis and Heed 1971; Kambysellis, et al. 1995; Sarikaya, et al. 2019). Finally, species-specific ovariole number may also be partly influenced by neutral protein sequence changes via random genetic drift (Kimura 1989; Kambysellis, et al. 1995). For these reasons, we sought to investigate whether evolutionary pressures on changes in proteins (dN/dS) involved in ovariole formation and function, especially in those genes that exhibit signs of evolvability and adaptive evolution, could underlie or even predict interspecies divergence in ovariole number, as is the case for certain other fitness-related phenotypes in animals (Montgomery, et al. 2011; Wagner and Zhang 2011; Luke, et al. 2014; Corso, et al. 2016; Chebbo, et al. 2021).

The most crucial developmental period that determines ovariole number in *D. melanogaster* is the larval stage (fig. 1) (King, et al. 1968; Godt and Laski 1995; Hodin and Riddiford 2000; Sarikaya, et al. 2012; Sarikaya and Extavour 2015; Slaidina, et al. 2020). Somatic gonad precursors specified during embryogenesis give rise to many different somatic ovarian cell types in the larval stage, and the numbers and behaviours of these somatic cells largely determine final ovariole number (Extavour and Akam 2003; Clark, et al. 2007; Dansereau and Lasko 2008). Specifically, the number of terminal filaments (TFs; fig. 1A), which are stacks of flattened intercalated terminal filament cells in the anterior ovary at the late third larval instar stage (LL3), determines adult ovariole number (King, et al. 1968; Godt and Laski 1995; Dansereau and Lasko 2008; Sarikaya, et al. 2012; Sarikaya and Extavour 2015). Each TF is the starting point for formation of a single ovariole (Sahut-Barnola, et al. 1996; Sarikaya, et al. 2012), which contains an anterior germarium housing germ line stem cells, and egg chambers that form the oocytes in an anterior to posterior pattern of oocyte maturation (Sahut-Barnola, et al. 1996; Eliazer and Buszczak 2011; Sarikaya, et al. 2012; Lebo and McCall 2021; Slaidina, et al. 2021). Single-celled RNA sequencing (sc-RNA seq) data (Slaidina, Banisch et al. 2020) suggest that LL3 TFs have anterior (TFa) and posterior (TFp) subgroups with distinct transcriptional profiles (fig. 1A). Another key somatic cell type are the sheath cells, also located at the anterior of the LL3 ovary (fig. 1A), and sub-categorized based on sc-RNA seq into anterior sheath cells (SHa) and migrating sheath cells (SHm). The latter cells migrate in an anterior to posterior direction between the TFs, depositing basement membrane that partitions the remaining cells of the ovary (germ cells and posterior somatic cells) into the developing ovarioles (King, et al. 1968; King 1970; Slaidina, et al. 2020). Additional somatic cells in the LL3 ovary include intermingled cells, which are interspersed between the germ cells and are involved in their proliferation (Gilboa and Lehmann 2006), cap cells, which form the adult germ line stem cell niche (Song, et al. 2002), follicle stem cell precursors, which give rise to adult follicle stem cells (Slaidina, et al. 2020; Slaidina, et al. 2021), and swarm cells, whose precise functions largely remain to be ascertained (Slaidina, et al. 2020) (fig. 1A). In this regard, understanding the interspecies evolution of ovariole number in *Drosophila* requires consideration of the genes and proteins regulating cell behaviour in the larval ovary, and particularly the behaviours of the TF and SH cells, which are instrumental to determining ovariole numbers in *D. melanogaster*.

**Figure 1.**
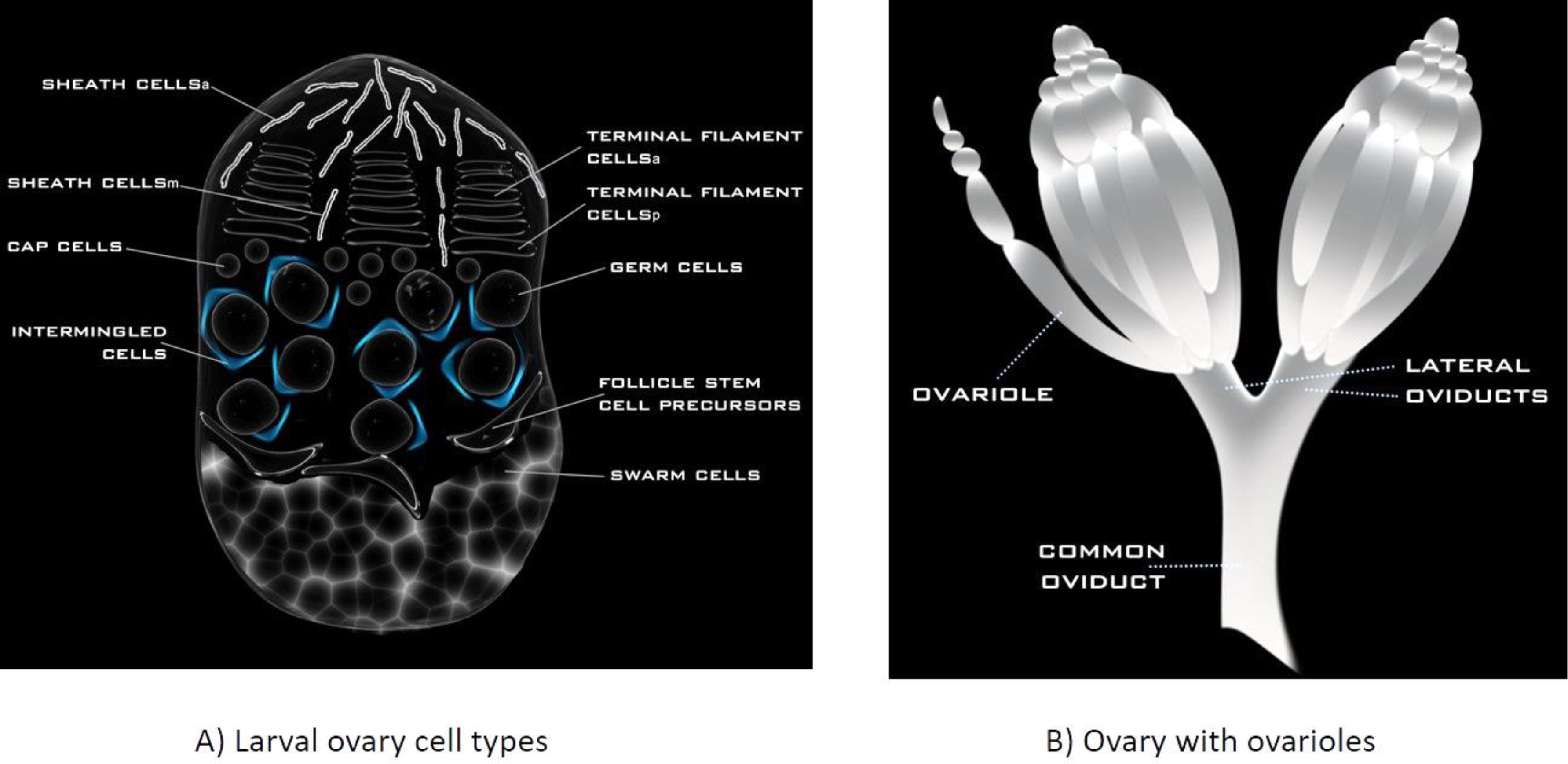
A schematic diagram of A) the late third instar larval ovary with its germ cells and various somatic cell types; and B) an external view of an adult ovary showing the ovarioles in each of the two ovaries that converge to the common oviduct in *D. melanogaster*. The relative cell positioning of cells in panel A is as denoted by Slaidina, et al. (2020). For orientation, anterior is up in both panels.

Until recently, research on the relationships between divergence in gene sequences and ovariole numbers in *Drosophila* was challenged by the lack of data on the identity of protein-coding genes expressed in somatic cells of the larval ovary that regulate ovariole number (Sarikaya, et al. 2012; Sarikaya and Extavour 2015). Recently available large-scale functional genetic and cell type-specific expression data from *D. melanogaster,* however, now provide a means to systematically identify genes linked to ovariole numbers, and in turn, assess their molecular evolution across species. A large-scale RNAi screen of 463 signalling genes from 14 conserved animal signalling pathways revealed that TF-mediated ovariole number determination is regulated by all conserved animal signalling pathways (Kumar, et al. 2020). Another study using bulk-RNA seq expression data from FACS-separated germ cells and somatic cells revealed additional genes differentially expressed throughout TF formation, suggesting their potential involvement in ovariole number regulation (Tarikere, et al. 2022). In addition to those studies, a recent sc-RNA seq study yielded unique transcriptional profiles for all of the known cell types in the *D. melanogaster* LL3 ovaries (fig. 1), providing a novel resource to identify and study the evolution of genes transcribed in TF and SH cells, the two crucial cell types in determining ovariole number (Slaidina, et al. 2020).

Collectively these datasets provide valuable empirical data from which to *a priori* identify sets of genes that regulate ovariole numbers or functions in *Drosophila*, and in turn, to evaluate which of these genes exhibit elevated or otherwise unusual rates of interspecies protein sequence evolution, including adaptive evolution, suggesting them as candidates for driving interspecies divergence of ovariole numbers in *Drosophila*. For example, by assessing dN/dS, we may ask whether ovariole-related gene protein sequences typically have been under strict purifying selection, which could mean that phenotypes regulated by these genes are likely to show high pleiotropy and low evolvability, and to have minimal potential to diverge neutrally or adaptively (Fisher 1930; Otto 2004; Wagner and Zhang 2011; Cutter and Bundus 2020; Munds, et al. 2021). If, in contrast, some ovariole-related genes have been subjected to relaxed selection and/or have commonly experienced adaptive changes, we might expect high phenotypic evolvability and adaptability (Otto 2004; Larracuente, et al. 2008; Clark, et al. 2009; Mank and Ellegren 2009; Montgomery, et al. 2011; Luke, et al. 2014; Corso, et al. 2016; Chebbo, et al. 2021). In this regard, the study of the evolution of protein-coding genes (from dN/dS) that are pre-screened for likely roles in ovariole numbers and/or functions by studies like the ones described above (Kumar, et al. 2020; Slaidina, et al. 2020; Tarikere, et al. 2022) provides a novel pathway to advance our understanding of the genetic factors and evolutionary forces that shape rapid interspecies divergence in ovariole numbers.

In the present study, we rigorously assess the molecular evolutionary patterns of genes that regulate ovariole numbers and/or functions, that were identified *a priori* based on one or both of functional genetic evidence (Kumar, et al. 2020) or transcriptional activity (Slaidina, et al. 2020; Tarikere, et al. 2022). We focus on the molecular evolution of ovariole-related genes within five species of the *melanogaster* subgroup of *Drosophila*, that is a closely related species clade that includes *D. melanogaster*, diverged from a common ancestor about 13 Mya (Tamura, et al. 2004), and exhibits substantial interspecies variation in ovariole numbers (Hodin and Riddiford 2000; Starmer, et al. 2003; Markow, et al. 2009). From our assessments, we identify 42 genes that are high confidence candidates for contributing to the genetic basis of interspecies divergence in ovariole numbers. We hypothesize that evolved changes in these genes are apt to underlie ovariole number divergence among taxa given that they exhibit an ovariole-related function (Kumar, et al. 2020; Slaidina, et al. 2020; Tarikere, et al. 2022), have a propensity to diverge in protein sequence, or high evolvability, show a high frequency of adaptive sequence evolution events in branches of the phylogeny, and are associated with low pleiotropy (Yanai, et al. 2005). Further, phylogenetic regressions show gene dN/dS has predictive associations to ovariole numbers. Collectively, our findings provide a genetic framework to explain the rapid interspecies divergence of ovariole numbers in *Drosophila*, which we propose is largely mediated by selection pressures shaping the evolution of functional protein sequences, and thus ovariole numbers.

## Results

### The Clade Under Study, the *melanogaster* subgroup

For our study, we focused on a multi-layered analysis of the molecular evolution of ovariole-related genes across five species from the *melanogaste*r subgroup of *Drosophila*: *D. simulans* (Dsim), *D. sechellia* (Dsec)*, D. melanogaster* (Dmel), *D. yakuba* (Dyak), and *D. erecta* (Dere) (fig. 2; *D. ananassae* of the *melanogaster* group was used as an outgroup for phylogeny construction, see “*Drosophila* Phylogeny” section; the abbreviated names were used in tables and figures). Using this closely-related species clade, we hypothesize that if genes with demonstrated roles in regulating ovariole numbers or formation are involved in the interspecies divergence of ovariole numbers, then they will exhibit relatively rapid evolution (dN/dS) as compared to the genome, as well as interspecies variation in dN/dS, signs of positive selection, and low pleiotropy (as inferred by high *tau* across tissues, table S1). We further hypothesize that, if evolutionary variation in these genes contributes to the genetic basis of evolved shifts in ovariole number, that dN/dS values for these genes may predict species ovariole numbers.

**Figure 2.**
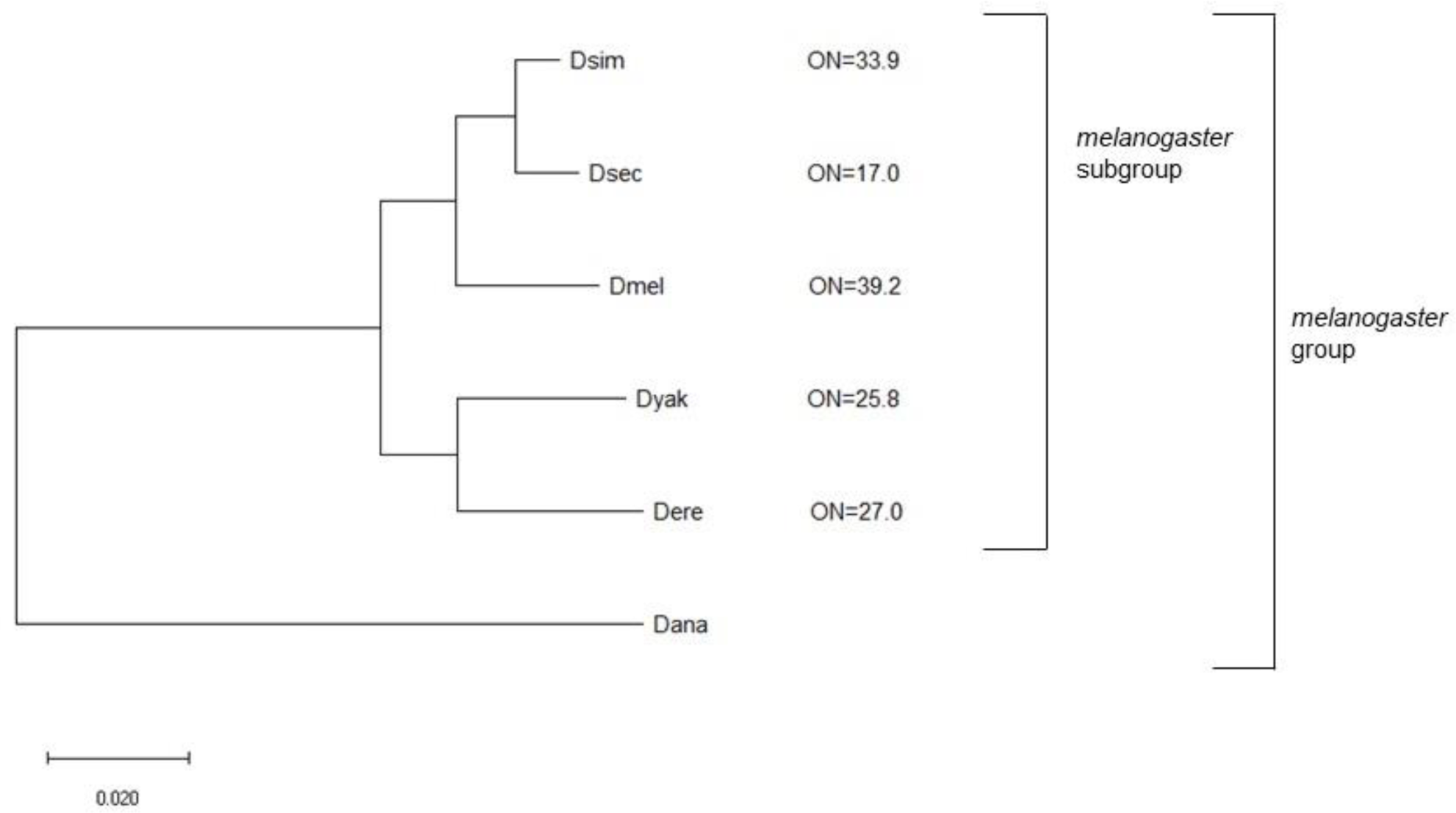
The phylogeny showing the five-species *melanogaster* subgroup under study that was based on a Maximum Likelihood tree generated in MEGA v. 11 (Tamura, et al. 2021) and DNA sequence data from DrosoPhyla (Finet, et al. 2021). The five species of the *melanogaster* subgroup are shown. The relatively distantly related *D. ananassae* (Dana) was used as an outgroup for tree construction. Ovariole numbers (ON) are shown and are for two ovaries per female and are from the following sources: *D. melanogaster* (Dmel), *D. sechellia* (Dsec), and *D. yakuba* (Dyak) (Hodin and Riddiford 2000), *D. simulans* (Dsim) (averaged, (Hodin and Riddiford 2000; Starmer, et al. 2003) and *D. erecta* (Dere) (Markow, et al. 2009) (see respective articles for variation). All nodes had 100/100 bootstrap support.

The five-species clade of *melanogaster* subgroup had the following advantages for our study: (1) all species within the clade are very closely related to *D. melanogaster* (Tamura, et al. 2004; Obbard, et al. 2012), the species for which experimental and transcriptome data on genes associated with ovariole numbers or functions are available (Kumar, et al. 2020; Slaidina, et al. 2020; Tarikere, et al. 2022), and thus we hypothesize are likely to share similarities in the genetic pathways affecting ovariole numbers, more so than we would be expect for distantly related species; (2) the clade exhibits substantial variation in ovariole numbers among species, typically about 39.2 (per female) for *D. melanogaster* and 17.0 for *D. sechellia* and intermediate values for *D. simulans* (33.9), *D. yakuba* (25.8) and *D. erecta* (27.0) (fig. 2; see values and variability (Hodin and Riddiford 2000; Starmer, et al. 2003; Markow, et al. 2009), and includes both some species with similar ovariole numbers and others that markedly differ; (3) the phylogeny is highly resolved (fig. 2 (Cutter 2008; Obbard, et al. 2012), unlike some other *Drosophila* clades and branches (Finet, et al. 2021)), and the five species are very closely related to each other (Tamura, et al. 2004; Cutter 2008). We made this choice to minimize biological differences other than ovariole numbers among taxa, and to facilitate the detection of putative cause-effect relationships (here, dN/dS and ovariole number (Felsenstein 1985; Bromham, et al. 1996; Whittle and Johnston 2003; Thomas, et al. 2010; Symonds and Blomberg 2014). The close relatedness of species is more conducive to accurate alignments, and retains a larger set of orthologous genes, including rapidly evolving genes, for study, than when studying more divergent species, which often skews toward the identification of fewer and more slowly evolving orthologous gene sets (*cf.* (Stanley and Kulathinal 2016; Bubnell, et al. 2022)), and may exclude some rapidly evolving genes of interest.; (4) each species has a whole genome sequence available (Gramates, et al. 2022) and; (5) the dN and dS values among the species in this subgroup have substantially diverged, yet are also unsaturated in the frequency of substitutions, and thus are within the ideal range for dN/dS analysis (Castillo-Davis, et al. 2004; Larracuente, et al. 2008; Treangen and Rocha 2011) (for example, from M0 dN/dS values (that is, the single clade-wide measure of dN/dS, (Stanley and Kulathinal 2016)), we found that the 95^th^ percentile for M0 dN=0.235 and M0 dS=0.791 for the 9,232 genes that had orthologs in all five species and M0 values). In sum, this closely related taxonomic group has multiple benefits for the study of the evolution of ovariole-related genes.

### Identification of Rapidly-Evolving Ovariole-Related Genes for Follow-up Study

To identify genes associated with ovariole numbers or functions for study, we focused on three recently available datasets from *D. melanogaster*. The first gene set we designate as the SIGNALC dataset, defined here as the signalling and connector genes (connectors identified by protein interaction networks) that were identified as affecting ovariole or egg numbers in a *hpo[RNAi]* and/or a *hpo[+]* background (Kumar, et al. 2020). Among 463 signalling genes and additional connector genes studied, the authors reported 67 genes that affected ovariole number in a *hpo[RNAi]* background (named therein *hpo[RNAi]* Ovariole Number), 59 and 49 genes that affected egg laying in a *hpo[RNAi]* background (*hpo[RNAi]* Egg Laying) and a wild type (*wt*) background (Egg Laying *[wt]*) respectively, and 17 connector genes that altered ovariole or egg laying phenotypes (and passed screening of z>1; note that genes may belong to more than one category) (Kumar, et al. 2020). The second is the BULKSG dataset, based on bulk-RNA seq data obtained from pooled larval ovarian somatic cells or germ cells from the early (72 hours after egg laying = 72h AEL), mid (96h AEL) and late (120h AEL) TF developmental stages (Tarikere, et al. 2022) and identified differentially expressed genes (P-values were from DeSeq2 (Love, et al. 2014)). The third is the SINGLEC dataset (Slaidina, et al. 2020), a sc-RNA seq dataset that provided expression data for each of the cell types of the *D. melanogaster* LL3 larval ovary (fig. 1) (Slaidina, et al. 2020). The SINGLEC study assessed average standardized expression to identify differentially expressed genes among cell types (P-values from Seurat v.2; some genes were upregulated in more than one cell type using the criteria therein (Slaidina, et al. 2020)).

The SIGNALC, BULKSG and SINGLEC gene sets were screened for further study using their clade-wide M0 dN/dS values (Yang 2007), that reflects the rate of protein divergence and the potential types of selective pressures that may have affected a gene (Yang 1997, 2007). Values of dN/dS <1 suggest a history of purifying selection on protein sequences, =1 infer neutral evolution, and >1 suggest a history of positive selection (Yang 1997, 2007); however, even when dN/dS <1 across an entire gene (Yang 2007), elevated dN/dS values in one gene relative to another suggest an enhanced degree of positive selection and/or neutral evolution (Yang 1998, 2007; Buschiazzo, et al. 2012; Ho and Smith 2016; Mitterboeck, et al. 2017; Whittle, et al. 2021). We identified those ovariole-related genes with an M0 dN/dS value at least 1.5-fold (SIGNALC; lower cut-off due to conserved nature of signalling genes, see Materials and Methods) or 2-fold (BULKSG and SINGLEC) higher than the genome-wide medians, and we then conducted a thorough follow-up analysis that included the M1 free-ratio species branch dN/dS (e.g., (Dorus, et al. 2004; Nadeau, et al. 2007; Clark, et al. 2009; Wlasiuk and Nachman 2010; Mensch, et al. 2013; Borges, et al. 2019; Kong, et al. 2019; LaBella, et al. 2021)), branch-site tests of positive selection (Zhang, et al. 2005; Yang 2007), *tau* (Yanai, et al. 2005) and phylogenetic regressions (R-Core-Team 2022) (see Materials and Methods).

### Some Signalling Pathway Genes that Regulate Ovariole Number Have Evolved Rapidly

We report that for the ovariole-related SIGNALC gene set (Kumar, et al. 2020), that included signalling genes that affected ovariole number and/or egg laying, many genes exhibited very low M0 dN/dS (MWU-tests had P<0.05 versus the genome-wide values; fig. 3A). This suggests a history of strong purifying selection on these highly conserved signalling genes, which may be partly due to their high pleiotropy, given that all of these signalling pathways play multiple roles in development and homeostasis (Kumar, et al. 2020). Consistent with this hypothesis, the *tau* values for these genes were statistically significantly lower than the genome-wide values (fig. 3B; MWU-tests P<0.05), suggesting that broad expression breadth may have acted to slow molecular evolution (Otto 2004; Kim, et al. 2007; Cui, et al. 2009; Mank and Ellegren 2009; Meisel 2011; Assis, et al. 2012; Masalia, et al. 2017; Whittle, et al. 2021).

**Figure. 3.**
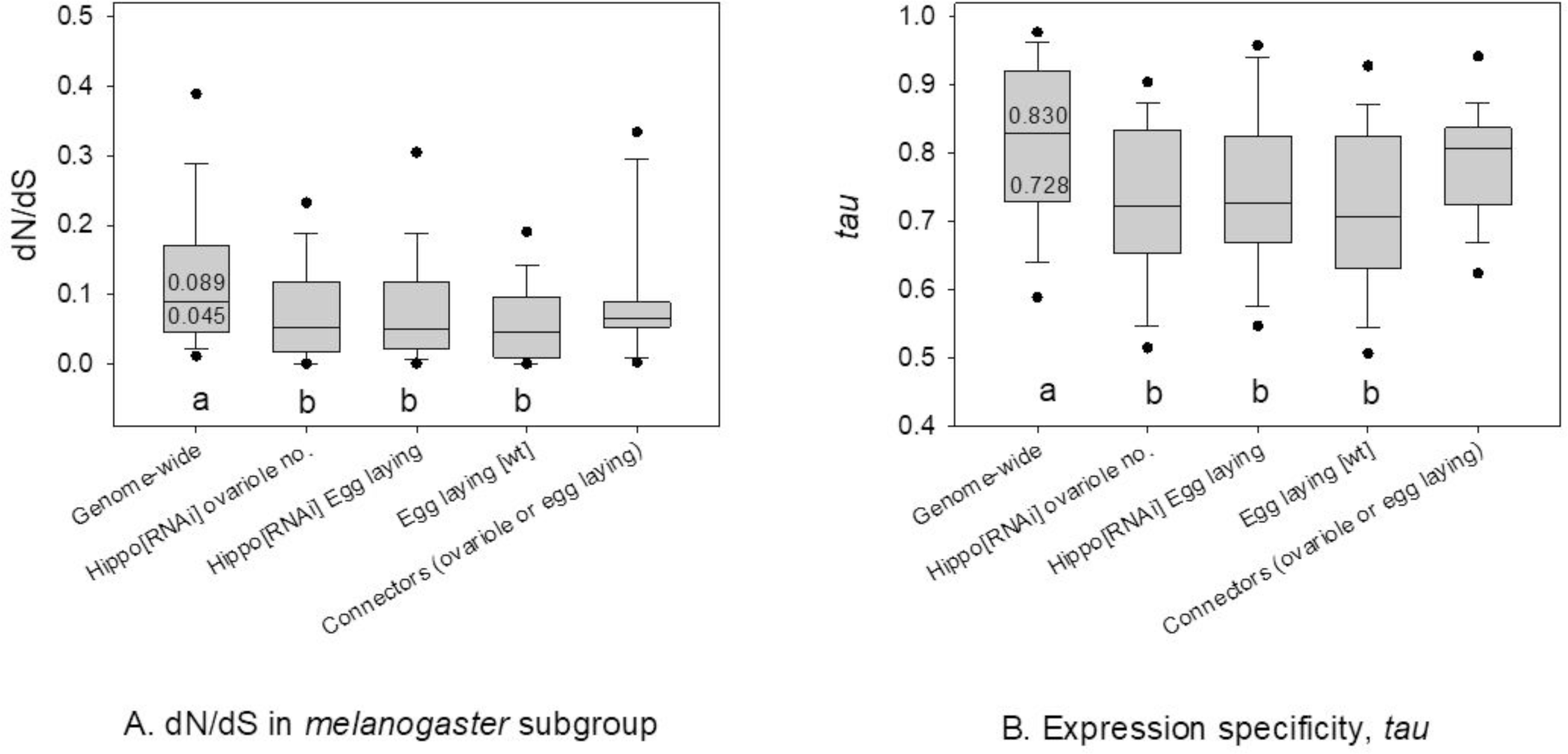
Box plots of A) M0 dN/dS of genes with five-species orthologs in the *melanogaster* subgroup for each of four groups of signalling/connector genes that affected ovariole/egg numbers using RNAi in *D. melanogaster* (Kumar, et al. 2020) and for the genome-wide values; and B) *tau* for all genes in each of the four groups of ovariole number/egg laying affecting genes and the genome-wide values. Different letters (a. b) below bars indicate a statistically significant difference (MWU-tests P<0.05) between the genome-wide values and each group of genes. The median and 25^th^ percentiles are shown for dN/dS and *tau* as reference points for the genome-wide values (that is, across all 9,232 genes with known dN/dS and five-species orthologs).

Importantly however, our main goal herein was to identify whether any ovariole-related SIGNALC genes evolved unusually rapidly, and showed signs of evolvability that could underlie interspecies ovariole number divergence. As shown in table S2, we indeed identified 27 SIGNALC genes that had elevated M0 dN/dS in at least one of the studied *Drosophila* taxon groups (≥1.5 fold higher than the genome-wide median; table 1, table S2, see also Supplementary Text File S1 Results, and table 1 Notes for *Paris*). The signalling pathways and example functions of each of these genes are provided in table S3: we found they were preferentially involved in developmental and cytoskeletal roles. Thus, it is apparent that while most of the ovariole number-related signaling genes evolved under strong purifying selection (fig.3A), a subset of them exhibited a high rate of amino acid sequence changes, well above the genome-wide median, in the *melanogaster* subgroup of *Drosophila*. This pattern shares similarities to the previous finding that while most *D. melanogaster* developmental genes expressed at the phylotypic stage of embryogenesis evolved under strong purifying selection (low dN/dS), a subset of genes expressed at this stage exhibited a history of positive selection (Mensch, et al. 2013).

**Table 1.**
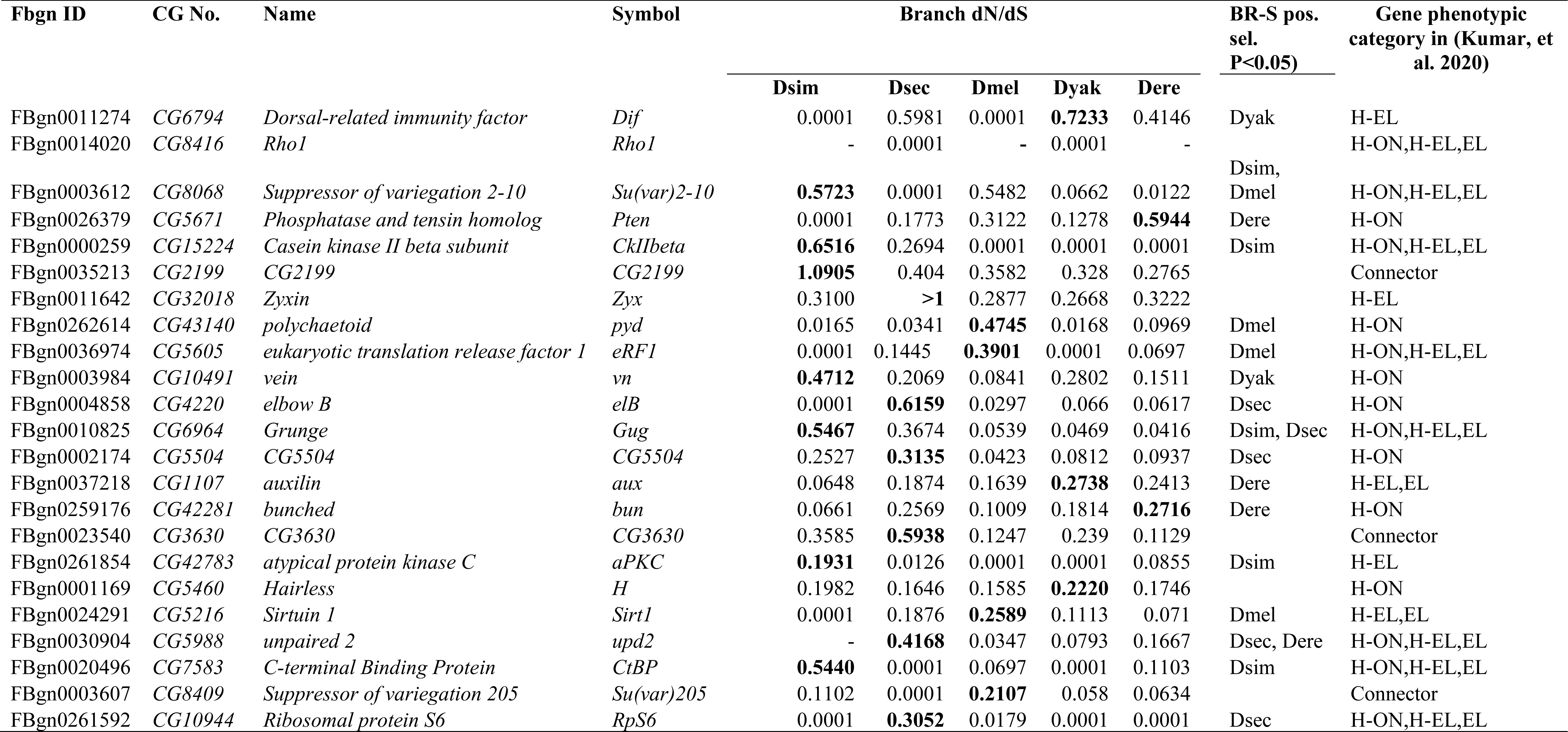
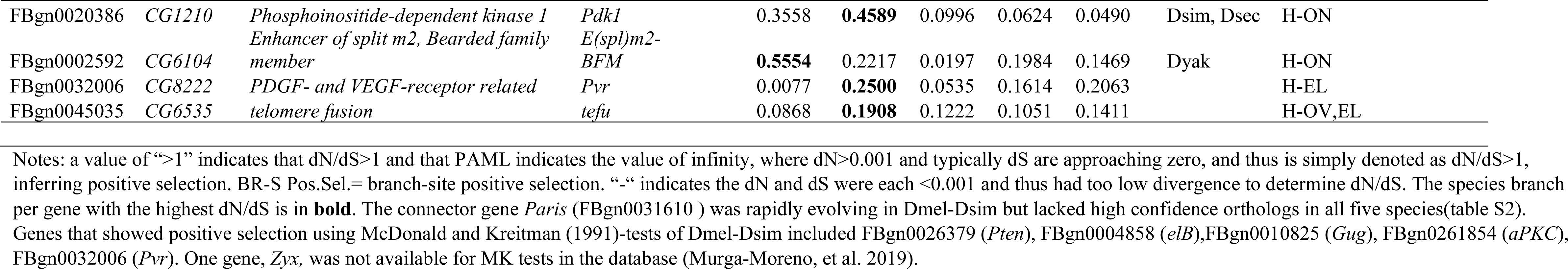
The gene-wide dN/dS per species branch values for each of the 27 signalling or connector genes (determined to be evolving rapidly in table S2) in the five terminal species branches in the *melanogaster* subgroup of *Drosophila*. Branch-site positive selection (BR-S pos. sel,) analysis and cases with P<0.05 are shown by species name (Dsim = *D. simulans*, Dsec = *D. sechellia*, Dmel = *D. melanogaster*, Dyak = *D. yakuba* and Dere = *D. erecta*). The ovariole number/egg laying phenotypic categories defined in the RNAi experiments from (Kumar, et al. 2020) are shown here as: H-ON for *hpo[RNAi]* Ovariole Number, H-EL for *hpo[RNAi]* Egg Laying, and EL for the Egg Laying *[wt]*, and genes designated in that study as “connector genes” with observed phenotypes (on ovariole number or egg laying) are also shown.

### Rapid and adaptive evolution of specific signalling genes coincides with ovariole number evolution

To examine potential lineage-specific patterns of molecular evolution and pleiotropy of the 27 rapidly evolving ovariole-related genes, we assessed dN/dS per species branch (table 1), branch-site positive selection (table 1), and *tau* (table S3). We found that these 27 genes showed marked differences in dN/dS values per gene among the five-species terminal branches in the *melanogaster* subgroup (the distribution of dN/dS for all genome-wide genes per species branch are shown in box plots in fig. S1). In addition, we observed branch-site positive selection in at least one species branch for 19 of the 27 genes (table 1), which is consistent with potential high adaptability of these genes. Of particular note is the *D. sechellia* branch, as this species evolved a very low ovariole number (17 ovarioles per female, fig. 2), only half that of its most closely related sister species *D. simulans* (33.9 ovarioles per female, fig. 2), since diverging from their recent common ancestor. Among the five species terminal branches, the *D. sechellia* terminal branch had the highest dN/dS values for nine genes (table 1), namely *Zyx*, *elB*, *CG5504, CG3630*, *upd2*, *RpS6*, *Pdk1,Pyr* and *tefu*, with values ranging from 0.191 to >1. Further, five of these genes exhibited branch-site positive selection on amino acids in the *D. sechellia* branch (*elB, CG5504, unp2*, *RpS6, Pdk1*, branch-site P <0.05 for all genes (Zhang, et al. 2005; Yang 2007)), explicitly showing a propensity for adaptive evolution in this species branch. In total, six of the 27 genes 22.2%) exhibited branch-site positive selection in the *D. sechellia* terminal branch. This was nearly double the genome-wide frequency for this species, which was 12.0% of 9,232 genes (one tailed Chi-square P=0.05). Thus the *D. sechellia* lineage, with the lowest ovariole numbers (fig. 2), has a dynamic molecular evolutionary history of ovariole number-regulating genes, consisting of rapid gene-wide evolution (dN/dS), combined with a pervasiveness of positive selection events on such genes in that species branch.

In *D. sechellia*’s sister species *D. simulans* (fig. 2), eight genes had the highest dN/dS values in the *D. simulans* terminal branch (table 1), five of which also exhibited statistically significant branch-site positive selection (*Su(var)2-10*, *CkIIbeta*, *Gug*, *aPKC*, *CtBP*, P<0.05, table 1). In total, six of the studied 27 SIGNALC genes (22.2%) presented branch-site positive selection in the *D. simulans* branch, which was more than four-fold higher than the genome-wide frequency for the species (5.4%, Chi-square P<0.05). In turn, four of 27 genes had the highest dN/dS in the *D. melanogaster* branch, and four genes had branch-site positive selection in *D. melanogaster* (14.8%), which was more than triple its genome-wide frequency (4.1%; Chi-square P<0.05). *D. yakuba* and *D. erecta* had the highest dN/dS for three and two genes respectively, and had branch-site positive selection in three and four genes (table 1). In sum, for the *melanogaster* subgroup, all five species terminal branches showed signs of having the highest dN/dS values for at least two (*D. erecta*) and up to nine (*D. sechellia*) genes, exhibited signals of branch-site positive selection, and had particularly high rates of protein sequence divergence.

The patterns in table 1 support the hypothesis that protein sequence changes, including adaptive changes, in these ovariole-related genes may underlie the genetic basis for the marked divergence in interspecies ovariole numbers (fig. 2). For many of these genes, their known molecular and genetic mechanisms of action in tissue morphogenesis make them prime candidates for future analyses of how their diverged functions between species may have contributed to species-specific ovariole number evolution. For example, *Zyx* (*Zyxin*) is an actin cytoskeleton regulator and a signal transducer in the Hippo pathway, and mis-regulation of either actin cytoskeleton function (Li, et al. 2003) or Hippo signaling function (Sarikaya and Extavour 2015; Kumar, et al. 2020) during ovariole morphogenesis can alter ovariole number. We provide further discussion of some of these ovariole-related signalling genes in table 1 within the Supplementary Text File S1.

### Multiple Genes Highly Upregulated in Larval Ovary Somatic Cells Have Evolved Rapidly

We identified genes whose high differential expression in the *D. melanogaster* larval ovary suggested a role in ovariole number regulation using the BULKSG RNA-seq datasets using pooled larval ovarian somatic versus pooled germ cells from different stages of TF formation (Tarikere, et al. 2022). First, we asked whether the 27 rapidly evolving ovariole-related SIGNALC genes in table 1 exhibited statistically significant differential expression between somatic and germ cells during TF formation (therein P<0.01, (Love, et al. 2014; Tarikere, et al. 2022)). Remarkably, as shown in table S4, we report that 25 of the 27 rapidly evolving SIGNALC ovariole-related genes showed up- or downregulation in the soma (versus germ cells; each cell type pooled across stages), or among the three different TF formation stages. Thus, this affirms that the SIGNALC genes in table 1 that experimentally affected ovariole numbers or functions using RNAi (Kumar, et al. 2020), and that showed signals of enhanced evolvability herein (table 1, table S3), also exhibited differential expression in the larval somatic ovary cells, based on an independent approach of bulk RNA-seq (Tarikere, et al. 2022). These two lines of evidence suggest that these genes are apt to have contributed towards the genetic basis of evolved ovariole number divergence.

### Rapidly Evolving Genes are Highly Transcribed in the Larval Ovary Somatic Cells

We aimed to further identify any rapidly evolving genes that were highly differentially expressed in the larval ovarian soma during TF formation, and thus potentially involved in the evolution of ovariole number, using the BULKSG datasets. For this, we identified genes that were upregulated in the soma versus the germ cells, ranked them by log_2_fold upregulation, and in that subset, screened for genes that were rapidly evolving in the *melanogaster* subgroup as compared to the genome-wide values (see Methods, M0 dN/dS>0.20). The top ten genes matching these criteria are shown in table 2, with the highest log_2_fold values ranging from 5.1 to 10.0, and includes the branch dN/dS, branch-site positive selection tests for each species of the *melanogaster* subgroup and *tau* values (see Supplementary Text File S1 for analysis of genes highly upregulated in germ cells, table S5).

**Table 2.**
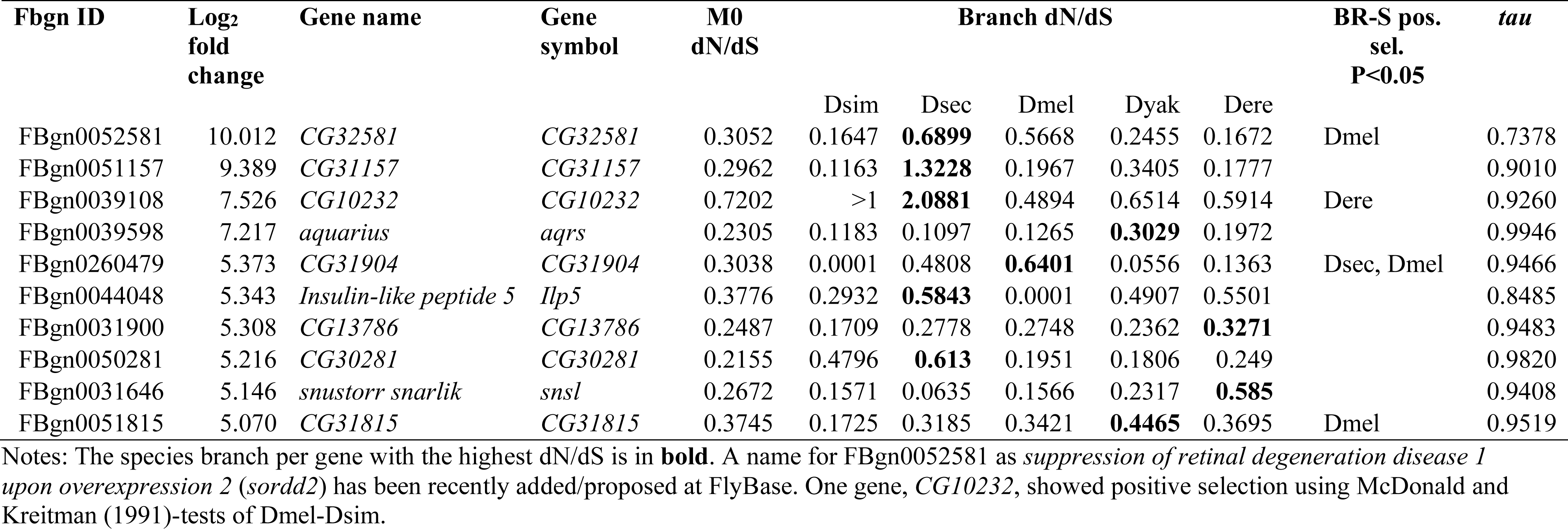
Genes that were highly upregulated in the larval ovary somatic cells relative to germ cells when pooled across three larval stages (Tarikere, et al. 2022) and that exhibited rapid protein sequence divergence in the *melanogaster* subgroup (M0 dN/dS>0.20). The dN/dS per species terminal branch, branch-site positive selection (P<0.05) and *tau* values are shown for each gene. The genes with the top 10 log_2_ fold change values matching these criteria are shown.

Remarkably, eight of the ten most highly upregulated and rapidly evolving somatic genes had extremely elevated *tau* values >0.90, and six had values above 0.94, indicating very narrow expression breadth (as compared to genome-wide values in fig. S2). This low pleiotropy may facilitate their rapid evolution, via neutral evolution, and/or by adaptive sequence evolution (Otto 2004; Larracuente, et al. 2008; Mank and Ellegren 2009). For the *D. sechellia* branch, five of the ten genes had the highest dN/dS in this species terminal branch, including *Ilp5 (Insulin-like peptide 5*, dN/dS=0.5843, discussed in Supplementary Text File S1) and four unnamed genes (CG identifiers only, *CG32581, CG31157*, *CG10232, CG30281*). Two of these, *CG31157* and *CG10232,* exhibited gene-wide positive selection with dN/dS values larger than 1, and the latter gene also had dN/dS >1 in *D. simulans* (table 2). Further, *CG31904* exhibited branch-site positive selection in *D. sechellia* (table 2). These patterns are consistent with a prevalent history of rapid protein evolution coupled with the ovariole number decline within the *D. sechellia* branch, as also observed for multiple SIGNALC genes (table 1). Further, three of the ten genes also showed branch-site positive selection in *D. melanogaster*, and one displayed this pattern in *D. erecta* (table 2), suggesting that many of these genes experienced a history of adaptive evolution across multiple lineages of the phylogeny.

### Terminal Filament Cells and Sheath Cells Express Rapidly Evolving Genes

The SINGLEC dataset was based on sc-RNA seq data generated from the late third instar *D. melanogaster* ovary (Slaidina, et al. 2020) and includes expression data for all the cell types shown in fig. 1 (the germ cells (GC) and eight somatic cell types, namely the cap cells (CC), follicle stem cell precursors (FSCP), intermingled cells (IC), anterior sheath cells (SHa), migrating sheath cells (SHm), swarm cells (SW), anterior terminal filament cells (TFa), and posterior terminal filament cells (TFp)). Using hierarchical clustering of average standardized gene expression per gene, across all genes (fig. S3), we found that the germ cells exhibited the most unique transcriptome of all studied cell types, and formed an outgroup to all somatic cells. Among the somatic cells, the two types of terminal filament (TF) cells, TFa and TFp, formed their own cluster, as did the two types of sheath (SH) cells, SHm and SHa; each of these clusters was separate from all other somatic cell types (fig. S3). The FSCP and SW cells had highly similar transcription profiles, as did the IC and CC cells. Thus, the TFs and SH cells had more distinctive transcriptomes than the other LL3 ovarian somatic cell types.

#### Rapidly evolving genes identified in both the BULKSG and SINGLEC datasets

To identify genes with roles in specific ovarian cell types that were putatively involved in interspecies ovariole number divergence, we first extracted those SINGLEC genes that were upregulated in one cell type relative to all others (P<0.05, analyzed in Seurat v. 2; genes could be upregulated in more than one cell type (Satija, et al. 2015; Slaidina, et al. 2020)), and that also had M0 dN/dS more than two fold above the genome-wide median (dN/dS>0.20) within the *melanogaster* subgroup. We then compared this SINGLEC gene set to the 30 most highly differentially expressed and rapidly evolving genes identified from the somatic larval ovary cells at three different stages of development for terminal filament formation (listed in table S6, extracted from BULKSG dataset) and determined whether any genes were upregulated in both datasets. We identified five genes that matched these criteria (table 3): *Drip*, *CG371*3, *MtnA*, *vkg*, and *Col4a1* (table 3). Among the nine somatic cell types, these genes were nearly exclusively upregulated in the TFs (TFa or TFp, or both) and/or the SHm cells. We note that *vkg* and *Col4a1* play roles in basement membrane formation (Yasothornsrikul, et al. 1997; Kiss, et al. 2019), and that SHm cells lay the membrane that separates the TFs for ovariole development (King 1970; Slaidina, et al. 2020). Given the crucial roles of these cell types in determining ovariole number (King, Aggarwal et al. 1968, Sarikaya and Extavour 2015), the rapid evolution of these five genes may partially underlie ovariole number divergence between species (King, et al. 1968; Sarikaya and Extavour 2015) in the *melanogaster* subgroup (table 3).

**Table 3.**
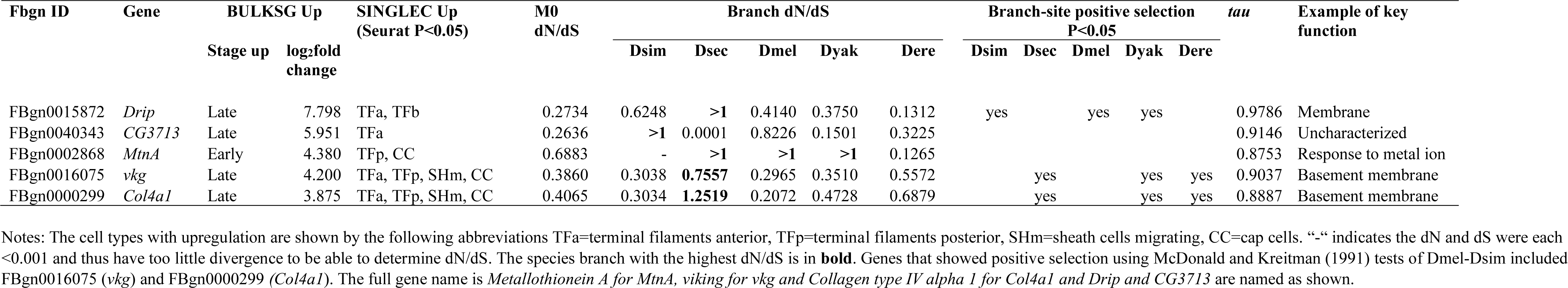
Genes with rapid divergence (M0 dN/dS>0.20) and that were highly upregulated at one stage of the larval ovary somatic cells (versus the others; three stages early, mid, late, among the top 30 most upregulated genes, table S5) in Dmel using BULKSG data (Tarikere, et al. 2022) and that also exhibited upregulation in at least one cell type (versus all others) using SINGLEC data among the nine studied LL3 ovary cell types (Slaidina, et al. 2020). Shown are the dN/dS per species branch, the presence of branch-site positive selection (P<0.05), the *tau* values and an example of key functionality as described in DAVID (Huang da, et al. 2009). “Stage up” indicates the larval ovary stage where the gene was upregulated (P<0.05). SINGLEC up indicates the cell type(s) with upregulation.

In terms of molecular evolution per terminal species branch, the five genes in table 3 exhibited a striking propensity for adaptive evolution. Four the five genes showed a gene-wide level of positive selection (terminal branch dN/dS values >1) in at least one species branch (table 3). Moreover, *Drip*, *vkg* and *Col4a1* each exhibited branch-site positive selection in three different species branches (P<0.05), suggesting a profound history of adaptive changes across multiple lineages. In addition, McDonald and Kreitman (1991) tests also showed positive selection for *vkg* and *Col4a1* (P<0.05, table 3 Notes). All five genes exhibited *tau* values above 0.875 with *Drip* having a value of 0.979, suggesting especially high expression specificity (see Materials and Methods, fig. S2), which may facilitate the observed adaptive evolution of the protein sequences (Otto 2004; Mank and Ellegren 2009; Whittle, et al. 2021). In sum, these five genes were identified from two distinct expression datasets (Slaidina, et al. 2020; Tarikere, et al. 2022), were upregulated in two of the most crucial cell types for ovariole number determination namely TFs and SH cells (table S6, table 3), and exhibited rapid protein changes, positive selection, and narrow expression breadth (table 3). Thus, multiple lines of evidence point towards these genes as having a central role in the interspecies divergence of ovariole number.

#### Genes upregulated in TF and SH cells frequently display branch-site positive selection

We assessed the frequency of genes that exhibited branch-site positive selection (P<0.05) per species terminal branch for the rapidly evolving genes that were upregulated in each of the nine cell types in the SINGLEC dataset (P<0.05). The results for *D. simulans*, *D. sechellia* and *D. melanogaster* (a very closely related species group with substantial differences in ovariole numbers (fig. 2)), are shown in fig. 4, and for all five species in fig. S4. The genes with the highest percent branch-site positive selection were those upregulated in the SH and TF cells (fig. 4; the TF and SH genes are listed in Table S7). Specifically, positive selection was most commonly observed for genes up-expressed in the SHm cells for the *D. sechellia* branch (45%), from the TFa (34.1%) and TFp (36.7%) cells in the *D. sechellia* branch, and for SHa cells in the *D. sechellia* (33.33%) and *D. simulans* (33.33%) branches (all values were statistically significantly higher than the genome-wide percentages of genes with branch-site positive selection per species, which were 5.4% for *D. simulans* and 12.0% for *D. sechellia*; Chi-square P<0.05, fig. 4). Thus, the most important somatic cell types for ovariole number determination (TF and SH cells) (King, et al. 1968; Godt and Laski 1995; Sarikaya, et al. 2012; Sarikaya and Extavour 2015; Slaidina, et al. 2020), are also those in which highly upregulated genes most commonly exhibited branch-site positive selection, particularly in *D. sechellia*.

**Figure 4.**
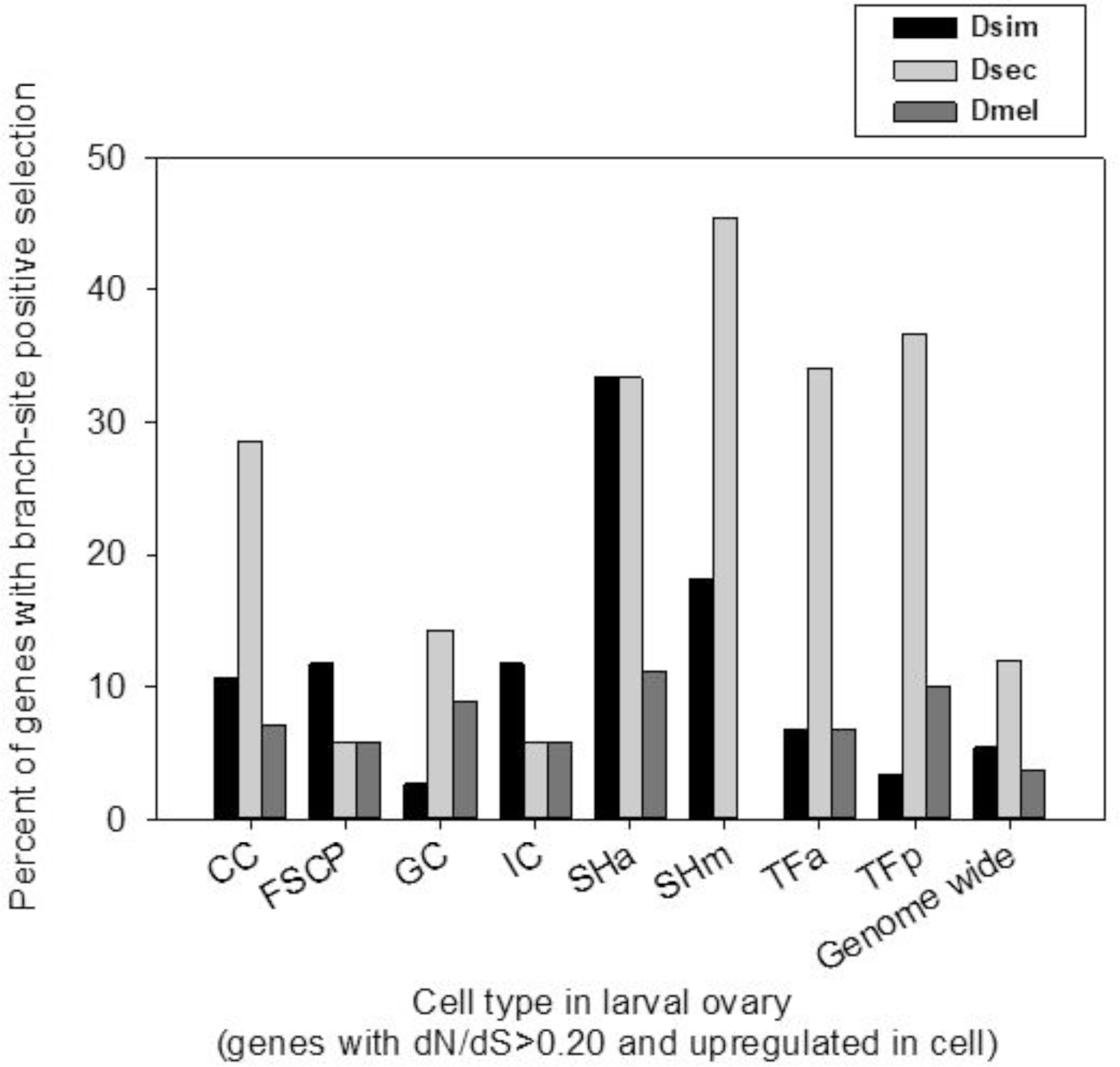
The percentage of the genes that were both upregulated in a particular cell type and rapidly evolving in the *melanogaster* subgroup (M0 dN/dS>0.20) that exhibited branch-site positive selection in the *D. simulans* (Dsim), *D. sechellia* (Dsec), and *D. melanogaster* (Dmel) branches (P<0.05). The number of genes per category were as follows: cap cells (CC: 28), follicle stem cell precursors (FSCP: 17), germ cells (GC: 112), intermingled cells (IC: 17), anterior sheath cells (SHa: 9), migrating sheath cells (SHm: 11), anterior terminal filament cells (TFa: 44), posterior terminal filament cells (TFp: 30). Swarm cells (SW) cells were excluded as too few genes were rapidly evolving for study (SW: 4). Note that a gene could be upregulated in more than one cell type. The genome-wide values are for all genes with five-species orthologs in the *melanogaster* subgroup.

The genes identified above as highly expressed in TF and SH cells, could also be highly expressed in additional cell types (Slaidina, et al. 2020). Indeed, on average we found that differentially expressed genes were upregulated in 1.9±0.02 cell types. Thus, for additional stringency we isolated the subset of rapidly evolving genes (with M0 dN/dS>0.20) that were upregulated in only one cell type. While most somatic cell types had very few genes matching this stringent criterion (N≤4 per cell type), by pooling the two types of SH cells (SHa and/or SHm) and TF cells (TFa and/or TFb) we found 8 and 26 such genes in these cell types respectively (provided in table S7 Notes). We found that *D. simulans*, *D. sechellia* and *D. melanogaster* showed branch-site positive selection in 25.0%, 25.0% and 0% of these genes respectively for SH cells, and in 11.5%, 23.1%, and 7.7% of these genes for TF cells. These values were well above the genome-wide frequency for *D. sechellia* and *D. simulans* (although tests were conservative due to sample size, Chi-square P values for SH for *D. simulans* = 0.047 and TF for *D. sechellia* = 0.077 relative to the genome-wide values). In sum, interpreting the results in fig. 4 conservatively, we observe that upregulation of a gene in TF or SH cells is correlated with enhanced rates of positive selection in the *D. sechellia* and/or *D. simulans* lineages, regardless of whether the genes were also upregulated in another cell type (fig. 4; table S7).

While we focused on the three-species clade in fig. 4, the results for all five *melanogaster* subgroup species are provided in fig. S4. Of particular note, those results showed that 45.5% of the genes that were upregulated in the SHm cells also exhibited positive selection in the *D. yakuba* and in the *D. erecta* terminal branches (similar to *D. sechellia* in fig. 4, table S7). This suggests a history of branch-site positive selection for genes expressed in the SHm cells across outgroup branches of the phylogeny, potentially partly contributing to the divergence in ovariole numbers or functions in the two outgroup species from the three ingroup species (fig. 2).

#### Functional predictions for upregulated TF and SH genes

The studied molecular evolutionary parameters for all genes studied in fig. 4 that were upregulated in SHa, SHm, TFa, and TFp are provided in table S7. Analysis of GO-predicted functions using DAVID (Huang da, et al. 2009) showed that the genes expressed in SHa and SHm cells, such as *Jupiter* and *Timp* (table S7), were preferentially involved in microtubule formation and basement membranes (Huang da, et al. 2009), consistent with roles in TF formation (Slaidina, et al. 2020). The highly upregulated and rapidly evolving TF genes in fig. 4 and table S7 were more than threefold more common than the SH cell genes, and thus allowed us to perform functional clustering (Huang da, et al. 2009). As shown in table S8, the TF genes were preferentially associated with extracellular matrix (20.5% and 23.3% of genes from TFa and TFp respectively), basement membranes (6.8 and 10%), and 40% of genes from TFp were an integral component of membranes.

Given that the TF and SH cells types in fig. 4 (and TF and SH cells are included in the larval ovary somatic cells in table 2) have been experimentally shown to regulate the formation and number of ovarioles in *D. melanogaster* larvae (King, et al. 1968; King 1970; Godt and Laski 1995; Dansereau and Lasko 2008; Sarikaya, et al. 2012; Sarikaya and Extavour 2015; Slaidina, et al. 2020), the genes that were both highly expressed in and/or required for these ovariole functions in these cells, and exhibited rapid sequence evolution and signals of adaptive evolution (fig. 4, table 2), have the potential to directly cause an interspecies shifts in ovariole numbers. In turn, it may also be the case that the protein sequence changes observed in some of these genes may be in response to evolved shifts in ovariole numbers (potentially mediated by other ovariole-involved genes identified herein), and thus that the adaptive changes that we report here reflect the physiological intracellular changes in TFs and SH cells needed to support ovariole number changes

### Molecular Evolutionary Rates of Key Genes Predicts Ovariole Number

Finally, we conducted follow-up assessments of the main genes identified throughout our study that showed signs of high evolvability, positive selection, and involvement in *Drosophila* ovariole number divergence, to determine to what extent the molecular evolutionary characteristics of these genes were predictive of ovariole numbers in the context of *Drosophila* phylogeny. Specifically, for all genes identified from SIGNALC (N=27; table 1), from BULKSG (N=10; table 2) and from BULKSG and SINGLEC combined (N=5; table 3), we conducted a phylogenetic generalized least squares (PGLS) assessment of the relationship between ovariole number and the dN/dS values for the 41 of these 42 genes that were testable (*MtnA* was untestable due to infinity dN/dS (near zero dS, dN>0) in several branches; table 4; a summary of McDonald and Kreitman (1991) test values for all genes is shown in table S9). We found that 17 of the 41 testable genes (41.5%) showed a statistically significant relationship between ovariole number and dN/dS value (table 4; P<0.05, *CG3630* had P<0.07 and was noted in the list), indicating that dN/dS values of these genes can be a predictive factor for ovariole number per species. This further demonstrates the high effectiveness of utilizing protein sequence analysis to identify genes putatively involved in the evolution of phenotypes, similar to suggestions for other diverse traits across multiple taxa (Dorus, et al. 2004; Nadeau, et al. 2007; Ramm, et al. 2008; Wlasiuk and Nachman 2010; Luke, et al. 2014; Corso, et al. 2016; Chebbo, et al. 2021).

**Table 4.**
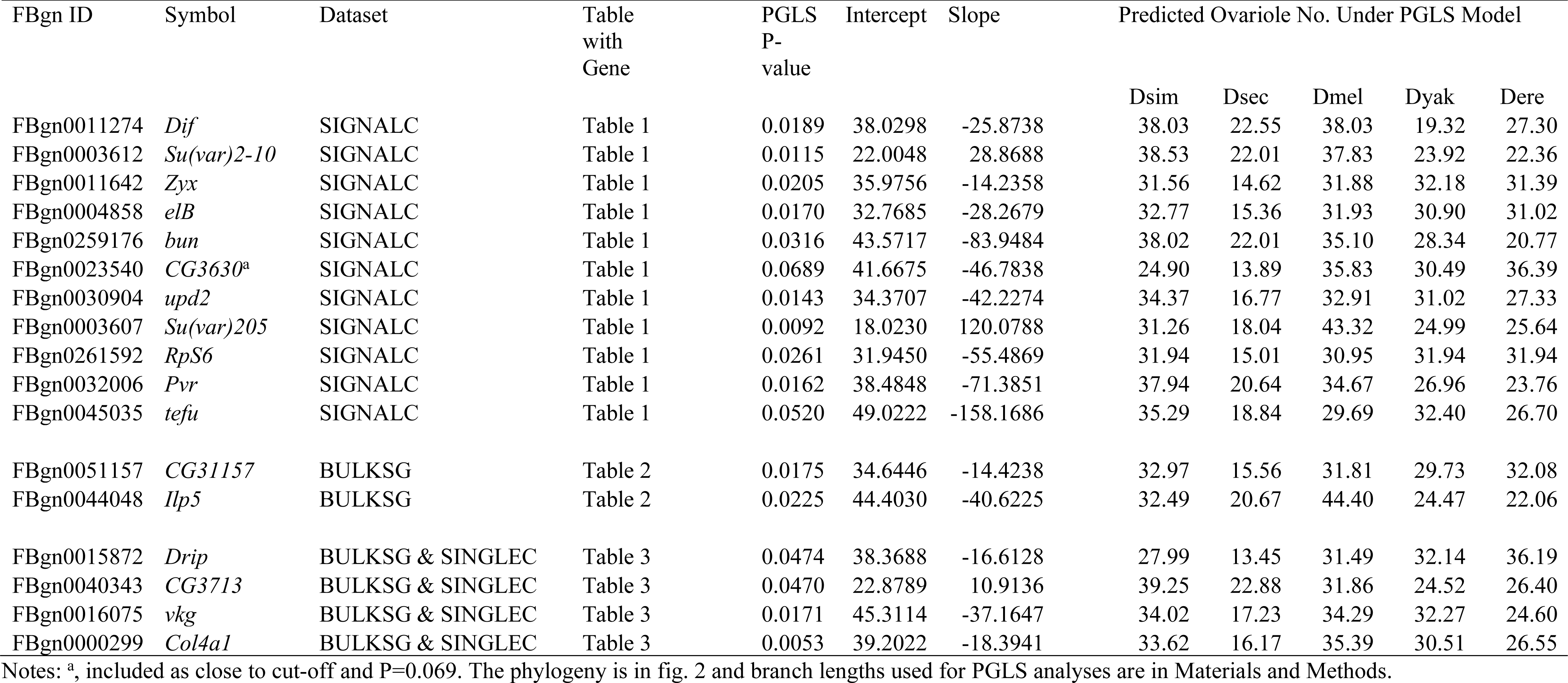
PGLS analysis of the relationship between ovariole number and dN/dS for genes putatively involved in ovariole number evolution from tables 1, 3 and 4 (42 genes total). The 17 genes that showed a relationship using PGLS are shown (P<0.05), and includes the intercept, the slope, and the predicted ovariole numbers using the model. In addition, the dataset that each gene was identified from and the table it was presented in are provided.

| FBgn ID | Symbol | Dataset | Table with Gene | PGLS P-value | Intercept | Slope | Predicted Ovariole No. Under PGLS Model |  |  |  |  |
| --- | --- | --- | --- | --- | --- | --- | --- | --- | --- | --- | --- |
|  |  |  |  |  |  |  | Dsim | Dsec | Dmel | Dyak | Dere |
| FBgn0011274 | *Dif* | SIGNALC | Table 1 | 0.0189 | 38.0298 | -25.8738 | 38.03 | 22.55 | 38.03 | 19.32 | 27.30 |
| FBgn0003612 | *Su(var)2-10* | SIGNALC | Table 1 | 0.0115 | 22.0048 | 28.8688 | 38.53 | 22.01 | 37.83 | 23.92 | 22.36 |
| FBgn0011642 | *Zyx* | SIGNALC | Table 1 | 0.0205 | 35.9756 | -14.2358 | 31.56 | 14.62 | 31.88 | 32.18 | 31.39 |
| FBgn0004858 | *elB* | SIGNALC | Table 1 | 0.0170 | 32.7685 | -28.2679 | 32.77 | 15.36 | 31.93 | 30.90 | 31.02 |
| FBgn0259176 | *bun* | SIGNALC | Table 1 | 0.0316 | 43.5717 | -83.9484 | 38.02 | 22.01 | 35.10 | 28.34 | 20.77 |
| FBgn0023540 | *CG3630*<sup>a</sup> | SIGNALC | Table 1 | 0.0689 | 41.6675 | -46.7838 | 24.90 | 13.89 | 35.83 | 30.49 | 36.39 |
| FBgn0030904 | *upd2* | SIGNALC | Table 1 | 0.0143 | 34.3707 | -42.2274 | 34.37 | 16.77 | 32.91 | 31.02 | 27.33 |
| FBgn0003607 | *Su(var)205* | SIGNALC | Table 1 | 0.0092 | 18.0230 | 120.0788 | 31.26 | 18.04 | 43.32 | 24.99 | 25.64 |
| FBgn0261592 | *RpS6* | SIGNALC | Table 1 | 0.0261 | 31.9450 | -55.4869 | 31.94 | 15.01 | 30.95 | 31.94 | 31.94 |
| FBgn0032006 | *Pvr* | SIGNALC | Table 1 | 0.0162 | 38.4848 | -71.3851 | 37.94 | 20.64 | 34.67 | 26.96 | 23.76 |
| FBgn0045035 | *tefu* | SIGNALC | Table 1 | 0.0520 | 49.0222 | -158.1686 | 35.29 | 18.84 | 29.69 | 32.40 | 26.70 |
| FBgn0051157 | *CG31157* | BULKSG | Table 2 | 0.0175 | 34.6446 | -14.4238 | 32.97 | 15.56 | 31.81 | 29.73 | 32.08 |
| FBgn0044048 | *Ilp5* | BULKSG | Table 2 | 0.0225 | 44.4030 | -40.6225 | 32.49 | 20.67 | 44.40 | 24.47 | 22.06 |
| FBgn0015872 | *Drip* | BULKSG & SINGLEC | Table 3 | 0.0474 | 38.3688 | -16.6128 | 27.99 | 13.45 | 31.49 | 32.14 | 36.19 |
| FBgn0040343 | *CG3713* | BULKSG & SINGLEC | Table 3 | 0.0470 | 22.8789 | 10.9136 | 39.25 | 22.88 | 31.86 | 24.52 | 26.40 |
| FBgn0016075 | *vkg* | BULKSG & SINGLEC | Table 3 | 0.0171 | 45.3114 | -37.1647 | 34.02 | 17.23 | 34.29 | 32.27 | 24.60 |
| FBgn0000299 | *Col4a1* | BULKSG & SINGLEC | Table 3 | 0.0053 | 39.2022 | -18.3941 | 33.62 | 16.17 | 35.39 | 30.51 | 26.55 |

Notes: <sup>a</sup>, included as close to cut-off and P=0.069. The phylogeny is in fig. 2 and branch lengths used for PGLS analyses are in Materials and Methods.

### Supplementary Analysis of a Three-Species Clade of Hawaiian *Drosophila*

While we focused on the *melanogaster* subgroup for our core analyses, as a supplementary assessment, we considered a three-species clade of Hawaiian *Drosophila* that matched our strict criteria for study (very closely related species, genome-wide data, known (and variable) ovariole numbers). We note, however, that these species are relatively distantly related to *D. melanogaster*, the species used to identify ovariole-involved genes on the basis of function and/or expression (the SIGNALC, BULKSG and SINGLESC datasets). Hawaiian *Drosophila* are paraphyletic to the *melanogaster* subgroup (Suvorov, et al. 2022), and estimates of divergence time since the last common ancestor of extant species from the two taxon groups exceed 60 Mya (Tamura, et al. 2004; Goldman-Huertas, et al. 2015). We chose the species *D. sproati* (mean 65.6 ovarioles), *D. murphyi* (mean 41.6 ovarioles) and *D. grimshawi* (mean 47.8 ovarioles) for study, with a phylogeny of: ((*D. sproati*, *D. murphyi*), *D. grimshawi*) shown in fig. S5 (Kim, et al. 2021; Suvorov, et al. 2022) (ovariole numbers from (Starmer, et al. 2003; Sarikaya, et al. 2019)). For dN/dS analysis, we focused on the ovariole-involved SIGNALC genes identified in table 1, as these signalling proteins are functionally confirmed to regulate ovariole number (Srivastava, et al. 2010; Kumar, et al. 2020). Thus, among the studied gene sets (SIGNALC, BULKSG, SINGLEC), we considered them the most appropriate for dN/dS analysis in a divergent group. We found that 21 of the 27 rapidly evolving ovariole-related genes in table 1, which were identified from study of the *melanogaster* subgroup, had a high confidence three-species orthologous gene set in the Hawaiian *Drosophila* clade (table S10). Our evaluation of branch-dN/dS values revealed that ten of the 21 genes evolved especially rapidly, with dN/dS>0.33 in at least one species terminal branch in the Hawaiian clade, which was more than two-fold higher than the genome-wide dN/dS values for the species under study (13 of 21 genes evolved rapidly using a criterion of 1.5 fold higher than the genome-wide medians; genome-wide dN/dS median values = 0.152, 0.164, and 0.160 for *D. murphyi*, *D. sproati* and *D. grimshawi* respectively; table S10; Supplementary Text File S1). Moreover, *D. sproati*, the ingroup species with highest ovariole number per female of all three Hawaiian species (fig. S5), had eight of the ten genes with dN/dS>0.33 (table S10). The ten most rapidly evolving genes included *upd2*, *CG2199*, *vn*, *elB*, *bun*, *CG3630*, *aPKC*, *H*, *Su(var)205* and *E(spl)m2-BFM*, six of which also exhibited branch-site positive selection in at least one branch. For *upd2*, we observed branch (dN/dS>1) and branch-site positive selection (P<0.05) for all three species branches (table S10), suggesting it may have a putative role in ovariole number divergence in all three species. Nonetheless, it is notable that in Table S10, eight of the 21 genes had branch-dN/dS below the aforementioned thresholds (were not 2-fold or 1.5-fold higher than the genome median) in all three Hawaiian species branches (table S10). This suggests that while these genes may be involved in ovariole functions in those taxa (as they are in *D. melanogaster* (Kumar, et al. 2020)), their protein sequence divergence may be less apt to shape interspecies shifts in ovariole numbers in these Hawaiian *Drosophila* species (table S10). Together, the data suggest that a substantial number of the rapidly evolving ovariole-involved genes in table 1, also evolved very rapidly in the Hawaiian clade, and thus may have possibly contributed to its interspecies divergence in ovariole numbers.

We also examined the Hawaiian *Drosophila* species orthologs of some of the rapidly and adaptively evolving genes in the *melanogaster* subgroup, which we identified from the SINGLEC transcription dataset (Slaidina, et al. 2020) shown in fig. 4 (and fig. S4, N values per cell type shown therein; the TF and SH cell genes are in table S7 that include certain genes from BULKSG in table 3). We hypothesized that for these genes, identified as candidate ovariole number regulators based on *D. melanogaster* expression profiles alone, it might be harder to confidently assume conservation of function in ovariole number regulation in a clade as distantly related as the Hawaiian *Drosophila* (Ranz, et al. 2003; Whittle and Extavour 2019). We therefore adopted a prudent approach, based on evaluation of the rate of high confidence ortholog detection in the Hawaiian group (see Methods and Results in Supplementary Text File S1). As shown in fig. S6, we found that genes in the TF and SH cells (fig. 4, fig. S4), had the fewest high confidence Hawaiian orthologous gene sets, as compared to genes highly expressed in the other ovarian cell types (orthologs were defined as having an ortholog found in all three Hawaiian species, and between *D. melanogaster-D. grimshawi* for gene identification). Specifically, genes upregulated in the SHa cells and those in the TFp cells (fig. S6), each had 66.6% of genes with an orthologous Hawaiian three-species orthologous gene set. In contrast, genes upregulated in CC had 85.7%, and FSCP and IC each had 82.4% (fig.S6). We speculate that genes expressed in the TF and SH cells may have evolved at a relatively higher rate (fig. 4, fig. S4, table S7) than those expressed in other ovarian cell types, making orthologs more frequently unrecognizable between *D. melanogaster* and the Hawaiian clade and/or among the three species in the Hawaiian clade (Tautz and Domazet-Loso 2011; Tautz, et al. 2013)(discussed further in Results within in Supplementary Text File S1). This rapid evolution could potentially be due to adaptive sequence changes associated with ovariole number divergence in the genus (fig. 4). It is also possible that there has been a greater propensity of genes directly involved in ovariole formation (TF and SH cells) to undergo gains and/or losses over evolutionary time (Tautz and Domazet-Loso 2011; Tautz, et al. 2013), than genes involved in regulating the other ovarian cell types. While our central focus herein was on the interspecies divergence of ovariole number and protein sequences of orthologous genes within the very closely related *Drosophila melanogaster* subgroup (table 1, table 2, table 3, table 4, fig. 4), these supplementary analyses in a Hawaiian clade provide insights into the dynamics potentially contributing to ovariole number divergence over extended time scales.”.

## Discussion

While insects exhibit a diverse number of ovarioles, including across two orders of magnitude in the genus *Drosophila* alone (Hodin and Riddiford 2000; Starmer, et al. 2003; Markow, et al. 2009; Sarikaya, et al. 2019; Church, et al. 2021), little has been known about the genetic basis of rapid interspecies divergence of this fundamental female reproductive trait. Here, we directly tackled this issue by comprehensively determining *a priori* genes with experimental and/or transcriptional evidence for roles in determining ovariole numbers or functions in *D. melanogaster* (Kumar, et al. 2020; Slaidina, et al. 2020; Tarikere, et al. 2022), and then assessing their molecular evolutionary characteristics within very closely related species in the *melanogaster* subgroup. The results revealed a highly evolvable set of ovariole-related genes that exhibited high gene-wide dN/dS and/or branch-site positive selection in patterns consistent with a role in the evolution of ovariole number divergence (table 1, table 2, table 3, table 4, table S7). Moreover, PGLS analyses supported a predictive relationship between ovariole number per species and dN/dS for many of the identified rapidly evolving ovariole-related genes (table 4). From these collective results, we propose that the rapid interspecies ovariole number divergence in *Drosophila* (fig. 2) has been facilitated by a group of highly evolvable genes with ovariole-related functions (42 identified and of focus herein, (Kumar, et al. 2020; Slaidina, et al. 2020; Tarikere, et al. 2022)) that exhibit a propensity for rapid evolution (gene-wide dN/dS) and adaptive protein sequence changes (table 1, table 2, table 3, table S7, fig. 4, fig. S4). This hypothesis is further supported by the fact that all of the ovariole-related genes revealed herein have been explicitly demonstrated to regulate ovariole number (Kumar, Blondel et al. 2020), and/or are highly and/or exclusively expressed in somatic ovarian cells whose behaviour determines ovariole number (King, et al. 1968; King 1970; Sarikaya, et al. 2012; Sarikaya, et al. 2019; Slaidina, et al. 2020; Tarikere, et al. 2022).

### Evolvability of Ovariole-Related Genes and *tau*

The evolvability, defined here as the propensity of traits or gene sequences to diverge (Wagner and Zhang 2011; Cutter and Bundus 2020), including adaptive evolution, for the ovariole-related genes identified herein for the *melanogaster* subgroup (table 1, table 2, table 3; and for the rapidly evolving ovariole genes for Hawaiian *Drosophila*, table S10), may potentially reflect fitness advantages of the fixed ovariole-related mutations, and/or may have been influenced by relaxed purifying selection. Previous studies have found that genes with high values of *tau* (Yanai, et al. 2005), which suggests low pleiotropy (Mank and Ellegren 2009; Meisel 2011; Dean and Mank 2016), may exhibit relaxed purifying selection, thereby allowing both elevated neutral protein sequence changes (and thus elevated dN/dS), and greater potential for adaptive evolution (Otto 2004; Larracuente, et al. 2008; Mank, et al. 2008; Mank and Ellegren 2009; Meisel 2011; Whittle, et al. 2021). Consistent with this pattern, we found that many of the rapidly evolving ovariole-associated genes, including those with explicit evidence of adaptive evolution from gene-wide dN/dS values larger than 1 or from branch-site positive selection tests (P<0.05), also exhibited relatively high *tau* (for example, those with values >0.90, table 1, table 1, table 3). Thus, low pleiotropy may have partly contributed to high evolvability, and enhanced adaptive potential. These events of positive selection in the ovariole-related genes (table 1, table 2, table 3, fig. 4), may have arisen by natural selection for adaption to changes in environment or oviposition substrates (Jagadeeshan and Singh 2007), and/or may have often been driven by the widely-reported and dynamic sexual behaviors of *Drosophila*, as described below.

### Putative Roles of Sexual Selection on Ovariole Number Evolution

Sexual selection may contribute to the adaptive evolution of reproductive characteristics and genes in animals (Swanson and Vacquier 2002; Clark, et al. 2009), including in *Drosophila* (Civetta and Singh 1998; Swanson, et al. 2004; Proschel, et al. 2006). Thus, one possibility is that this phenomenon may shape the evolution of ovariole-related genes observed herein (table 1, table 2, table 3, fig. 4). Different species of *Drosophila* exhibit wide variation in their reproductive behaviors (Markow and O’Grady 2005), and examples of sexual selection in the genus include intrasexual selection from sperm competition (Singh, et al. 2002; Singh and Singh 2014) and male-male (Singh and Singh 2014) and female-female competition (Bath, et al. 2018). In addition, there is evidence of intersexual selection including female- and male-mate choice (Friberg and Arnqvist 2003; LeVasseur-Viens, et al. 2015). In the latter case, if males favor larger females, a choice that may correlate with female fecundity in species where body size correlates positively with ovariole number (Bonduriansky 2001; Byrne and Rice 2006; Sinclair, et al. 2021), then this could result in positive selection on amino acid changes favoring increased ovariole numbers. Moreover, *Drosophila* exhibits sexual antagonism, which could also potentially shape female (and male) reproductive characteristics and their underlying genes (Arnqvist 1995; Rice 1996; Swanson, et al. 2004; Innocenti and Morrow 2010). For example, in *D. melanogaster*, some male reproductive traits and behaviors (e.g. seminal fluid toxicity, aggressive male re-mating behaviors) may be harmful to female reproduction and/or survival (Civetta and Clark 2000; Chapman, et al. 2001; Sirot, et al. 2014). Some studies have suggested that this could prompt female adaptive responses, and give rise to adaptive changes in the *D. melanogaster* ovaries or eggs and in the protein sequences of genes expressed in the ovaries (Civetta and Clark 2000; Jagadeeshan and Singh 2005; Sirot, et al. 2014). If this phenomenon also occurs across other members of the *melanogaster* subgroup, it may contribute to positive selection on ovariole numbers and thus on ovariole genes observed here. Significantly, sexual selection may affect reproductive phenotypes and genes (Swanson and Vacquier 2002; Proschel, et al. 2006) in a polygenic manner (Lande 1981; Coyne and Charlesworth 1997; Singh, et al. 2001; Markow and O’Grady 2005; Singh and Singh 2014), which is relevant to ovariole number evolution as this is a highly polygenic trait (Coyne, et al. 1991; Wayne and McIntyre 2002; Bergland, et al. 2008; Green and Extavour 2012; Sarikaya and Extavour 2015; Lobell, et al. 2017; Kumar, et al. 2020).

### Neutral Evolution and Ovariole Number

While we propose that our results could suggest an important role for adaptive evolution in ovariole-related genes in the interspecies divergence of ovariole numbers, it is worthwhile to consider the potential, and possibly complementary, roles of neutral evolution. Relaxed purifying selection in itself may lead to accelerated evolution and protein sequence changes (Kimura 1983; Mank and Ellegren 2009; Gossmann, et al. 2012), and to an elevated gene-wide dN/dS in a particular branch. Thus, it may be possible that some selectively neutral amino acids in ovariole-related genes were fixed via random genetic drift and affected ovariole numbers, possibly facilitated by low pleiotropy (high *tau*) (Fisher 1930; Meisel 2011; Assis, et al. 2012; Whittle, et al. 2021). Crucially however, such neutral (non-directional) changes would not be expected to yield the striking patterns we found for gene-wide dN/dS per species in ovariole-related genes and ovariole numbers (across species table 1, table 2, table 3), nor to give rise to the observed predictive relationships between dN/dS and ovariole numbers using PGLS (table 4). Moreover, our explicit evidence of adaptive evolution across many ovariole-related genes, by gene-wide dN/dS values larger than 1, branch-site positive selection analysis and McDonald and Kreitman (1991) tests (P<0.05, table 1, table 2, table 3, table S9, fig. 4, fig. S4), is unlikely to be explained by neutral evolution alone. Thus, the present data suggest that neutral evolution has not been the only or main driving factor shaping amino acid changes in ovariole-related genes in the *melanogaster* group, which we propose instead are best explained by a history of adaptive evolution.

Another factor in addition to narrow expression breadth (a factor that affects individual genes) that could in theory lead to relaxed purifying selection on nonsynonymous mutations in ovariole genes is small population size, which may affect entire genomes (Kimura 1962; Strasburg, et al. 2011; Gossmann, et al. 2012). As an example, under this scenario, relaxed selection may be expected to be more common in the *D. sechellia* lineage (fig. 2), in which the extant species has been suggested to have a smaller population size than other closely related *Drosophila* species such as *D. simulans* (Legrand, et al. 2009). Thus, we do not exclude the possibility that certain gene-wide nonsynonymous changes (dN in dN/dS) in that species branch may have contributed to its altered ovariole numbers, under an assumption that some slightly deleterious mutations may behave as selectively neutral mutations (as effective population size (N_e_) and selection coefficient (s) may yield, N_e_s<1) and be fixed by random genetic drift (Strasburg, et al. 2011; Gossmann, et al. 2012). However, as outlined above, the analyses showing affirmative branch-site positive selection tests here and the findings of gene-wide dN/dS values larger than 1 each control for neutral evolution (Zhang, et al. 2005; Yang 2007), and showed that positive selection was common in the *D. sechellia* branch (table 1, table 2, table 4,table S7, and fig. 4). Furthermore, the results revealed a high frequency of positive selection in genes upregulated in the TFs and SH cells in *D. sechellia* (fig. 4, table S7), a pattern not explainable by neutral evolution (relaxed selection) due to population size. Collectively, the evidence suggests that relaxed purifying selection, while potentially accelerating divergence rates of some ovariole-related genes studied here (Duret and Mouchiroud 2000; Mank and Ellegren 2009; Meisel 2011; Whittle, et al. 2021), may have its most significant role in the evolvability of ovariole-related genes (e.g., under high *tau*), enhancing the potential for adaptive evolution of protein sequences (Otto 2004; Larracuente, et al. 2008; Mank and Ellegren 2009; Whittle, et al. 2021), and in that manner potentially affecting interspecies ovariole number evolution.

### Evolution of Multiple Developmental Processes via Rapid Divergence of Genes that Regulate Ovariole Number

Generating the right number of ovarioles for a given species relies on multiple developmental processes that begin during embryogenesis and are not completed until puparium formation. These include establishment of a specific number of somatic gonad precursor cells in the embryonic primordial gonad, proliferation at a specific rate and to a specific degree during larval stages, morphogenetic movements including intercalation and migration to establish terminal filaments, and extracellular matrix deposition to separate ovarioles from each other within the gonad (King 1970). Any of these developmental processes could in principle be the target of evolutionary change in interspecies ovariole number divergence. Indeed, we previously showed that evolution of different developmental mechanisms underlies convergent evolution of similar ovariole numbers between or within species (Green and Extavour 2012). Accordingly, we would expect that the genes underlying these evolutionary changes might play roles in multiple different developmental processes, and this prediction is supported by our findings herein. The genes that we have identified here as not only rapidly evolving in the *melanogaster* subgroup (table1, table 2, table 3), but also with molecular evolutionary rates that are highly predictive of lineage-specific ovariole numbers (table 4), have known functional roles in cell-cell signalling, cell proliferation, cell shape change, cell migration, and extracellular matrix composition and function (table 3, table S8; see gene descriptions in Supplementary Text File S1), including in but not limited to ovariole formation in *D. melanogaster*. Further, the distinct patterns of branch-site positive selection in different lineages, suggest that ovariole number evolution involved modification of distinct developmental processes in different lineages. For example, the rapid evolution of *Zyx*, *vkg*, *col4a1*, *Ilp5,* and *CG3630* in the lineage leading to *D. sechellia* (table 1,table 2, table 3) suggests that alteration of the TF morphogenesis program was an important mechanism through which this species evolved its unusually low ovariole number (relative both to the other extant subgroup members and to its hypothesized last common ancestor (Green and Extavour 2012)). In contrast, evolutionary changes in the JAK/STAT, Wnt, EGF and Notch signaling pathways may have played a comparatively larger role in the evolution of more ovarioles in *D. simulans*, given the rapid evolution of *Su(var)2*, *CKIIbeta*, *vn*, *Gug* and *E(spl)m2-BFM* along this branch (table 1, table S3).

### Future Directions

The present study reveals a set of ovariole-involved genes, with established roles in ovariole numbers and functions, whose protein sequence divergence is linked to ovariole number divergence in the *Drosophila melanogaster* subgroup, based on a multi-layered analysis of branch-dN/dS, branch-site analyses, *tau*, and PGLS. For many genes, the branch-dN/dS value was predictive of ovariole numbers among species (table 4), consistent with an interdependent relationship. Further, our analyses of ovariole-involved genes the in Hawaiian *Drosophila* clade suggests that protein divergence of ovariole-related genes may shape ovariole number changes broadly across disparate clades of the *Drosophila* genus (table S10, fig. S6). The molecular evolutionary approach used herein may provide valuable opportunities for the discovery of genes and evolutionary processes involved in interspecies phenotype divergence, particularly important for reproductive and fitness related traits (Dorus, et al. 2004; Nadeau, et al. 2007; Ramm, et al. 2008; Wlasiuk and Nachman 2010; Luke, et al. 2014; Corso, et al. 2016; Chebbo, et al. 2021),which remains a central challenge in evolutionary developmental biology (Hoekstra and Coyne 2007; Cutter and Bundus 2020).

We suggest that future examinations of the genetic basis of interspecies divergence in ovariole number and other related reproductive traits will be most fruitfully pursued along one or more of the following major directions: First, assessments of protein sequence changes in ovariole-related genes identified here at the population level using genome-wide association studies and mutational frequency spectra (Akashi 1997; Whittle, et al. 2012; Lobell, et al. 2017), combined with McDonald-Kreitman tests (McDonald and Kreitman 1991; Murga-Moreno, et al. 2019), for multiple *Drosophila* species, will help discern evolutionary dynamics of these genes at the microevolutionary scale. Second, studies of expression divergence and functional divergence of genes in each species for the rapidly evolving ovariole-related genes identified here (table 1, table 2, table 3, table S7). Third, studies on the mating behaviors and sexual selection pressures, including male-mate-choice, female competition, and sexual antagonism, in species of the *melanogaster* subgroup (Bonduriansky 2001; Sirot, et al. 2014; Bath, et al. 2018; Veltsos, et al. 2022), will be valuable to revealing their possible links to ovariole numbers. Fourth, while we focused on the ovariole-related genes that had five-species orthologs in the *melanogaster* subgroup, ovariole number divergence may be also partly influenced by gene losses and gains in *Drosophila* lineages (Coyne and Hoekstra 2007; Tautz and Domazet-Loso 2011; Tautz, et al. 2013), as well as by genes that have diverged too rapidly to allow identification of orthologs (fig. S6) (Tautz and Domazet-Loso 2011; Tautz, et al. 2013), and thus those topics warrant further study. Finally, further research should include studies in the Hawaiian *Drosophila,* given our results suggest protein divergence of numerous ovariole-related genes may contribute to ovariole number changes in the three-species Hawaiian clade studied herein (table S10, fig. S5, fig. S6). The Hawaiian group is known for its wide phenotypic diversity in sexual characteristics, ranging from behaviours to ovariole numbers (Carson 1997; Singh and Singh 2014; Sarikaya, et al. 2019). Studies on the relationships between protein sequence changes and ovariole numbers in Hawaiian *Drosophila* will be facilitated by increased collection of whole genomes and transcriptomic data for the larval ovaries, including TFs and SH cells, and potentially by the use of expanding tools aimed to correlate gene and phenotype evolution (Kowalczyk, et al. 2019). Such research will help further decipher the genetic factors shaping the rapid evolution of ovariole numbers in the *Drosophila* genus, and thus in insects more broadly.

## Materials and Methods

### Identification of Rapidly-Evolving Ovariole-Related Genes for Follow-up Analyses

For the SIGNALC gene set, that was based on *D. melanogaster* RNAi data (Kumar, et al. 2020), we screened the 67 genes that directly affected ovariole numbers, named *hpo[RNAi]* Ovariole Number, 59 and 49 genes that affected egg laying, named *hpo[RNAi]* Egg Laying and Egg Laying *[wt]* and the 17 connector genes. For these four SIGNALC genes sets, we identified any genes with M0 dN/dS ≥1.5 higher than the genome-wide median. The cut-off was marginally lower than the BULKSG and SINGLEC because of the innate conserved nature of these signalling pathway genes, which are largely at least as old as animal divergence, in excess of 600 million years (Srivastava, et al. 2010; Kumar, et al. 2020). For the BULKSG dataset (Tarikere, et al. 2022), we screened for any differentially expressed genes that had M0 dN/dS≥0.20 in the *melanogaster* subgroup for further study. This represents a value ≥2.2 higher than the genome-wide median. With respect to the SINGLEC dataset (Slaidina, et al. 2020), for the genes with differential expression in one cell type relative to the others (P<0.05), we identified those with M0 dN/dS≥0.20, similar to the BULKSG dataset. The M0 dN/dS values for the five-species under study in the *melanogaster* subgroup were from FlyDivas (Stanley and Kulathinal 2016) that matched our own M0 dN/dS calculations in PAML (Yang 2007) (additional details on screening is available in Supplemental Text File 1).

### Follow-up Assessments: dN/dS per Species Terminal Branch, Branch-Site Positive Selection, and tau

#### Determining dN/dS for each species terminal branch

We calculated the M1 free ratios dN/dS per species terminal branch using codeml package in PAML (Yang 2007), which allows a separate dN/dS value for each branch, using as input publicly available high confidence genome-wide five-species sequence alignments from FlyDivas, which has data for various species groups of *Drosophila* (Stanley and Kulathinal 2016). Codeml is based on maximum likelihood in deriving estimates of dN/dS values, and default parameters were used in the assessments (Yang 2007). Using the dN/dS for each of the five terminal species branches, we assessed associations with respect to species transitions in ovariole numbers (terminal species branch analysis), an approach that has proven effective for determining relationships between dN/dS values and phenotypes of interest (Dorus, et al. 2004; Nadeau, et al. 2007; Wlasiuk and Nachman 2010).

We assessed the distributions of dN/dS for all studied genes per species branch (fig. S1). To affirm the suitability of the obtained data to determine dN/dS in each individual species terminal branch, we examined the magnitude of dN and dS values. The vast majority of genes had dN and dS <1.5 per species terminal branch and thus were unsaturated: 99.95 and 99.5% of genes in *D. simulans* respectively had values below this threshold, and we found even higher percentages (up to 100%) for the four other species. Only gene branches that had dN or dS >0.001 were included for further assessment to ensure sufficient divergence for study (Cusack and Wolfe 2007; Whittle, et al. 2021). The minority of cases of a branch where dN was >0.001 and dS was at or near zero were denoted simply as “dN/dS>1” (e.g., 0.2% of all 9,232 genes studied in *D. melanogaster*, 2.2% in *D. simulans*), rather than infinity (see also other approaches to cases of dS near 0 and dN>0 (Wlasiuk and Nachman 2010) and were interpreted conservatively.

#### Branch-site positive selection analysis

Branch-site codon analyses was used to assess positive selection at specific codon sites for each species terminal branch of the *melanogaster* subgroup (fig. 2) as described in the PAML manual (Yang and Nielsen 2002; Zhang, et al. 2005; Yang 2007). For all aligned genes from the *melanogaster* subgroup (N=9,237 alignments; note 9,232 had M0 values for study) (Stanley and Kulathinal 2016), including for the identified rapidly evolving ovariole-related genes, one of the five *Drosophila* species was assigned as the foreground branch in its own individual branch-site analysis. Thus, a separate branch-site analysis was conducted for all studied genes for *D. simulans*, *D. sechellia*, *D. melanogaster*, *D. yakuba* and *D. erecta*. For each gene, the maximum likelihood values were compared between a model with and without branch-site positive selection (codeml Model=2, NSsites=2, with fix_omega=1 versus 0, and P value of Chi-square for 2XΔlnL). P values <0.05 for 2XΔlnL for any gene were interpreted as evidence of positive selection at one or more codon sites in that species branch. We studied the presence or absence of branch-site positive selection within each gene, suggested by Zhang, et al. (2005), without including the post-hoc option for BEB probability analysis per codon site that has low power (Zhang, et al. 2005). The frequency of genes with branch-site positive selection in the ovariole-related gene sets under study were compared to the genome-wide frequency per species branch. Multiple test corrections were not applied as this was deemed overly conservative for our purposes of identification of ovariole-related genes with signals of positive selection, and these results were combined with other multiple layers of analyses (branch dN/dS, *tau*, and PGLS). The input tree for branch and branch-site analysis was an unrooted Newick phylogeny (unrooted version of fig. 2) as required by PAML (Yang 2007).

#### Expression specificity quantification using tau

We used the index *tau* to measure expression specificity of the genes under study here (Yanai, et al. 2005). For this, we accessed expression data from 59 tissue types and developmental stages from *D. melanogaster* (30 developmental stages and 29 tissues, table S1). The data include gene expression levels (RPKM) across development for embryos (12 stages), larvae (6 stages), pupae (6 stages) and adults (3 stages of males/females), and for major tissue types of the adult males and females (including heads, gonads, and central nervous system). The expression data were from modEncode and included the RNA-seq datasets generated by Graveley, et al. (2011) (available at: https://flybase.org/commentaries/2013_05/rna-seq_bulk.html; downloaded March 2022; see also

Supplementary Text File S1) which comprise among the widest scope of expression data available in insects (Li, et al. 2014). The *tau* value per gene was calculated as follows:

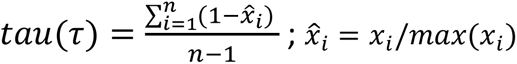

where *n*=number of tissues/stages studied, *i*= tissue/stage, *x*_*i*_= expression level of gene in tissue/stage *i*, and max (*x_i_*)= the expression level in the tissue/stage type with maximum expression (Yanai, et al. 2005).

Elevated values in one gene relative to another indicate greater expression specificity, such that most transcripts originate from few tissues/stages (see fig. S2 and Supplementary Text File S1 for an overview of the genome-wide *tau* values herein). Genes with *tau* values above 0.90 were considered highly specific in expression.

### Phylogenetic Generalized Least Squares (PGLS) Analysis

PGLS was assessed for ovariole number (dependent parameter) with respect to branch-dN/dS (independent parameter) using the five terminal species branches of the *melanogaster* subgroup (fig. 2). PGLS was conducted using the Comparative analysis of phylogenetics and evolution (Caper) package available in R (R-Core-Team 2022) (https://cran.r-project.org/web/packages/caper/index.html). The covariance matrix of species relationships was obtained under the assumption of Brownian motion using the vcv function in caper. Under a five-species tree, any genes showing P<0.05 suggest a strong relationship between ovariole number and dN/dS, sufficient to be detected under this sample size. In turn, P>0.05 does not necessarily preclude a relationship, which may be inferred from our combined analysis of dN/dS, positive selection analysis, and *tau*. The phylogenetic tree used for the covariance matrix in PGLS is shown in fig. 2.

### McDonald-Kreitman Tests

We conducted McDonald and Kreitman (1991) tests for genes of interest, using The Integrative McDonald and Kreitman test (iMKT) database (Murga-Moreno, et al. 2019). For these tests, we examined the Raleigh NC and Zambia populations, and the interspecies divergence was conducted using *D. melanogaster*-*D. simulans* contrasts (Murga-Moreno, et al. 2019). Thus, this analysis tests positive selection since divergence of the *D. melanogaster*-*D. simulans* branches only.

### Drosophila Phylogeny

To obtain the phylogeny for the five-species *melanogaster* subgroup in fig. 2, we used aligned sequence data from DrosoPhyla (Finet, et al. 2021) that contains a pre-screened dataset of 17 genes across 704 species of *Drosophilidae* (which were screened for quality, sufficient divergence, and phylogenetic informativeness). We extracted the concatenated aligned sequences for *D. simulans*, *D. sechellia*, *D. melanogaster*, *D. yakuba* and *D. erecta*, included *D. ananassae* as an outgroup as a reference (for the phylogeny construction), and removed all gaps and any sites with unknown nucleotides, yielding a total of 9,235 nucleotide sites. Using MEGA11 (Tamura, et al. 2021), we generated a maximum likelihood (ML) phylogenetic tree, including the tree lengths, based on the default parameters. We also obtained a tree using the Neighbor-Joining (NJ) Method, with nearly identical results. The relative relationships of the species in the obtained trees matched those previously observed for these five species (Obbard, et al. 2012; Finet, et al. 2021).

### Hierarchical Clustering of Expression in the SINGLEC Dataset

The relationships in gene expression across the nine different cell types of the *D. melanogaster* LL3 ovary (fig. 1A) from the SINGLEC dataset (Slaidina, et al. 2020) were assessed using hierarchical clustering under the average linkage method applied to the average standardized expression values per gene for all genes with nonzero expression (determined in Suerat v2, see Slaidina, et al. (2020)). The analysis was conducted in the Morpheus program (https://software.broadinstitute.org/morpheus).

### Gene Ontology

To study inferred gene functions and the clustering of genes by inferred function we used the program DAVID (Huang da, et al. 2009), which provides inferred gene function data for *D. melanogaster* using the FlyBase gene identifiers (Gramates, et al. 2022).

### Supplementary Analyses of a three-species Hawaiian Clade

We followed up on our main assessments of the *melanogaster* subgroup, with a supplementary evaluation of ovariole numbers and ovariole-related gene dN/dS in a three-species clade from the distantly related Hawaiian *Drosophila*, that included *D. sproati*, *D. murphyi* and *D. grimshawi*. The methods applied for CDS extraction, ortholog identification, gene alignments and dN/dS analyses for that assessment are described in Supplementary Text File S1.

## Data Availability

All data used in the present study are publicly available as described in Materials and Methods and Supplementary Text File S1.

## Acknowledgements

The authors thank members of the Extavour lab for valuable discussions. The experimental and transcriptome data generated by cited research of Dr. Tarun Kumar, Dr. Leo Blondel, Dr. Shreeharsha Tarikere and Dr. Guillem Ylla, that allowed pre-screening of genes for ovariole functions is appreciated, as well as by the authors of the sc-RNA seq datasets cited in the Materials and Methods. The authors also appreciate valuable comments by the anonymous Reviewers that helped improve the manuscript.

## Author Contributions

CAW and CGE conceived the study, wrote the manuscript and conducted analysis.

