## Supplementary Material for "Gene protein sequence evolution can predict the rapid divergence of ovariole numbers in *Drosophila*"

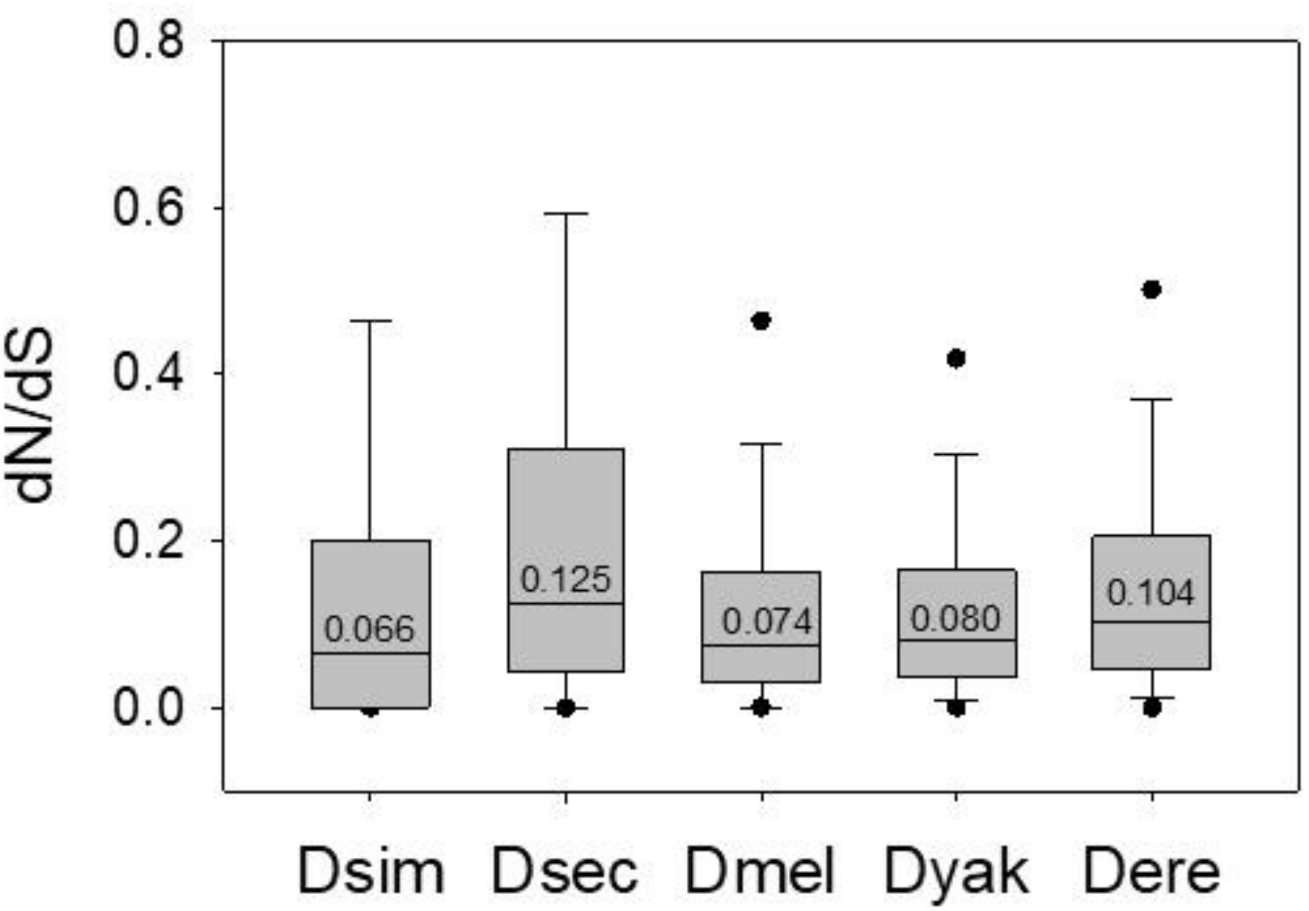

**Figure S1.** Box plots of the distribution of the free-ratio dN/dS values in each species terminal branch for all genes per genome with five-species orthologs in the *melanogaster* subgroup (Dsim = *Drosophila simulans*; Dsec = *D. sechellia*, Dmel = *D. melanogaster*, Dyak = *D. yakuba*, Dere = *D. erecta*). The median dN/dS value is shown within each box. Outliers are excluded for Dsim and Dsec as they were outside the range of the Y axis.

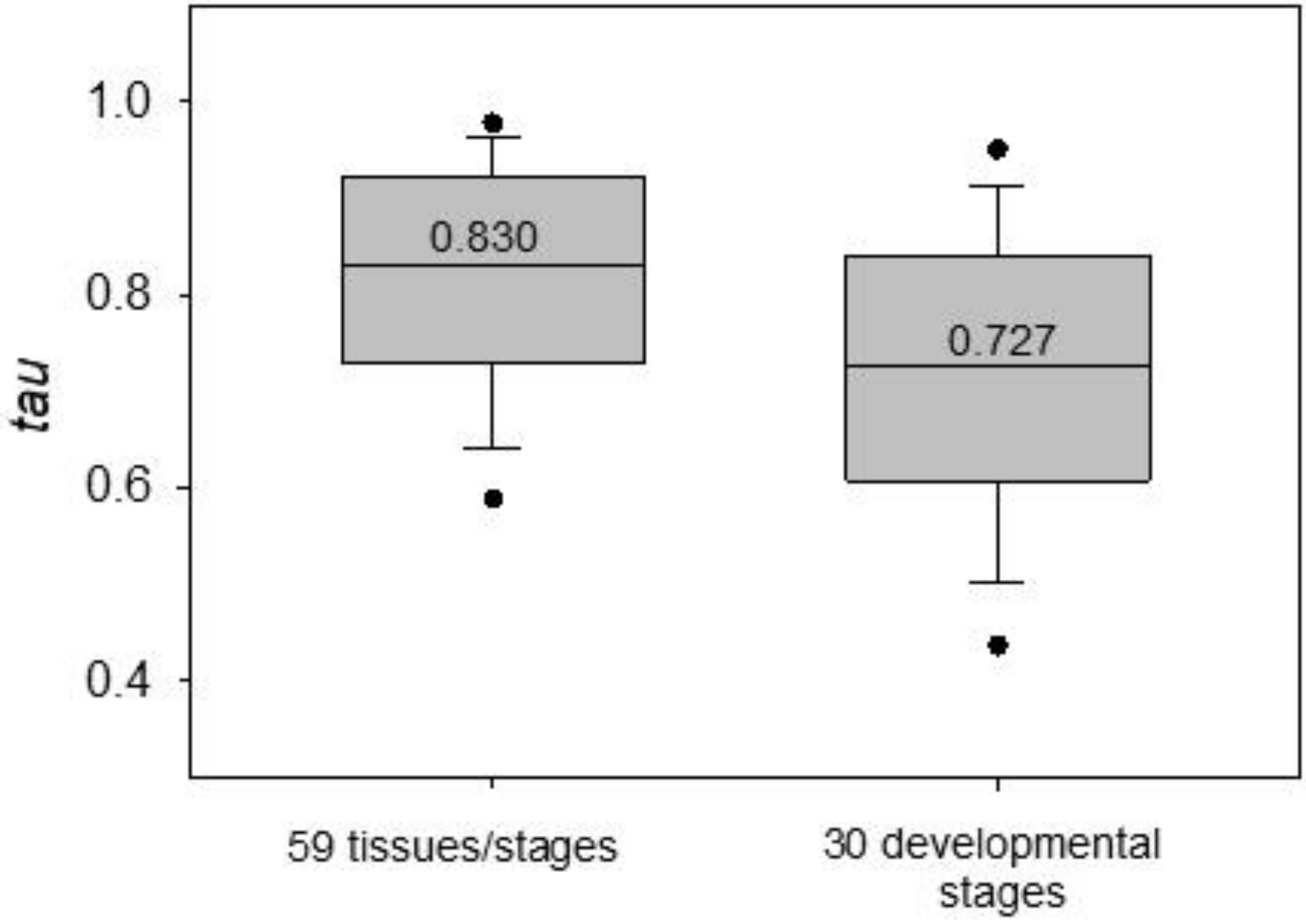

**Figure S2.** Box plots of the *tau* values across all studied genes in *D. melanogaster* using expression data across 59 tissues and developmental stages (combined) and solely using the data from 30 developmental stages. Values are for all genes with orthologs among the five studied species in the *melanogaster* subgroup. Median values are shown within bars. Tissues and stages are described in table S1.

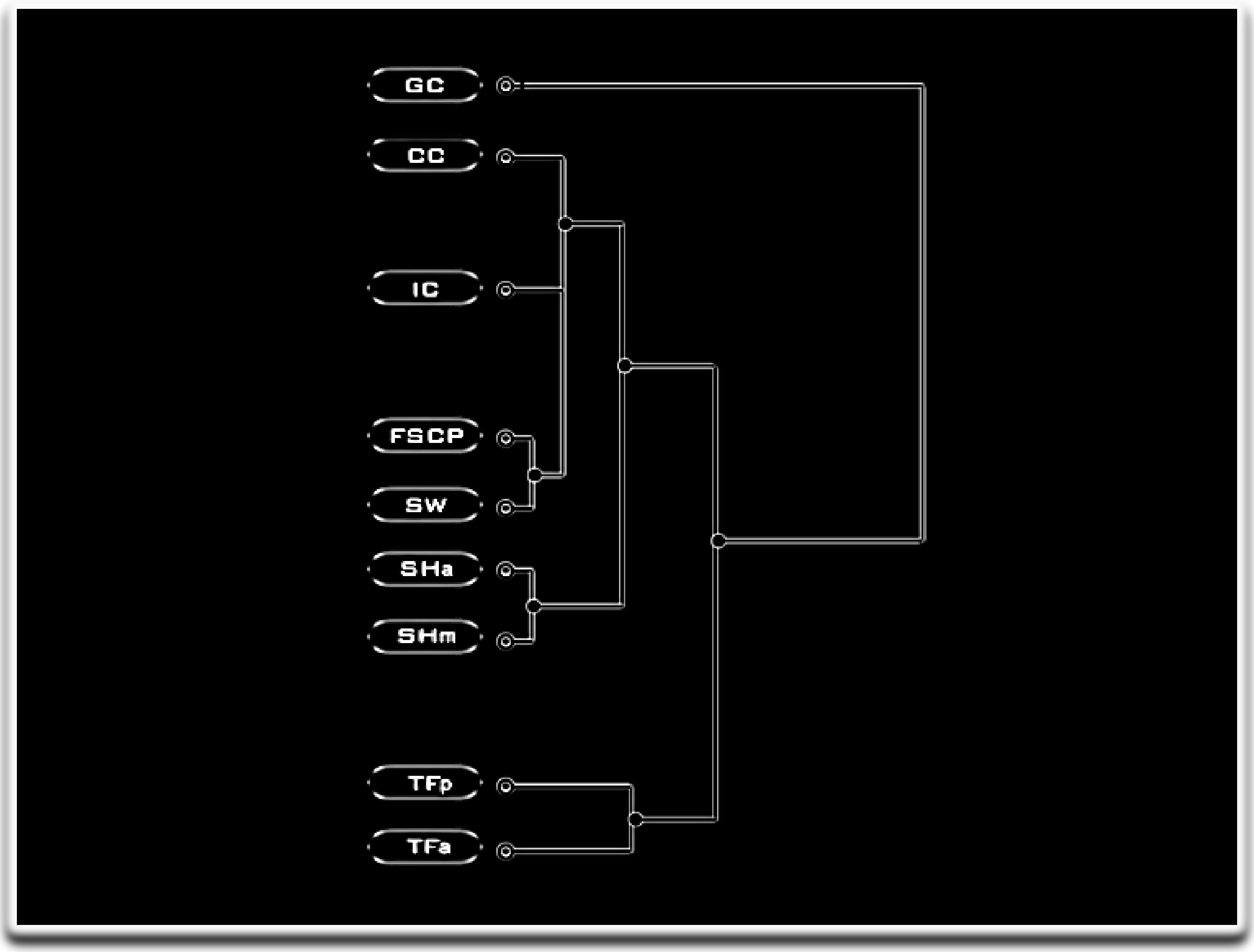

**Figure S3.** Clustering of the average standardized expression of all genes in the *D. melanogaster* genome (with non-zero expression) in the various cell types obtained from the study of LL3 larval ovaries in Slaidina, et al. (2020). The expression was clustered using hierarchical clustering and average linkage in the Morpheus program (see Materials and Methods). CC=cap cells, FSCP=follicle stem cell precursors, GC=germ cells, IC=intermingled cells, SHa= anterior sheath cells, SHm= migrating sheath cells, SW=swarm cells, TFa= anterior terminal filament cells, TFp= posterior terminal filament cells.

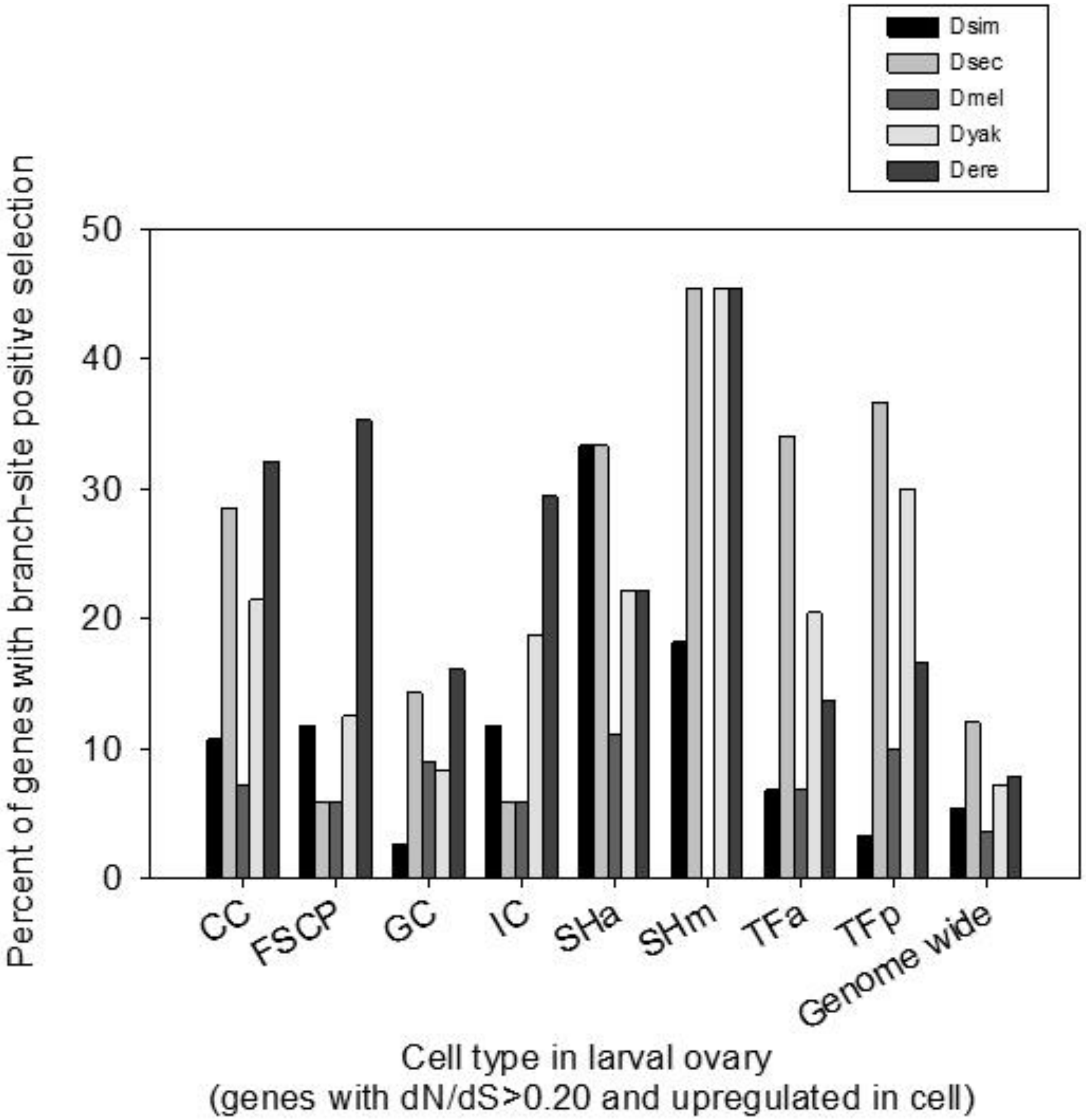

**Figure S4.** The percentage of the genes upregulated in a particular cell type using *D. melanogaster* sc-RNA seq(Slaidina, et al. 2020) and rapidly evolving in the *melanogaster* subgroup (M0 dN/dS>0.20) that exhibited branch-site positive selection in the *D. simulans* (Dsim), *D. sechellia (*Dsec), *D. melanogaster* (Dmel), *D. yakuba* (Dyak) or *D. erecta* (Dere) branches. The number of genes per category were as follows: CC (28), FSCP (17), GC (112), IC (17), SHa (9), SHm (11), SW (4), TFa (44), TFp (30). SW cells were excluded as too few genes were rapidly evolving for study. Note that a gene could be upregulated in more than one cell type. The genome-wide values are for all genes with orthologs in the *melanogaster* subgroup.

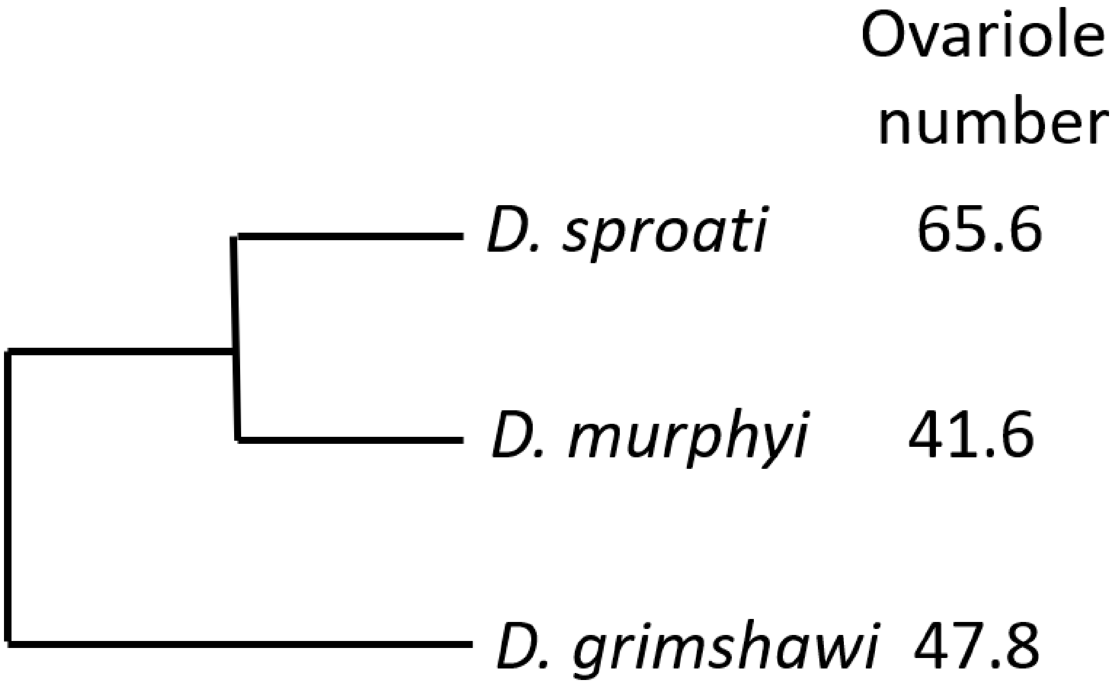

**Figure S5.** The phylogenetic relationship of the three Hawaiian *Drosophila* species under study and their mean ovariole numbers per female (phylogeny from (Kim, et al. 2021; Suvorov, et al. 2022), ovariole numbers from (Starmer, et al. 2003; Sarikaya, et al. 2019).

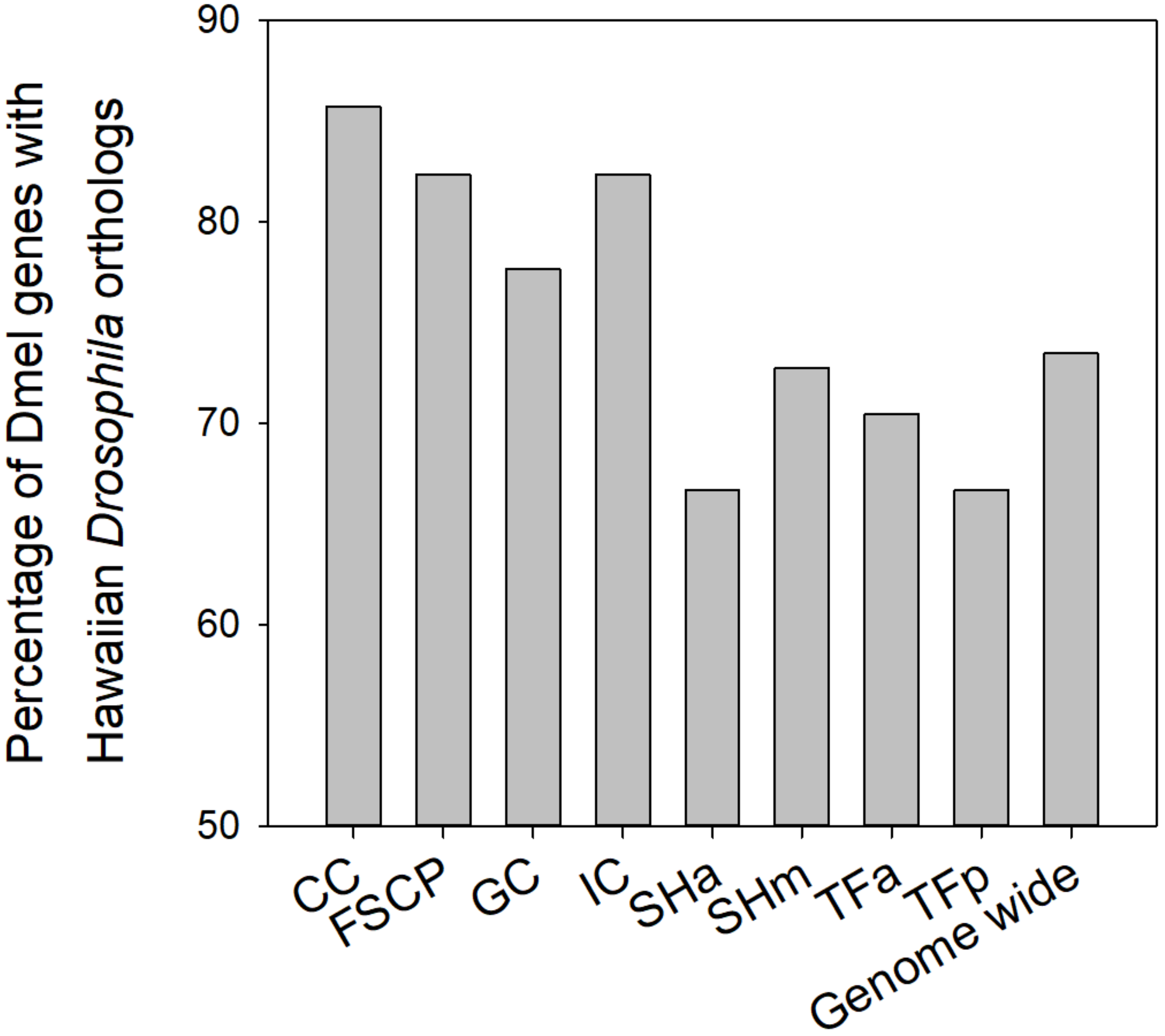

**Figure S6.** The percent of studied *D. melanogaster* sc-RNA seq genes upregulated and rapidly evolving in *D. melanogaster* that had three-species orthologs in Hawaiian *Drosophila*. N values in *D. melanogaster* are in Fig. 4. To be defined as having an orthologous gene set, the *D. melanogaster* gene had to have a match to the annotated genome *D. grimshawi*, and the three Hawaiian species had to all have high confidence orthologs to each other.

**Table S1.** The 30 developmental stages and 29 tissue types used for expression breadth analysis, or *tau*. The RPKM data is from ModEncode (Li, et al. 2014) and is available at FlyBase (Gramates, et al. 2022). Downloaded in April 2022.

| **Developmental stages** | **Tissue types** |
|---|---|
| em0-2hr | A_MateF_1d_head |
| em2-4hr | A_MateF_4d_ovary |
| em4-6hr | A_MateM_1d_head |
| em6-8hr | A_VirF_1d_head |
| em8-10hr | A_VirF_4d_head |
| em10-12hr | A_MateF_20d_head |
| em12-14hr | A_MateF_4d_head |
| em14-16hr | A_MateM_20d_head |
| em16-18hr | A_MateM_4d_acc_gland |
| em18-20hr | A_MateM_4d_head |
| em20-22hr | A_MateM_4d_testis |
| em22-24hr | A_1d_carcass |
| L1 | A_1d_dig_sys |
| L2 | A_20d_carcass |
| L3_12hr | A_20d_dig_sys |
| L3_PS1-2 | A_4d_carcass |
| L3_PS3-6 | A_4d_dig_sys |
| L3_PS7-9 | P8_CNS |
| WPP | L3_CNS |
| P5 | L3_Wand_carcass |
| P6 | L3_Wand_dig_sys |
| P8 | L3_Wand_fat |
| P9-10 | L3_Wand_imag_disc |
| P15 | L3_Wand_saliv |
| AdF_Ecl_1days | A_VirF_20d_head |
| AdF_Ecl_5days | A_VirF_4d_ovary |
| AdF_Ecl_30days | WPP_fat |
| AdM_Ecl_1days | WPP_saliv |
| AdM_Ecl_5days | P8_fat |
| AdM_Ecl_30days | |

**Table S2.** The 28 genes (27 with five-species orthologs for further follow-up study) that were identified herein as having a high dN/dS value (≥1.5 fold higher than genome-wide median) based on assessment of the gene sets that affected ovariole numbers and/or egg laying in *D. melanogaster* based on RNAi in a *hpo* loss of function genetic background from Kumar, et al. (2020). The observed phenotypes were named as *hpo[RNAi]* Ovariole number, *hpo[RNAi]* Egg laying and Egg laying *[wt]*, or as "connectors". Results are shown for dN/dS obtained for *D. melanogaster-D. simulans* (Dmel-Dsim) (genome-wide median=0.086), the *melanogaster* subgroup (median=0.089), and the *melanogaster* group (0.062) using data available from (Stanley and Kulathinal 2016) and the M0 model of PAML that provides a single dN/dS per alignment (Yang 2007). For completeness, all details for genes that belong to more than one of the four phenotype gene sets described above, and those with dN/dS in more than one of the three sets of *Drosophila* species are shown. Genes are listed in descending order with respect to the M0 dN/dS of the highest value per gene. Occasional genes near the cut-offs were retained for analysis. These genes were subjected to follow-up molecular evolutionary analyses.

| Fbgn ID | CG number | Name | Symbol | dN/dS | Species set for dN/dS | Gene phenotype observed/Status |
|---|---|---|---|---|---|---|
| FBgn0011274 | CG6794 | *Dorsal-related immunity factor* | *Dif* | 0.475 | *melanogaster* subgroup | *hpo[RNAi]* Egg laying |
| FBgn0011274 | CG6794 | *Dorsal-related immunity factor* | *Dif* | 0.311 | *melanogaster* group | *hpo[RNAi]* Egg laying |
| FBgn0014020 | CG8416 | *Rho1* | *Rho1* | 0.418 | Dmel-Dsim | *hpo[RNAi]* Ovariole number |
| FBgn0014020 | CG8416 | *Rho1* | *Rho1* | 0.418 | Dmel-Dsim | *hpo[RNAi]* Egg laying |
| FBgn0014020 | CG8416 | *Rho1* | *Rho1* | 0.418 | Dmel-Dsim | Egg laying [wt] |
| FBgn0003612 | CG8068 | *Suppressor of variegation 2-10* | *Su(var)2-10* | 0.387 | Dmel-Dsim | *hpo[RNAi]* Ovariole number |
| FBgn0003612 | CG8068 | *Suppressor of variegation 2-10* | *Su(var)2-10* | 0.387 | Dmel-Dsim | *hpo[RNAi]* Egg laying |
| FBgn0003612 | CG8068 | *Suppressor of variegation 2-10* | *Su(var)2-10* | 0.387 | Dmel-Dsim | Egg laying [wt] |
| FBgn0026379 | CG5671 | *Phosphatase and tensin homolog* | *Pten* | 0.360 | Dmel-Dsim | *hpo[RNAi]* Ovariole number |
| FBgn0026379 | CG5671 | *Phosphatase and tensin homolog* | *Pten* | 0.289 | *melanogaster* subgroup | *hpo[RNAi]* Ovariole number |
| FBgn0026379 | CG5671 | *Phosphatase and tensin homolog* | *Pten* | 0.232 | *melanogaster* group | *hpo[RNAi]* Ovariole number |
| FBgn0000259 | CG15224 | *Casein kinase II beta subunit* | *CkIIbeta* | 0.359 | Dmel-Dsim | *hpo[RNAi]* Ovariole number |
| FBgn0000259 | CG15224 | *Casein kinase II beta subunit* | *CkIIbeta* | 0.359 | Dmel-Dsim | *hpo[RNAi]* Egg laying |
| FBgn0000259 | CG15224 | *Casein kinase II beta subunit* | *CkIIbeta* | 0.359 | Dmel-Dsim | Egg laying [wt] |
| FBgn0000259 | CG15224 | *Casein kinase II beta subunit* | *CkIIbeta* | 0.195 | *melanogaster* subgroup | *hpo[RNAi]* Ovariole number |
| FBgn0000259 | CG15224 | *Casein kinase II beta subunit* | *CkIIbeta* | 0.195 | *melanogaster* subgroup | *hpo[RNAi]* Egg laying |

| | | | | | | |
|---|---|---|---|---|---|---|
| FBgn0000259 | CG15224 | *Casein kinase II beta subunit* | *CkIIbeta* | 0.195 | *melanogaster* subgroup | Egg laying [wt] |
| FBgn0035213 | CG2199 | - | *CG2199* | 0.355 | Dmel-Dsim | Connector |
| FBgn0035213 | CG2199 | - | *CG2199* | 0.334 | *melanogaster* subgroup | Connector |
| FBgn0011642 | CG32018 | *Zyxin* | *Zyx* | 0.317 | *melanogaster* subgroup | *hpo[RNAi]* Egg laying |
| FBgn0011642 | CG32018 | *Zyxin* | *Zyx* | 0.290 | Dmel-Dsim | *hpo[RNAi]* Egg laying |
| FBgn0011642 | CG32018 | *Zyxin* | *Zyx* | 0.127 | *melanogaster* group | *hpo[RNAi]* Egg laying |
| FBgn0262614 | CG43140 | *polychaetoid* | *pyd* | 0.316 | Dmel-Dsim | *hpo[RNAi]* Ovariole number |
| FBgn0031610 | CG15436 | *Parkin Interacting Substrate* | *Paris* | 0.290 | Dmel-Dsim | Connector |
| FBgn0036974 | CG5605 | *eukaryotic translation release factor 1* | *eRF1* | 0.267 | Dmel-Dsim | *hpo[RNAi]* Ovariole number |
| FBgn0036974 | CG5605 | *eukaryotic translation release factor 1* | *eRF1* | 0.267 | Dmel-Dsim | *hpo[RNAi]* Egg laying |
| FBgn0036974 | CG5605 | *eukaryotic translation release factor 1* | *eRF1* | 0.267 | Dmel-Dsim | Egg laying [wt] |
| FBgn0003984 | CG10491 | *vein* | *vn* | 0.258 | *melanogaster* subgroup | *hpo[RNAi]* Ovariole number |
| FBgn0003984 | CG10491 | *vein* | *vn* | 0.221 | *melanogaster* group | *hpo[RNAi]* Ovariole number |
| FBgn0004858 | CG4220 | *elbow B* | *elB* | 0.211 | *melanogaster* subgroup | *hpo[RNAi]* Ovariole number |
| FBgn0004858 | CG4220 | *elbow B* | *elB* | 0.138 | *melanogaster* group | *hpo[RNAi]* Ovariole number |
| FBgn0010825 | CG6964 | *Grunge* | *Gug* | 0.211 | Dmel-Dsim | *hpo[RNAi]* Ovariole number |
| FBgn0010825 | CG6964 | *Grunge* | *Gug* | 0.211 | Dmel-Dsim | *hpo[RNAi]* Egg laying |
| FBgn0010825 | CG6964 | *Grunge* | *Gug* | 0.211 | Dmel-Dsim | Egg laying [wt] |
| FBgn0002174 | CG5504 | - | *CG5504* | 0.191 | *melanogaster* subgroup | *hpo[RNAi]* Ovariole number |
| FBgn0002174 | CG5504 | - | *CG5504* | 0.124 | *melanogaster* group | *hpo[RNAi]* Ovariole number |
| FBgn0037218 | CG1107 | *auxilin* | *aux* | 0.190 | *melanogaster* subgroup | *hpo[RNAi]* Egg laying |

| | | | | | | |
|---|---|---|---|---|---|---|
| FBgn0037218 | CG1107 | *auxilin* | *aux* | 0.190 | *melanogaster* subgroup | Egg laying [wt] |
| FBgn0037218 | CG1107 | *auxilin* | *aux* | 0.157 | Dmel-Dsim | *hpo[RNAi]* Egg laying |
| FBgn0037218 | CG1107 | *auxilin* | *aux* | 0.157 | Dmel-Dsim | Egg laying [wt] |
| FBgn0037218 | CG1107 | *auxilin* | *aux* | 0.099 | *melanogaster* group | *hpo[RNAi]* Egg laying |
| FBgn0037218 | CG1107 | *auxilin* | *aux* | 0.099 | *melanogaster* group | Egg laying [wt] |
| FBgn0259176 | CG42281 | *bunched* | *bun* | 0.168 | *melanogaster* subgroup | *hpo[RNAi]* Ovariole number |
| FBgn0259176 | CG42281 | *bunched* | *bun* | 0.136 | Dmel-Dsim | *hpo[RNAi]* Ovariole number |
| FBgn0023540 | CG3630 | - | *CG3630* | 0.159 | Dmel-Dsim | Connector |
| FBgn0023540 | CG3630 | - | *CG3630* | 0.138 | *melanogaster* subgroup | Connector |
| FBgn0023540 | CG3630 | - | *CG3630* | 0.098 | *melanogaster* group | Connector |
| FBgn0261854 | CG42783 | *atypical protein kinase C* | *aPKC* | 0.184 | *melanogaster* subgroup | *hpo[RNAi]* Egg laying |
| FBgn0261854 | CG42783 | *atypical protein kinase C* | *aPKC* | 0.098 | *melanogaster* group | *hpo[RNAi]* Egg laying |
| FBgn0001169 | CG5460 | *Hairless* | *H* | 0.155 | *melanogaster* subgroup | *hpo[RNAi]* Ovariole number |
| FBgn0001169 | CG5460 | *Hairless* | *H* | 0.149 | Dmel-Dsim | *hpo[RNAi]* Ovariole number |
| FBgn0001169 | CG5460 | *Hairless* | *H* | 0.148 | *melanogaster* group | *hpo[RNAi]* Ovariole number |
| FBgn0024291 | CG5216 | *Sirtuin 1* | *Sirt1* | 0.153 | Dmel-Dsim | *hpo[RNAi]* Egg laying |
| FBgn0024291 | CG5216 | *Sirtuin 1* | *Sirt1* | 0.153 | Dmel-Dsim | Egg laying [wt] |
| FBgn0030904 | CG5988 | *unpaired 2* | *upd2* | 0.152 | *melanogaster* subgroup | *hpo[RNAi]* Ovariole number |
| FBgn0030904 | CG5988 | *unpaired 2* | *upd2* | 0.152 | *melanogaster* subgroup | *hpo[RNAi]* Egg laying |
| FBgn0030904 | CG5988 | *unpaired 2* | *upd2* | 0.152 | *melanogaster* subgroup | Egg laying [wt] |
| FBgn0030904 | CG5988 | *unpaired 2* | *upd2* | 0.114 | *melanogaster* group | *hpo[RNAi]* Ovariole number |
| FBgn0030904 | CG5988 | *unpaired 2* | *upd2* | 0.114 | *melanogaster* group | *hpo[RNAi]* Egg laying |
| FBgn0030904 | CG5988 | *unpaired 2* | *upd2* | 0.114 | *melanogaster* group | Egg laying [wt] |
| FBgn0020496 | CG7583 | *C-terminal Binding Protein* | *CtBP* | 0.148 | Dmel-Dsim | *hpo[RNAi]* Ovariole number |

| | | | | | | |
|---|---|---|---|---|---|---|
| FBgn0020496 | CG7583 | *C-terminal Binding Protein* | *CtBP* | 0.148 | Dmel-Dsim | *hpo[RNAi]* Egg laying |
| FBgn0020496 | CG7583 | *C-terminal Binding Protein* | *CtBP* | 0.148 | Dmel-Dsim | Egg laying [wt] |
| FBgn0003607 | CG8409 | *Suppressor of variegation 205* | *Su(var)205* | 0.144 | Dmel-Dsim | Connector |
| FBgn0261592 | CG10944 | *Ribosomal protein S6* | *RpS6* | 0.139 | *melanogaster* subgroup | *hpo[RNAi]* Ovariole number |
| FBgn0261592 | CG10944 | *Ribosomal protein S6* | *RpS6* | 0.139 | *melanogaster* subgroup | *hpo[RNAi]* Egg laying |
| FBgn0261592 | CG10944 | *Ribosomal protein S6* | *RpS6* | 0.139 | *melanogaster* subgroup | Egg laying [wt] |
| FBgn0020386 | CG1210 | *Phosphoinositide-dependent kinase 1* | *Pdk1* | 0.138 | Dmel-Dsim | *hpo[RNAi]* Ovariole number |
| FBgn0002592 | CG6104 | *Enhancer of split m2, Bearded family member* | *E(spl)m2-BFM* | 0.133 | *melanogaster* subgroup | *hpo[RNAi]* Ovariole number |
| FBgn0002592 | CG6104 | *Enhancer of split m2, Bearded family member* | *E(spl)m2-BFM* | 0.110 | *melanogaster* group | *hpo[RNAi]* Ovariole number |
| FBgn0032006 | CG8222 | *PDGF- and VEGF-receptor related* | *Pvr* | 0.129 | *melanogaster* subgroup | *hpo[RNAi]* Egg laying |
| FBgn0032006 | CG8222 | *PDGF- and VEGF-receptor related* | *Pvr* | 0.117 | *melanogaster* group | *hpo[RNAi]* Egg laying |
| FBgn0045035 | CG6535 | *telomere fusion* | *tefu* | 0.107 | *melanogaster* group | *hpo[RNAi]* Ovariole number |
| FBgn0045035 | CG6535 | *telomere fusion* | *tefu* | 0.107 | *melanogaster* group | Egg laying [wt] |

**Table S3.**The 27 rapidly evolving signalling and connector genes under study. For each gene, the associated pathways, *tau* and key functional terms at FlyBase (Gramates, et al. 2017) are shown. From calculation of *tau*, the x̂ for adult virgin ovaries is shown as an example of this parameter (values of 1 underlined). Genes are listed in descending order from the highest M0 dN/dS values from PAML (Table S2). Pathways are from Kumar, et al. (2020)

| **Fbgn ID** | **Gene Symbol** | **Pathways** | ***Tau* 59-All** | **$\hat{x}$ Ovary virgin** | **Key functional terms at FlyBase** |
|---|---|---|---|---|---|
| FBgn0011274 | *Dif* | Toll | 0.717 | 0.027 | Bacterial response |
| FBgn0014020 | *Rho1* | JNK, EGF, Wnt, TGFβ | 0.669 | 0.266 | Actin, cytoskeleton |
| FBgn0003612 | *Su(var)2-10* | JAK/STAT | 0.786 | 1.000 | Chromosome structure and function |
| FBgn0026379 | *Pten* | mTOR, FOXO | 0.663 | 1.000 | Controlling cytoskeletal rearrangements. |
| FBgn0000259 | *CkIIbeta* | Wnt | 0.825 | 0.356 | Functions in oogenesis, neurogenesis |
| FBgn0035213 | *CG2199* | Connector | 0.808 | 1.000 | Unknown, transcriptional regulation |
| FBgn0011642 | *Zyx* | Hippo | 0.647 | 0.220 | Actin cytoskeleton regulator |
| FBgn0262614 | *pyd* | JNK | 0.714 | 0.233 | Cytoskeleton |
| FBgn0036974 | *eRF1* | SHH | 0.567 | 0.564 | Termination of nascent peptide synthesis |
| FBgn0003984 | *vn* | EGF, Hippo | 0.740 | 0.091 | Growth and patterning of tissues |
| FBgn0004858 | *elB* | Notch | 0.739 | 0.042 | Development, cell proliferation |
| FBgn0010825 | *Gug* | Hippo | 0.706 | 0.886 | Normal developmental patterning |
| FBgn0002174 | *CG5504* | SHH | 0.843 | 0.129 | Tumor suppressor, larvae |
| FBgn0037218 | *aux* | Notch | 0.674 | 1.000 | Sperm individualization and neuron death |
| FBgn0259176 | *bun* | TGF B | 0.606 | 0.541 | Eye development and oogenesis |
| FBgn0023540 | *CG3630* | Connector | 0.856 | 0.097 | Actin binding activity, cytoskeleton, Rho activator |
| FBgn0261854 | *aPKC* | Hippo | 0.679 | 0.395 | Neuroblast proliferation and self-renewal |
| FBgn0001169 | *H* | Notch | 0.829 | 0.800 | Imaginal development |
| FBgn0024291 | *Sirt1* | FOXO | 0.671 | 0.922 | Gene silencing, apoptosis, development |
| FBgn0030904 | *upd2* | Hippo, JAK/STAT | 0.962 | 0.000 | Development |
| FBgn0020496 | *CtBP* | Wnt, Notch | 0.670 | 0.584 | Embryonic segmentation |

| | | | | | |
|---|---|---|---|---|---|
| FBgn0003607 | *Su(var)205* | Connector | 0.831 | 0.403 | Gene repression, epigenetic repression |
| FBgn0261592 | *RpS6* | mTOR | 0.658 | 0.211 | Component of small (40S) ribosomal subunit. |
| FBgn0020386 | *Pdk1* | mTOR, FOXO | 0.565 | 0.857 | Embryo development, inhibits apoptosis, spermatogenesis |
| FBgn0002592 | *E(spl)m2-BFM* | Notch | 0.871 | 0.000 | Development |
| FBgn0032006 | *Pvr* | VEGF | 0.768 | 0.034 | Cell migration regulation |
| FBgn0045035 | *tefu* | FOXO | 0.722 | 0.889 | Development, reproduction |

Notes: The *Paris* connector gene had *tau*=0.798, $\hat{x}$=1.0 and functions in the negative regulation of transcription.

**Table S4.** The 27 rapidly evolving SIGNALC genes identified from Kumar, et al. (2020) and their expression status in the soma and germ cells in the larval ovary (each pooled across stages), and among somatic cells at the early, mid and late stages (DeSeq2 P<0.01 (Tarikere, et al. 2022)). Note that if a gene is designated as upregulated in the germ cells, this automatically indicates it is downregulated in soma. A total of 25 of 27 genes showed differential expression using at least one of these comparisons.

| **Fbgn ID** | **Gene symbol** | **Upregulation observed** (Tarikere, et al. 2022) | |
|---|---|---|---|
| | | Somatic versus germ cells | Somatic cells-stage |
| FBgn0011274 | *Dif* | Soma | Early |
| FBgn0014020 | *Rho1* | Soma | - |
| FBgn0003612 | *Su(var)2-10* | - | Late |
| FBgn0026379 | *Pten* | - | Late |
| FBgn0000259 | *CkIIbeta* | - | Late |
| FBgn0035213 | *CG2199* | - | - |
| FBgn0011642 | *Zyx* | - | Late |
| FBgn0262614 | *Pyd* | Soma | Late |
| FBgn0036974 | *eRF1* | Soma | - |
| FBgn0003984 | *vn* | Soma | - |
| FBgn0004858 | *elB* | Soma | Early |
| FBgn0010825 | *Gug* | Soma | - |
| FBgn0002174 | *CG5504* | - | Early |
| FBgn0037218 | *aux* | - | - |
| FBgn0259176 | *bun* | Soma | - |
| FBgn0023540 | *CG3630* | Germ | - |
| FBgn0261854 | *aPKC* | Soma | - |
| FBgn0001169 | *H* | - | Late |
| FBgn0024291 | *Sirt1* | - | Late |
| FBgn0030904 | *upd2* | Soma | - |
| FBgn0020496 | *CtBP* | - | Late |
| FBgn0003607 | *Su(var)205* | Soma | - |
| FBgn0261592 | *RpS6* | - | Early |
| FBgn0020386 | *Pdk1* | - | Late |
| FBgn0002592 | *E(spl)m2-BFM* | Soma | - |
| FBgn0032006 | *Pvr* | Soma | Late |
| FBgn0045035 | *Tefu* | Germ | - |

Notes: the gene *Paris* (FBgn0031610), that was rapidly evolving in Dmel-Dsim but lacked high confidence orthologs in all five species was reported as upregulated in the germ cells relative to the soma cells in (Tarikere, et al. 2022).

**Table S5.** Genes that were highly upregulated in germ cells of the *D. melanogaster* larval ovary (relative to somatic cells) pooled across three larval stages (Tarikere, et al. 2022) and that exhibited rapid divergence in the *melanogaster* subgroup (M0 dN/dS>0.20). Free-ratio dN/dS per branch, branch-site positive selection and *tau* values are shown for each gene. The genes with the top 10 $\log_2$ fold change values matching these criteria are shown. Upregulation in germ cells implies downregulation in the somatic cells.

| **Fbgn ID** | **$\log_2$ Fold Change** | **Gene name** | **Gene Symbol** | **M0 dN/dS** | **Free-ratio dN/dS** | | | | | **Branch-site positive selection P<0.05** | | | | | ***tau*** |
|---|---|---|---|---|---|---|---|---|---|---|---|---|---|---|---|
| | | | | | **Dsim** | **Dsec** | **Dmel** | **Dyak** | **Dere** | **Dsim** | **Dsec** | **Dmel** | **Dyak** | **Dere** | |
| FBgn0031946 | 27.531 | *CG7164* | *CG7164* | 0.2652 | 0.0001 | 0.2922 | 0.2698 | 0.269 | 0.5347 | | | | | | 0.9678 |
| FBgn0031620 | 10.754 | *CG11929* | *CG11929* | 0.4818 | 0.8017 | 0.3598 | 0.4851 | 0.3439 | 0.5503 | | | | | yes | 0.9407 |
| FBgn0032375 | 10.728 | *CG14932* | *CG14932* | 0.6195 | >1 | 0.7898 | 2.0139 | 0.2495 | 0.4939 | | yes | yes | | | 0.9462 |
| FBgn0047199 | 10.422 | *CG31517* | *CG31517* | 1.4448 | - | >1 | >1 | >1 | 0.5721 | | | | | | 0.8702 |
| FBgn0034838 | 10.363 | *CG12782* | *CG12782* | 0.2148 | 0.4158 | 0.3893 | 0.1321 | 0.2894 | 0.2932 | | | | | | 0.9753 |
| FBgn0051475 | 10.354 | *CG31475* | *CG31475* | 0.2820 | 0.1872 | 0.4101 | 0.0408 | 0.1468 | 0.0676 | | yes | | yes | | 0.8798 |
| FBgn0034839 | 10.218 | *CG13540* | *CG13540* | 0.6728 | 0.3081 | >1 | 0.7091 | 102.6503 | 0.3557 | | | | yes | | 0.9307 |
| FBgn0051619 | 10.150 | *no long nerve cord* | *nolo* | 0.2090 | 0.327 | 1.0443 | 0.2099 | 0.2221 | 0.1103 | yes | | | yes | | 0.9218 |
| FBgn0003009 | 9.910 | *orientation disruptor* | *ord* | 0.2084 | 0.0285 | 0.2861 | 0.1592 | 0.2163 | 0.2876 | | yes | | | yes | 0.9138 |
| FBgn0039343 | 9.846 | *CG5111* | *CG5111* | 0.2013 | 0.138 | 0.1722 | 0.1472 | 0.1256 | 0.1282 | | | | | | 0.9522 |

Notes: "-" indicates the dN and dS were each <0.001 and thus too low divergence to determine dN/dS.

**Table S6.** The genes that were highly upregulated in the somatic cells of the *D. melanogaster* larval ovary in one of three stages of development (early, mid or late) as defined by Tarikere, et al. (2022) and that had elevated M0 dN/dS values in the *melanogaster* subgroup (>0.20). All differentially expressed genes with M0 dN/dS>0.20 were ranked according to $log_2$ fold change and the 30 genes with the greatest degree of upregulation at one stage of the soma are shown. Also shown is the free ratio branch dN/dS for each species, the presence of branch-site positive selection (indicated by "yes" (P<0.05)), and the *tau* value. "Stage" indicates the developmental stage in which the gene displayed upregulation.

| **Fbgn ID** | **Log$_2$ fold change** | **Stage** | **Gene name** | **Symbol** | **M0 dN/dS** | **dN/dS per branch** | | | | | **Branch-site positive selection** | | | | | ***tau*** |
|---|---|---|---|---|---|---|---|---|---|---|---|---|---|---|---|---|
| | | | | | | **Dsim** | **Dsec** | **Dmel** | **Dyak** | **Dere** | **Dsim** | **Dsec** | **Dmel** | **Dyak** | **Dere** | |
| FBgn0052057 | 8.698 | Late | *defective proboscis extension response 10* | *dpr10* | 0.2104 | 0.0001 | 0.6537 | 0.0983 | 0.1774 | 0.1875 | | yes | | | yes | 0.8784 |
| FBgn0039099 | 8.607 | Late | *Gamma-interferon-inducible lysosomal thiol reductase 2* | *GILT2* | 0.2453 | 0.7806 | 1.5581 | 0.0437 | 0.4597 | 0.1061 | | | | | | 0.9546 |
| FBgn0051296 | 8.161 | Early | *CG31296* | *CG31296* | 0.3219 | 0.3233 | 0.2084 | 0.2412 | 0.3172 | 0.5849 | | | | | yes | 0.9893 |
| FBgn0015872 | 7.798 | Late | *drip* | *Drip* | 0.2734 | 0.6248 | >1 | 0.4140 | 0.3750 | 0.1312 | yes | | yes | yes | | 0.9786 |
| FBgn0003651 | 7.644 | Late | *seven up* | *svp* | 0.2482 | 0.0001 | 0.3980 | 0.3406 | 0.0001 | 0.0001 | | | | yes | | 0.7428 |
| FBgn0085222 | 7.440 | Mid | *CG34193* | *CG34193* | 0.2104 | 0.0001 | 0.2387 | 0.4706 | 0.0001 | 0.3696 | | | | | | 0.9757 |
| FBgn0260943 | 6.235 | Late | *RNA-binding protein 6* | *Rbp6* | 0.5015 | 0.5253 | 0.0001 | 0.5512 | 0.0001 | 0.0001 | yes | | yes | | | 0.9251 |
| FBgn0040343 | 5.951 | Late | *CG3713* | *CG3713* | 0.2636 | >1 | 0.0001 | 0.8226 | 0.1501 | 0.3225 | | | | | | 0.9146 |
| FBgn0033076 | 5.417 | Late | *CG15233* | *CG15233* | 0.6252 | 0.1922 | 0.6592 | 1.2160 | 1.0832 | 0.7722 | | yes | | yes | | 0.9498 |
| FBgn0035426 | 5.001 | Late | *CG12078* | *CG12078* | 0.2513 | 0.6469 | 0.2561 | 0.2993 | 0.0911 | 0.1641 | | | | | | 0.9517 |
| FBgn0260479 | 4.938 | Mid | *CG31904* | *CG31904* | 0.3038 | 0.0001 | 0.4808 | 0.6401 | 0.0556 | 0.1363 | | yes | yes | | | 0.9466 |
| FBgn0052259 | 4.863 | Late | *CG32259* | *CG32259* | 0.9261 | >1 | 1.1050 | 0.8184 | 0.5433 | 1.6762 | | | | yes | yes | 0.9540 |
| FBgn0026315 | 4.804 | Early | *UDP-glycosyltransferase family 35 member A1* | *Ugt35A1* | 0.2279 | 0.1670 | 0.2725 | 0.2604 | 0.1125 | 0.3850 | | | | | yes | 0.7528 |
| FBgn0030776 | 4.620 | Mid | *CG4653* | *CG4653* | 0.2657 | 0.1363 | 0.2587 | 0.3934 | 0.1863 | 0.3557 | | | yes | | | 0.8962 |
| FBgn0032780 | 4.397 | Late | *CG13085* | *CG13085* | 0.2262 | 0.2711 | 0.4829 | 0.1077 | 0.1360 | 0.3106 | | | | | | 0.8017 |
| FBgn0002868 | 4.380 | Early | *Metallothionein A* | *MtnA* | 0.6883 | - | >1 | >1 | >1 | 0.1265 | | | | | | 0.8753 |
| FBgn0052246 | 4.351 | Late | *CG32246* | *CG32246* | 0.6842 | 2.9026 | 1.4628 | 0.4046 | 0.3813 | 0.8883 | | yes | | | yes | 0.9569 |
| FBgn0033541 | 4.345 | Late | *CG12934* | *CG12934* | 0.4946 | 0.0939 | 0.4220 | 0.1443 | 0.6220 | 0.7280 | | | | | yes | 0.9633 |
| FBgn0087012 | 4.305 | Early | *5-hydroxytryptamine (serotonin) receptor 2A* | *5-HT2A* | 0.3593 | 0.4309 | 0.4999 | 0.3961 | 0.5155 | 0.5550 | | | | | yes | 0.8491 |
| FBgn0032780 | 4.243 | Mid | *CG13085* | *CG13085* | 0.2262 | 0.2711 | 0.4829 | 0.1077 | 0.1360 | 0.3106 | | | | | | 0.8017 |
| FBgn0035917 | 4.230 | Late | *Z band alternatively spliced PDZ-motif protein 66* | *Zasp66* | 0.2118 | 0.0001 | 0.4500 | 0.0646 | 0.0340 | 0.0811 | | | | | | 0.8459 |
| FBgn0016075 | 4.200 | Late | *viking* | *vkg* | 0.3860 | 0.3038 | 0.7557 | 0.2965 | 0.3510 | 0.5572 | | yes | | yes | yes | 0.9037 |
| FBgn0004607 | 4.199 | Late | *Zn finger homeodomain 2* | *zfh2* | 0.2410 | 0.1792 | 0.4571 | 0.2484 | 0.2128 | 0.2496 | | yes | | | | 0.7816 |
| FBgn0039679 | 4.130 | Mid | *pickpocket 19* | *ppk19* | 0.2142 | 0.2244 | 0.1435 | 0.3729 | 0.1705 | 0.3416 | | | | | | 0.9724 |
| FBgn0039938 | 4.025 | Late | *Sox102F* | *Sox102F* | 0.2506 | 0.1676 | 0.3726 | 0.2315 | 0.1600 | 0.3848 | | | | | | 0.7231 |
| FBgn0039679 | 3.933 | Late | *pickpocket 19* | *ppk19* | 0.2142 | 0.2244 | 0.1435 | 0.3729 | 0.1705 | 0.3416 | | | | | | 0.9724 |
| FBgn0000299 | 3.875 | Late | *Collagen type IV alpha 1* | *Col4a1* | 0.4065 | 0.3034 | 1.2519 | 0.2072 | 0.4728 | 0.6879 | | yes | | yes | yes | 0.8887 |
| FBgn0003380 | 3.700 | Early | *Shaker* | *Sh* | 0.2416 | 0.4047 | 0.0001 | 0.0001 | 0.0001 | 0.0975 | yes | | | | | 0.8968 |
| FBgn0046874 | 3.519 | Late | *PFTAIRE-interacting factor 1B* | *Pif1B* | 0.3485 | 0.4171 | 0.3360 | 0.2116 | 0.3971 | 0.2675 | | | | | | 0.6881 |
| FBgn0030776 | 3.331 | Late | *CG4653* | *CG4653* | 0.2657 | 0.1363 | 0.2587 | 0.3934 | 0.1863 | 0.3557 | | | yes | | | 0.8962 |

Notes: "-" indicates the dN and dS were each <0.001 and thus too low divergence to determine dN/dS. "yes" indicates P<0.05 in branch-site selection test.

**Table S7.** Rapidly evolving genes (M0 dN/dS>0.20 in the *melanogaster* subgroup) that were statistically significantly upregulated in one cell type of the LL3 ovary relative to all other cell types from the Slaidina et al. (2020) sc-RNA seq data (P<0.05 using Seurat program v.2 (where log=$log_e$ in v2 (Satija, et al. 2015)). The genes listed include all those with M0 dN/dS>0.20 and with upregulation in the cell type provided. Genes could be upregulated in more than one cell type and thus may be listed more than once (Slaidina, et al. 2020). The subset of genes that were exclusively upregulated in the terminal filament cells (TF) or sheath cells (SH) cells are listed under Notes. The results are shown for the SHa= anterior sheath cells, SHm= migrating sheath cells, TFa= anterior terminal filament cells, TFp= posterior terminal filament cells.

| **Fbgn** | **Gene name** | **Symbol** | **Avg_logFC** | **Cell type** | **M0 dN/dS** | **dN/dS per branch** | | | | | **Branch-site positive selection** | | | | | ***tau*** |
|---|---|---|---|---|---|---|---|---|---|---|---|---|---|---|---|---|
| | | | | | | **Dsim** | **Dsec** | **Dmel** | **Dyak** | **Dere** | **Dsim** | **Dsec** | **Dmel** | **Dyak** | **Dere** | |
| **Anterior sheath cells (SHa)** | | | | | | | | | | | | | | | | |
| FBgn0051363 | *Jupiter* | *Jupiter* | 0.9101 | SHa | 0.3236 | 0.0001 | 0.3714 | >1 | 0.3439 | 0.2218 | | | | yes | | 0.8004 |
| FBgn0025879 | *Tissue inhibitor of metalloproteases* | *Timp* | 0.6981 | SHa | 0.3563 | - | 0.5309 | 0.1187 | 0.152 | 0.5264 | | yes | | | | 0.7205 |
| FBgn0033438 | *Matrix metalloproteinase 2* | *Mmp2* | 0.3692 | SHa | 0.3464 | 0.0328 | 0.3251 | 0.3121 | 0.2215 | 0.528 | | | | | yes | 0.8539 |
| FBgn0014133 | *bifocal* | *bif* | 0.3673 | SHa | 0.3763 | 0.2948 | 0.7923 | 0.1719 | 0.1024 | 0.151 | | yes | | | | 0.7583 |
| FBgn0033724 | *CG8501* | *CG8501* | 0.3626 | SHa | 0.4481 | >1 | 0.3405 | 0.1569 | 0.782 | 0.7183 | yes | | | | | 0.8952 |
| FBgn0261552 | *pasilla* | *ps* | 0.3091 | SHa | 0.3513 | 0.1772 | 1.0481 | 0.1473 | 0.1014 | 0.0519 | yes | | | yes | yes | 0.7835 |
| FBgn0260768 | *CG42566* | *CG42566* | 0.2971 | SHa | 0.2277 | - | 0.3063 | 0.5437 | 0.0807 | 0.3225 | | | yes | | | 0.8882 |
| FBgn0000719 | *folded gastrulation* | *fog* | 0.2724 | SHa | 0.3551 | 0.754 | 0.6134 | 0.5948 | 0.4491 | 0.2274 | | | | | | 0.6477 |
| FBgn0003507 | *serpent* | *srp* | 0.2635 | SHa | 0.2258 | 0.3777 | 0.2928 | 0.1872 | 0.1613 | 0.2537 | yes | yes | | | | 0.8608 |
| **Migrating sheath cells (SHm)** | | | | | | | | | | | | | | | | |
| FBgn0025879 | *Tissue inhibitor of metalloproteases* | *Timp* | 0.9614 | SHm | 0.3563 | - | 0.5309 | 0.1187 | 0.152 | 0.5264 | | yes | | | | 0.7205 |
| FBgn0051363 | *Jupiter* | *Jupiter* | 0.9497 | SHm | 0.3236 | 0.0001 | 0.3714 | >1 | 0.3439 | 0.2218 | | | | yes | | 0.8004 |
| FBgn0033724 | *CG8501* | *CG8501* | 0.5135 | SHm | 0.4481 | >1 | 0.3405 | 0.1569 | 0.782 | 0.7183 | yes | | | | | 0.8952 |
| FBgn0027932 | *A kinase anchor protein 200* | *Akap200* | 0.4686 | SHm | 0.3478 | 0.3369 | 0.2055 | 0.5401 | 0.3198 | 0.5119 | | | | | | 0.8698 |
| FBgn0000299 | *Collagen type IV alpha 1* | *Col4a1* | 0.4654 | SHm | 0.4065 | 0.3034 | 1.2519 | 0.2072 | 0.4728 | 0.6879 | | yes | | yes | yes | 0.8887 |
| FBgn0036101 | *Ninjurin A* | *NijA* | 0.3684 | SHm | 0.2904 | 0.1572 | 0.4455 | 0.0821 | 0.4177 | 0.1882 | | yes | | yes | | 0.8700 |
| FBgn0014133 | *bifocal* | *bif* | 0.3325 | SHm | 0.3763 | 0.2948 | 0.7923 | 0.1719 | 0.1024 | 0.151 | | yes | | | | 0.7583 |
| FBgn0261552 | *pasilla* | *ps* | 0.3178 | SHm | 0.3513 | 0.1772 | 1.0481 | 0.1473 | 0.1014 | 0.0519 | yes | | | yes | yes | 0.7835 |
| FBgn0016075 | *viking* | *vkg* | 0.3121 | SHm | 0.3860 | 0.3038 | 0.7557 | 0.2965 | 0.351 | 0.5572 | | yes | | yes | yes | 0.9037 |
| FBgn0033438 | *Matrix metalloproteinase 2* | *Mmp2* | 0.3011 | SHm | 0.3464 | 0.0328 | 0.3251 | 0.3121 | 0.2215 | 0.528 | | | | | yes | 0.8539 |
| FBgn0051955 | *CG31955* | *CG31955* | 0.2604 | SHm | 0.6646 | 5.8793 | 0.2682 | 0.7278 | 0.5471 | 0.7476 | | | | | yes | 0.8951 |
| **Anterior terminal filament cells (TFa)** | | | | | | | | | | | | | | | | |
| FBgn0015229 | *gliolectin* | *glec* | 1.7677 | TFa | 0.5567 | 0.14 | 0.1492 | 0.8271 | 0.4095 | 0.6083 | | | | | | 0.6855 |
| FBgn0034199 | *Growth-blocking peptide 1* | *Gbp1* | 1.6379 | TFa | 0.4162 | >1 | 0.4375 | 0.0001 | 1.1754 | 0.3524 | | | | yes | | 0.9484 |
| FBgn0025879 | *Tissue inhibitor of metalloproteases* | *Timp* | 1.4819 | TFa | 0.3563 | - | 0.5309 | 0.1187 | 0.152 | 0.5264 | | yes | | | | 0.7205 |
| FBgn0034198 | *CG11400* | *CG11400* | 1.4270 | TFa | 0.3592 | 0.0001 | 0.188 | 0.3328 | 1.3207 | 0.355 | | | | | | 0.7854 |
| FBgn0001253 | *Ecdysone-inducible gene E1* | *ImpE1* | 1.3992 | TFa | 0.2267 | 0.2315 | 0.5900 | 0.2456 | 0.1932 | 0.2401 | | yes | | | | 0.9160 |
| FBgn0034638 | *CG10433* | *CG10433* | 1.0711 | TFa | 0.2948 | 0.0001 | 0.8546 | 0.0001 | 0.1363 | 0.0362 | | | | | | 0.8591 |
| FBgn0262563 | *CG43103* | *CG43103* | 1.0423 | TFa | 0.2694 | 0.0001 | 0.3256 | 0.1094 | 0.4067 | 0.4623 | | | | yes | | 0.7215 |
| FBgn0033134 | *Tetraspanin 42El* | *Tsp42El* | 0.9933 | TFa | 0.2052 | 0.0001 | 0.133 | 0.1603 | 0.1166 | 0.39 | | | | | | 0.7592 |
| FBgn0001227 | *Heat shock gene 67Ba* | *Hsp67Ba* | 0.9834 | TFa | 0.2828 | 0.0001 | 0.7352 | 0.1914 | 0.2109 | 0.4008 | | yes | | | yes | 0.9249 |

| | | | | | | | | | | | | | | | | |
|---|---|---|---|---|---|---|---|---|---|---|---|---|---|---|---|---|
| FBgn0261822 | *Basigin* | *Bsg* | 0.9380 | TFa | 0.3411 | 0.1543 | 0.4645 | 0.0416 | 0.7968 | 0.157 | | yes | | yes | | 0.6604 |
| FBgn0038071 | *Dpp target gene* | *Dtg* | 0.8901 | TFa | 0.3946 | 0.2878 | 0.3957 | 0.503 | 0.4808 | 0.506 | | | | | | 0.8793 |
| FBgn0036101 | *Ninjurin A* | *NijA* | 0.8596 | TFa | 0.2904 | 0.1572 | 0.4455 | 0.0821 | 0.4177 | 0.1882 | | yes | | yes | | 0.8700 |
| FBgn0020503 | *Cytoplasmic linker protein 190* | *CLIP-190* | 0.7221 | TFa | 0.2379 | 0.1466 | 0.4152 | 0.2074 | 0.2021 | 0.1592 | | yes | | | | 0.6998 |
| FBgn0015872 | *Drip* | *Drip* | 0.6456 | TFa | 0.2734 | 0.6248 | >1 | 0.414 | 0.375 | 0.1312 | yes | | yes | yes | | 0.9786 |
| FBgn0003435 | *smooth* | *sm* | 0.6382 | TFa | 0.3992 | >1 | 0.6092 | 0.0309 | 0.0001 | 0.5679 | | yes | | | | 0.8599 |
| FBgn0031968 | *CG7231* | *CG7231* | 0.6366 | TFa | 0.2445 | 0.3018 | 0.6806 | 0.1559 | 0.1826 | 0.3635 | | yes | | | | 0.8268 |
| FBgn0034398 | *CG15098* | *CG15098* | 0.5951 | TFa | 0.4849 | >6 | 0.743 | 0.8262 | 0.3037 | 0.6975 | | | | | yes | 0.7146 |
| FBgn0032949 | *Lamp1* | *Lamp1* | 0.5880 | TFa | 0.3451 | 0.0001 | 0.4433 | 0.5723 | 0.3839 | 0.4147 | | | | | | 0.8346 |
| FBgn0038476 | *kugelkern* | *kuk* | 0.5606 | TFa | 0.2970 | 0.1612 | 0.3325 | 0.3852 | 0.1806 | 0.3661 | | | | | | 0.9247 |
| FBgn0000299 | *Collagen type IV alpha 1* | *Col4a1* | 0.5519 | TFa | 0.4065 | 0.3034 | 1.2519 | 0.2072 | 0.4728 | 0.6879 | | yes | | yes | yes | 0.8887 |
| FBgn0033889 | *CG6701* | *CG6701* | 0.5396 | TFa | 0.3091 | 0.245 | 1.0488 | 0.3838 | 0.288 | 0.4045 | | yes | | | | 0.6936 |
| FBgn0030174 | *CG15312* | *CG15312* | 0.5228 | TFa | 0.3686 | 0.0001 | 0.6218 | 0.1324 | 0.5161 | 0.1388 | | yes | | | | 0.8095 |
| FBgn0038682 | *CG5835* | *CG5835* | 0.4978 | TFa | 0.3324 | 0.0384 | 0.1801 | 0.6696 | 0.2774 | 0.5286 | | | | | | 0.8715 |
| FBgn0030237 | *CG15209* | *CG15209* | 0.4724 | TFa | 0.2491 | 0.2992 | 0.2840 | 0.2506 | 0.1882 | 1.1764 | | | | | | 0.7312 |
| FBgn0051361 | *defective proboscis extension response 17* | *dpr17* | 0.4614 | TFa | 0.3000 | 0.2126 | 0.2577 | 0.255 | 0.6819 | 0.3233 | | | | | | 0.8514 |
| FBgn0035020 | *CG13585* | *CG13585* | 0.4568 | TFa | 0.2637 | 0.2087 | >1 | 0.1471 | 0.0931 | 0.188 | | | | | | 0.9061 |
| FBgn0086686 | *lethal (3) L1231* | *l(3)L1231* | 0.4452 | TFa | 0.2198 | 0.1294 | 0.4058 | 0.2666 | 0.1919 | 0.2303 | | | | | | 0.6575 |
| FBgn0040343 | *CG3713* | *CG3713* | 0.4337 | TFa | 0.2636 | >1 | 0.0001 | 0.8226 | 0.1501 | 0.3225 | | | | | | 0.9146 |
| FBgn0085407 | *PDGF- and VEGF-related factor 3* | *Pvf3* | 0.4201 | TFa | 0.2440 | 0.0245 | 0.3884 | 0.0525 | 0.1055 | 0.1217 | | | | | | 0.9222 |
| FBgn0016075 | *viking* | *vkg* | 0.4074 | TFa | 0.3860 | 0.3038 | 0.7557 | 0.2965 | 0.351 | 0.5572 | | yes | | yes | yes | 0.9037 |
| FBgn0039914 | *maverick* | *mav* | 0.3770 | TFa | 0.2504 | 0.3964 | 0.532 | 0.5972 | 0.2124 | 0.4124 | | | | | | 0.6731 |
| FBgn0038498 | *beaten path IIa* | *beat-IIa* | 0.3759 | TFa | 0.2430 | 0.1713 | 0.0001 | 0.2514 | 0.3582 | 0.2106 | | | | | | 0.8736 |
| FBgn0034200 | *Growth-blocking peptide 2* | *Gbp2* | 0.3751 | TFa | 0.3622 | 0.3128 | 0.2299 | 0.3764 | 0.294 | 0.3957 | | | | | | 0.8323 |
| FBgn0027598 | *CIN85 and CD2AP related* | *cindr* | 0.3680 | TFa | 0.3297 | 0.2759 | 0.6744 | 0.0758 | 0.091 | 0.0772 | yes | yes | | | | 0.6435 |
| FBgn0035452 | *CG10359* | *CG10359* | 0.3640 | TFa | 0.2203 | 0.2327 | 0.132 | 0.3775 | 0.2047 | 0.3199 | | | | | | 0.8757 |
| FBgn0085432 | *pangolin* | *pan* | 0.3497 | TFa | 0.3218 | 0.2243 | 0.5643 | 0.1367 | 0.126 | 0.165 | yes | | | | | 0.8586 |
| FBgn0010415 | *Syndecan* | *Sdc* | 0.3348 | TFa | 0.3006 | 0.6124 | 0.6265 | 0.2136 | 0.2055 | 0.5697 | | yes | | | yes | 0.8616 |
| FBgn0045842 | *yuri gagarin* | *yuri* | 0.3305 | TFa | 0.3037 | 0.2003 | 0.3102 | 0.4189 | 0.255 | 0.2499 | | | | | | 0.9364 |
| FBgn0037350 | *Diphthamide biosynthesis 4* | *CG2911* | 0.3238 | TFa | 0.2303 | 0.0001 | 0.4093 | 0.5159 | 0.2446 | 0.1918 | | | | yes | | 0.9143 |
| FBgn0259740 | *CG42394* | *CG42394* | 0.3003 | TFa | 0.3499 | 0.0001 | 0.0001 | 2.6837 | 0.0001 | 0.2091 | | | | | | 0.7401 |
| FBgn0052803 | *Lipid droplet assembly factor 1* | *CG32803* | 0.2780 | TFa | 0.2664 | 0.0001 | 0.4282 | 0.8583 | 0.311 | 0.1959 | | | yes | | | 0.6557 |
| FBgn0033683 | *CG18343* | *CG18343* | 0.2762 | TFa | 1.0011 | 1.2209 | 0.4346 | 4.108 | 0.3598 | 1.0348 | | | yes | yes | | 0.5615 |
| FBgn0026620 | *transforming acidic coiled-coil protein* | *tacc* | 0.2659 | TFa | 0.3878 | 0.2304 | 0.5105 | 0.3432 | 0.4609 | 0.435 | | yes | | | yes | 0.5372 |
| FBgn0034724 | *babos* | *babos* | 0.2632 | TFa | 0.2450 | 0.7171 | 0.0834 | 0.2045 | 0.2332 | 0.3787 | | | | | | 0.7684 |
| **Posterior terminal filament cells (TFp)** | | | | | | | | | | | | | | | | |
| FBgn0034199 | *Growth-blocking peptide 1* | *Gbp1* | 1.6762 | TFp | 0.4162 | >1 | 0.4375 | 0.0001 | 1.1754 | 0.3524 | | | | yes | | 0.9484 |
| FBgn0000299 | *Collagen type IV alpha 1* | *Col4a1* | 1.4987 | TFp | 0.4065 | 0.3034 | 1.2519 | 0.2072 | 0.4728 | 0.6879 | | yes | | yes | yes | 0.8887 |
| FBgn0015229 | *gliolectin* | *glec* | 1.4901 | TFp | 0.5567 | 0.14 | 0.1492 | 0.8271 | 0.4095 | 0.6083 | | | | | | 0.6855 |
| FBgn0001253 | *Ecdysone-inducible gene E1* | *ImpE1* | 1.3009 | TFp | 0.2267 | 0.2315 | 0.5900 | 0.2456 | 0.1932 | 0.2401 | | yes | | | | 0.9160 |

| | | | | | | | | | | | | | | | | | |
|---|---|---|---|---|---|---|---|---|---|---|---|---|---|---|---|---|---|
| FBgn0025879 | *Tissue inhibitor of metalloproteases* | *Timp* | 1.1705 | TFp | 0.3563 | - | 0.5309 | 0.1187 | 0.152 | 0.5264 | | | yes | | | | 0.7205 |
| FBgn0262563 | *CG43103* | *CG43103* | 1.1400 | TFp | 0.2694 | 0.0001 | 0.3256 | 0.1094 | 0.4067 | 0.4623 | | | | | yes | | 0.7215 |
| FBgn0002868 | *Metallothionein A* | *MtnA* | 1.0706 | TFp | 0.6883 | - | >1 | >1 | >1 | 0.1265 | | | | | | | 0.8753 |
| FBgn0033134 | *Tetraspanin 42El* | *Tsp42El* | 0.9692 | TFp | 0.2052 | 0.0001 | 0.133 | 0.1603 | 0.1166 | 0.39 | | | | | | | 0.7592 |
| FBgn0016075 | *viking* | *vkg* | 0.9189 | TFp | 0.3860 | 0.3038 | 0.7557 | 0.2965 | 0.351 | 0.5572 | | | yes | | yes | yes | 0.9037 |
| FBgn0034198 | *CG11400* | *CG11400* | 0.8837 | TFp | 0.3592 | 0.0001 | 0.188 | 0.3328 | 1.3207 | 0.355 | | | | | | | 0.7854 |
| FBgn0001227 | *Heat shock gene 67Ba* | *Hsp67Ba* | 0.8687 | TFp | 0.2828 | 0.0001 | 0.7352 | 0.1914 | 0.2109 | 0.4008 | | | yes | | | yes | 0.9249 |
| FBgn0034638 | *CG10433* | *CG10433* | 0.7609 | TFp | 0.2948 | 0.0001 | 0.8546 | 0.0001 | 0.1363 | 0.0362 | | | | | | | 0.8591 |
| FBgn0015872 | *Drip* | *Drip* | 0.7607 | TFp | 0.2734 | 0.6248 | >1 | 0.414 | 0.375 | 0.1312 | yes | | | yes | yes | | 0.9786 |
| FBgn0036101 | *Ninjurin A* | *NijA* | 0.7144 | TFp | 0.2904 | 0.1572 | 0.4455 | 0.0821 | 0.4177 | 0.1882 | | | yes | | yes | | 0.8700 |
| FBgn0259740 | *CG42394* | *CG42394* | 0.6697 | TFp | 0.3499 | 0.0001 | 0.0001 | 2.6837 | 0.0001 | 0.2091 | | | | | | | 0.7401 |
| FBgn0031968 | *CG7231* | *CG7231* | 0.5292 | TFp | 0.2445 | 0.3018 | 0.6806 | 0.1559 | 0.1826 | 0.3635 | | | yes | | | | 0.8268 |
| FBgn0035020 | *CG13585* | *CG13585* | 0.4914 | TFp | 0.2637 | 0.2087 | >1 | 0.1471 | 0.0931 | 0.188 | | | | | | | 0.9061 |
| FBgn0052803 | *Lipid droplet assembly factor 1* | *CG32803* | 0.4869 | TFp | 0.2664 | 0.0001 | 0.4282 | 0.8583 | 0.311 | 0.1959 | | | | yes | | | 0.6557 |
| FBgn0003435 | *smooth* | *sm* | 0.4832 | TFp | 0.3992 | >1 | 0.6092 | 0.0309 | 0.0001 | 0.5679 | | | yes | | | | 0.8599 |
| FBgn0038476 | *kugelkern* | *kuk* | 0.4056 | TFp | 0.2970 | 0.1612 | 0.3325 | 0.3852 | 0.1806 | 0.3661 | | | | | | | 0.9247 |
| FBgn0010415 | *Syndecan* | *Sdc* | 0.4054 | TFp | 0.3006 | 0.6124 | 0.6265 | 0.2136 | 0.2055 | 0.5697 | | | yes | | | yes | 0.8616 |
| FBgn0261822 | *Basigin* | *Bsg* | 0.3741 | TFp | 0.3411 | 0.1543 | 0.4645 | 0.0416 | 0.7968 | 0.157 | | | yes | | yes | | 0.6604 |
| FBgn0033683 | *CG18343* | *CG18343* | 0.3315 | TFp | 1.0011 | 1.2209 | 0.4346 | 4.108 | 0.3598 | 1.0348 | | | | yes | yes | | 0.5615 |
| FBgn0034200 | *Growth-blocking peptide 2* | *Gbp2* | 0.2992 | TFp | 0.3622 | 0.3128 | 0.2299 | 0.3764 | 0.294 | 0.3957 | | | | | | | 0.8323 |
| FBgn0038498 | *beaten path IIa* | *beat-IIa* | 0.2919 | TFp | 0.2430 | 0.1713 | 0.0001 | 0.2514 | 0.3582 | 0.2106 | | | | | | | 0.8736 |
| FBgn0035452 | *CG10359* | *CG10359* | 0.2855 | TFp | 0.2203 | 0.2327 | 0.132 | 0.3775 | 0.2047 | 0.3199 | | | | | | | 0.8757 |
| FBgn0034398 | *CG15098* | *CG15098* | 0.2681 | TFp | 0.4849 | >1 | 0.743 | 0.8262 | 0.3037 | 0.6975 | | | | | | yes | 0.7146 |
| FBgn0030174 | *CG15312* | *CG15312* | 0.2649 | TFp | 0.3686 | 0.0001 | 0.6218 | 0.1324 | 0.5161 | 0.1388 | | yes | | | | | 0.8095 |
| FBgn0038071 | *Dpp target gene* | *Dtg* | 0.2564 | TFp | 0.3946 | 0.2878 | 0.3957 | 0.503 | 0.4808 | 0.506 | | | | | | | 0.8793 |
| FBgn0043841 | *virus-induced RNA 1* | *vir-1* | 0.2541 | TFp | 0.2419 | 0.1295 | 0.111 | 0.1677 | 0.4729 | 0.3974 | | | | | yes | | 0.7401 |

Notes: Genes that were exclusively upregulated in the SH cells: FBgn0051363, FBgn0033438, FBgn0014133, FBgn0033724, FBgn0000719, FBgn0003507, FBgn0027932, FBgn0051955. Genes that were exclusively upregulated in the TFs: FBgn0001253, FBgn0003435, FBgn0015872, FBgn0020503, FBgn0026620, FBgn0027598, FBgn0030174, FBgn0030237, FBgn0033134, FBgn0033683, FBgn0034198, FBgn0034200, FBgn0034638, FBgn0037350, FBgn0038071, FBgn0038476, FBgn0038498, FBgn0038682, FBgn0039914, FBgn0040343, FBgn0045842, FBgn0051361, FBgn0085407,FBgn0085432, FBgn0086686, FBgn0262563. Branch-site positive selection in *D. simulans*, *D. sechellia* and *D. melanogaster* using the subset of genes exclusively upregulated in SH cells was 25.0, 25.0 and 0% of studied genes respectively and for TFs was 11.5, 23.1, and 7.7% respectively. "-" indicates the dN and dS were each <0.001 and thus too low divergence to determine dN/dS.

**Table S8.** The top GO functional classifications of rapidly evolving genes that were upregulated in TFa and TFp cell types. Classifications were determined using DAVID (Huang da, et al. 2009) for all genes per category. P-values indicate the statistical significance of preferential classification of the gene into each GO group. Genes could match more than one GO group.

| Functional class | Percent of genes matching | P-value |
|---|---|---|
| **Terminal filament cells anterior (a)** | | |
| basement membrane organization | 6.8 | 9.10E-04 |
| cytokine activity | 6.8 | 1.00E-03 |
| extracellular space | 20.5 | 2.80E-03 |
| cell adhesion | 6.8 | 1.50E-02 |
| growth factor activity | 4.5 | 4.30E-02 |
| extracellular matrix structural constituent | 4.5 | 5.80E-02 |
| **Terminal filament cells posterior (p)** | | |
| basement membrane organization | 10 | 3.70E-04 |
| extracellular space | 23.3 | 4.10E-03 |
| cell adhesion | 10 | 6.50E-03 |
| collagen type IV trimer | 6.7 | 6.80E-03 |
| cytokine activity | 6.7 | 2.90E-02 |
| extracellular matrix structural constituent | 6.7 | 3.60E-02 |
| integral component of membrane | 40 | 3.80E-02 |
| axon guidance | 10 | 3.90E-02 |
| extracellular matrix | 10 | 4.20E-02 |
| | 6.7 | 4.40E-02 |

**Table S9**. The results for the McDonald and Kreitman (1991)tests of positive selection for the 42 genes identified in Tables 1, 2, and 3. Only those genes with P<0.05 are shown. The *D. melanogaster* populations examined are the Raleigh North Carolina (RAL) and the Zambia population. The P-values are shown as MK-P. Alpha ($\alpha$) indicates the proportion of substitutions apt to have experienced positive selection. Analysis was conducted using the integrative McDonald Kreitman test (Murga-Moreno, et al. 2019) that uses *D. simulans* as the comparison species (Dmel-Dsim). Note that iMKT may not necessarily have the exact same alignment per gene as FlyDivas (Stanley and Kulathinal 2016), which was used for all our core analyses, but the alignments from both analyses are highly similar and representative per gene for this supplementary assessment. Data are available publicly for each dataset (Stanley and Kulathinal 2016; Murga-Moreno, et al. 2019). MK-P indicates the P-value.

| | | **RAL population** | | **Zambia population** | |
|---|---|---|---|---|---|
| Fbg ID | Gene | $\alpha$ | MK-P | $\alpha$ | MK-P |
| FBgn0026379 | *Pten* | 0.880 | 0,012 | 0.771 | 0.030 |
| FBgn0004858 | *elB* | 0.756 | 0.057 | 0.703 | 0.001 |
| FBgn0010825 | *Gug* | 0.530 | 0.010 | 0.598 | 0 |
| FBgn0261854 | *aPKC* | 0.969 | 0 | 0.944 | 0 |
| FBgn0032006 | *Pvr* | 0.558 | 0.020 | 0.525 | 0.014 |
| FBgn0039108 | *CG10232* | 0.632 | 0.031 | 0.669 | 0.002 |
| FBgn0016075 | *vkg* | 0.681 | 0 | 0.769 | 0 |
| FBgn0000299 | *Col4a1* | 0.711 | 0,001 | 0,757 | 0 |

**Table S10.** The branch dN/dS and branch-site analysis Hawaiian *Drosophila* (*D. murphyi*, *D. sproati* and *D. grimshawi*) for orthologs to the SIGNALC genes identified in table 1 that exhibited signs of involvement in ovariole number evolution (from analysis of the *melanogaster* subgroup, and (Kumar, et al. 2020)). Genes having a dN/dS value of >1 at the gene-wide level and those with values above 0.330 in at least one species branch are each shown (that is double the median of the genome with the highest median dN/dS, *D. sproati*). Species that exhibited positive selection using branch-site (BR-S) analysis in PAML (P<0.05) are indicated by the abbreviated species name.

| Fbgn | Gene name | Symbol | Hawaiian branch-dN/dS | | | BR-S positive selection | | |
|---|---|---|---|---|---|---|---|---|
| | | | *D. murphyi* | *D. sproati* | *D. grimshawi* | | | |
| **Genes with dN/dS>0.330 in at least one branch** | | | | | | | | |
| FBgn0030904 | *unpaired 2* | *upd2* | **2.7852** | 1.6904 | 1.1397 | Dmur | Dspr | Dgri |
| FBgn0035213 | *CG2199* | *CG2199* | **0.637** | 0.5455 | 0.5134 | | | |
| FBgn0003984 | *vein* | *vn* | 0.1162 | **0.3489** | **0.3484** | | | Dgri |
| FBgn0004858 | *elbow B* | *elB* | 0.0738 | **0.4486** | 0.0744 | | Dspr | |
| FBgn0259176 | *bunched* | *bun* | 0.5683 | 0.3779 | **0.7306** | Dmur | | Dgri |
| FBgn0023540 | *CG3630* | *CG3630* | **0.5018** | 0.2986 | 0.3435 | | | |
| FBgn0261854 | *atypical protein kinase C* | *aPKC* | 0.1756 | 0 | **0.5173** | | | Dgri |
| FBgn0001169 | *Hairless* | *H* | 0.3588 | **0.8827** | 0.4621 | | | Dgri |
| FBgn0003607 | *Suppressor of variegation 205* | *Su(var)205* | 0.4541 | **0.4620** | 0.1688 | | | |
| FBgn0002592 | *Enhancer of split m2, Bearded family member* | *E(spl)m2-BFM* | NA | **0.3478** | 0.2298 | | | |
| **Genes with all branches dN/dS≤0.330** | | | | | | | | |
| FBgn0014020 | *Rho1* | *Rho1* | NA | NA | 0 | | | |
| FBgn0003612 | *Suppressor of variegation 2-10* | *Su(var)2-10* | 0.0001 | 0.0001 | 0.2512 | | | Dgri |
| FBgn0000259 | *Casein kinase II beta subunit* | *CkIIbeta* | 0.0001 | NA | 0.0001 | | | |
| FBgn0262614 | *polychaetoid* | *pyd* | 0.191 | 0.0652 | 0.0777 | Dmur | | |
| FBgn0036974 | *eukaryotic translation release factor 1* | *eRF1* | NA | 0 | 0.0001 | | | |
| FBgn0002174 | *CG5504* | *CG5504* | 0.1637 | 0.1607 | 0.2089 | | | Dgri |
| FBgn0037218 | *auxilin* | *aux* | 0.2491 | 0.1765 | 0.1276 | | | |

| | | | | | |
|---|---|---|---|---|---|
| FBgn0024291 | *Sirtuin 1* | *Sirt1* | 0.2621 | 0.294 | 0.2123 |
| FBgn0020496 | *C-terminal Binding Protein* | *CtBP* | 0 | 0.0001 | 0.0001 |
| FBgn0261592 | *Ribosomal protein S6* | *RpS6* | 0.0001 | 0.0001 | 0.0274 |
| FBgn0045035 | *telomere fusion* | *tefu* | 0.2727 | 0.1711 | 0.2456 |
| **No Orthologs** | | | | | |
| FBgn0011274 | *Dorsal-related immunity factor* | *Dif* | No Orth-HD | | |
| FBgn0026379 | *Phosphatase and tensin homolog* | *Pten* | No Orth-HD | | |
| FBgn0011642 | *Zyxin* | *Zyx* | No Orth-HD | | |
| FBgn0010825 | *Grunge* | *Gug* | No Orth-HD | | |
| FBgn0020386 | *Phosphoinositide-dependent kinase 1* | *Pdk1* | No Orth-HD | | |
| FBgn0032006 | *PDGF- and VEGF-receptor related* | *Pvr* | No Orth-HD | | |

Notes: No Orth-HD=no high confidence reciprocal BLASTX orthologs identified among the three Hawaiian *Drosophila* (HD). NA indicates the branch did not meet the criteria that dN or dS>0.001 for dN/dS analysis. Ten genes had dN/dS>2 fold higher (value>0.330) than the genome-wide medians in at least one species branch (the branch with the highest dN/dS value is in bold). Three additional genes, *aux*, *Sirt1*, and *tefu* had an elevated value when using a criterion of dN/dS >1.5 fold higher (value>0.246) than the genome-wide medians.

### SUPPLEMENTARY TEXT FILE S1

#### Supplementary Methods for the *melanogaster* subgroup

##### Identification of Rapidly Evolving Ovariole-Related Genes

With respect to the SIGNALC dataset (Kumar, et al. 2020), for each of four gene sets identified therein as having roles in regulating egg-laying and/or ovariole numbers, namely *hpo[RNAi]* Ovariole Number, *hpo[RNAi]* Egg Laying, Egg Laying [*wt*], and the so-called "Connectors" (genes not included in the initial screen but identified as candidates by network topology analysis), we identified the subset of genes that had orthologs in all five species of the *melanogaster* subgroup and exhibited a markedly elevated M0 dN/dS value relative to the genome-wide values (Yang 2007) using data from FlyDivas (Stanley and Kulathinal 2016). Given that signalling genes are typically components of complex pathways and have pleiotropic roles in a wide range of cells or tissues (Cui, et al. 2009; Kumar, et al. 2020), they may be expected to typically, but, not always (Cui, et al. 2009; Darfour-Oduro, et al. 2016), have been subjected to strong purifying selection and thus evolve slowly in their protein sequence (Cui, et al. 2009; Mank and Ellegren 2009; Meisel 2011; Assis, et al. 2012; Darfour-Oduro, et al. 2016; Masalia, et al. 2017; Whittle, et al. 2021). We therefore defined rapidly evolving ovariole-related genes as those having a value of ≥1.5 fold above the genome-wide median M0 dN/dS in the *melanogaster* subgroup (genome-wide median M0 dN/dS=0.0887, N=9,232). The cut-off was 1.5 (M0 dN/dS value ≥0.133) rather than higher, as for the other studied datasets (see below, BULKSG and SINGLEC), given the innate conserved nature of these genes. In addition, for this one dataset, the SIGNALC, to identify the widest scope possible of genes with a propensity for rapid evolution for follow-up study, we also included any genes with ≥1.5 fold higher M0 dN/dS in the *D. simulans*-*D. melanogaster* pair (≥1.5 fold higher than the median=0.857, N=10,765 aligned genes) and the six-species *melanogaster* group (which includes *D. ananassae* as a outgroup, fig. 2; genome-wide median=0.062, N=8,696 genes) (Stanley and Kulathinal 2016). These genes were added to the gene list for our follow-up study within the *melanogaster* subgroup (fig. 2).

For the BULKSG transcription dataset (Tarikere, et al. 2022), we identified and extracted all genes that were statistically significantly upregulated (using DeSeq2 $P<0.01$ criterion in that assessment (Love, et al. 2014)) in the *D. melanogaster* pooled larval ovary somatic cells and those upregulated in the pooled germ cells. Upregulated in germ cells automatically indicates downregulation in soma cells, and thus both up- and downregulated genes in soma were studied. We also extracted for study those genes that were upregulated at a single one of the larval stages studied (early, mid, or late) for the pooled ovary somatic cells; these stages correspond to different stages of TF formation (Tarikere, et al. 2022). For these gene sets, we then identified those genes with M0 dN/dS>0.20 in the *melanogaster* subgroup, which represent a value ≥2.2 fold higher than the genome-wide median. The cut-off was higher for the BULKSG than SIGNALC dataset, given that we examined all genes in the genome that were differentially expressed in the former dataset, which are apt to be less conserved as a group than the SIGNALC genes. The genes matching these criteria (dN/dS>0.20, and up- or downregulated in soma cells), were ranked based on degree of differential expression, and subjected to follow-up analysis in the *melanogaster* subgroup.

With respect to the SINGLEC dataset (Slaidina, et al. 2020), genes were extracted for further study that were statistically significantly differentially transcribed in a specific cell type of the LL3 larval ovary relative to all other cell types based on sc-RNA seq (using average standardized expression and P-values from Seurat v.2; some genes were upregulated in more than one cell type based on these criteria (Slaidina, et al. 2020)). The nine cell types studied (shown in fig. 1) included the germ cells (GC) and eight somatic cell types, namely the cap cells (CC), follicle stem cell precursors (FSCP), intermingled cells (IC), anterior sheath cells (SHa), migrating sheath cells (SHm), swarm cells (SW), anterior terminal filaments (TFa), and posterior terminal filaments (TFp), with a particular focus on the two types of SH cells and the two types of TF cells, which are thought to largely shape species-specific ovariole numbers and function (King, et al. 1968; Sarikaya, et al. 2012; Sarikaya and Extavour 2015; Slaidina, et al. 2020). Similar to the BULKSG assessment, among the genes that were upregulated per cell type, we identified the genes that had signs of rapid protein sequence divergence (M0 dN/dS >0.20 in genes with five-species orthologs) in the *melanogaster* subgroup (Stanley and Kulathinal 2016; Gramates, et al. 2017), for further analyses.

#### PGLS Analysis

Any species branch for a gene with an infinity value for dN/dS (denoted as “dN/dS>1” and cited in the tables; that is, dN>0.001, and dS near or at 0); table 1, table 2, table3) was assigned a value of 1.5 for PGLS analysis to indicate that the value was larger than one, while conservatively ascribing a low range value (between one and infinity). If more than one branch had “dN/dS>1”, then the gene was excluded from the PGLS (one case, FBgn0002868 in table 3). Any species branch for a gene without a dN/dS value (dN and dS<0.001) was assigned “N/A” for PGLS.

#### *tau* Analysis

The *tau* value is a relative measure, and can vary with the number of tissues/stages studied. For instance, we found that *tau* values across all studied genes in the genome determined using all 59 tissues/stages yielded elevated values as compared to using only the 30 developmental stages alone (median= 0.830 versus 0.727, fig. S2), and the two sets of values were strongly correlated ($R=0.873$, $P<10^{-6}$) suggesting that both values effectively reflect *tau*. Throughout this study, we used the complete available dataset of 59 tissues/stages for all *tau* analyses, and considered genes with values >0.90 as narrowly transcribed (just below the 75$^{th}$ percentile), while those with values below 0.73 (at the 25$^{th}$ percentile) were considered as broadly transcribed, with all other values considered relative to each other and intermediate in *tau*.

### Supplementary Methods for the Three-Species Hawaiian Clade

As a supplementary analyses we studied a three species clade of Hawaiian *Drosophila*, shown in fig. S5 (phylogeny from (Kim, et al. 2021; Suvorov, et al. 2022)), that included *D. sproati* (Dspr, N=65.6 ovarioles), *D. murphyi* (Dmur, N=41.6 ovarioles) and *D. grimshawi* (47.8 ovarioles) (ovariole numbers from (Starmer, et al. 2003; Sarikaya, et al. 2019). For *D. sproati* and *D. murphyi* and the genomes were available as unannotated scaffolds from the SRA (Starmer, et al. 2003; Suvorov, et al. 2022) and the closely related species *D. grimshawi* was used as a reference (that has an annotated genome, N=12,825 CDS; longest CDS per gene; all three species genome data under project PRJNA675888 at the SRA database). Using the program Maker2 (Holt and Yandell 2011) and the RNA and proteins sequence list for *D. grimshawi*, we searched *D. sproati* and *D. murphyi* scaffolds for evidence-based gene predictions. We also obtained ab initio gene predictions in Augustus (Hoff and Stanke 2013) which recommends *D. melanogaster* gene models as a reference for predictions in Dipteran species (Stanke and Waack 2003). In the assessments, we excluded any CDS that lacked a start or a stop codon, or had an internal stop codon, such that only complete CDS were used for study. Using these criteria, we identified 13,395 CDS for *D. murphyi* and 12,670 CDS for *D. sproati* . Thus, these comprise high quality full-length CDS. To assess the completeness of our final *D. sproati* and *D. murphyi* CDS lists we applied BUSCO v. 5.5.2 (Seppey, et al. 2019; Manni, et al. 2021), a program that evaluates the completeness of a given transcriptome by comparing it to a subset of "universal" single-copy genes. For this, we used as the control the Dipteran gene set, named diptera_odb10 (N genes=3,250). The results showed that for the 13,395 full-length CDS for *D. murphyi*, 93.4% had a complete (unfragmented) match in the BUSCO dataset, indicating we obtained a very complete CDS list for this species (note that this value would be elevated using all 14,108 CDS identified for *D. murphyi*, but we focused solely on the obtained full-length CDS). In turn, for *D. sproati* , we found 93.2% of the full-length CDS were represented in the Dipteran BUSCO dataset. Taken together, the results show that our approach yielded a high confidence and highly complete CDS list for *D. sproati* and *D. murphyi.*

We next used reciprocal BLASTX (BLAST v. 2.13.0+ (Altschul, et al. 1997)) to identify orthologs between all three Hawaiian species using all CDS per genome in each contrast, for each combination of paired species, namely *D. grimshawi-D. murphyi*, *D. grimshawi-D. sproati* and *D. murphyi-D. sproati*. For each pair, in order to be identified as orthologs, two CDS had to be the best match (lowest e-score, any matched with a tied e-score the best match was the one with the highest bit value, $e<10^{-6}$) in both the forward and reverse contrast and have an e-value $<10^{-6}$. Using the three sets of paired orthologs, we then compared the orthologs identified in the newly annotated *D. murphyi-D. sproati* pair to the *D. grimshawi*.-*D. murphyi* and *D. grimshawi*. -*D. murphyi* pairs, to ensure that the orthologs per gene identified between *D. murphyi* and *D. sproati* each independently matched the same single CDS in *D. grimshawi*. Thus, this comprises very stringent criteria for identification of three-way reciprocal orthologs among *D. grimshawi*.-*D. murphyi*-*D. sproati*.

To identify Hawaiian species orthologs to the genes linked to ovariole/egg number evolution obtained from analysis within the *melanogaster* subgroup (SIGNALC in table 1;

SIGNALC in fig. 4), and for genome-wide orthologs, we compared the CDS of the most well annotated *melanogaster* species *D. melanogaster* (13,986 coding genes with longest isoform per gene) to the annotated Hawaiian species *D. grimshawi* CDS lists using reciprocal BLASTX (using same criteria described for Hawaiian *Drosophila* above). We report that 11,011 high confidence orthologs were identified between *D. melanogaster* and *D. grimshawi* (78.7% of the Dmel CDS list). A total of 10,276 of those genes had high confidence three-species orthologs in Hawaiian clade and were used for study of dN/dS and ortholog detection rates.

Each set of three-species orthologs among *D. grimshawi*.-*D. murphyi*-*D. sproati* were aligned using MUSCLE (Edgar 2004) set to default parameters in MEGA (Kumar, et al. 2016). Each alignment was filtered in GBlocks v. 0.91b set at default parameters (Castresana 2000; Talavera and Castresana 2007) to remove gaps and any highly divergent segments. For each aligned CDS in *D. grimshawi*, *D. murphyi* and *D. sproati*(gaps removed ), we determined dN/dS per species branch using the free-ratios model of PAML (M1) and conducted branch-site analyses (Yang 2007), as outlined for the *melanogaster* subgroup in the main text. Branches were unsaturated in substitutions, and the $90^{th}$ percentile of dS values across all genes was <0.090 and of dN was <0.032 for each of the three Hawaiian species branches. The median gene-wide dN/dS value across all genes for each species branch were 0.152, 0.164, and 0.160 for *D. murphyi*, *D. sproati* and *D. grimshawi* respectively.

#### Supplementary Results for the *Drosophila melanogaster* subgroup

##### Descriptions of Genes Linked to Ovariole Evolution

Each of the ovariole-related genes listed in the tables herein has properties suggesting a role in interspecies ovariole number divergence. In the following sections, we highlight some of these genes, which we consider especially promising candidates for functional analyses in future work, and briefly describe published evidence for their role in ovariole number determination and/or interspecies ovariole number divergence. We note that none of the 42 ovariole-related genes identified herein as linked to interspecies divergence in ovariole numbers (tables 1-5, fig. 4) overlapped with any of the 24 genes identified using a recent genome-wide association assessment of variation in 205 lines of the *D. melanogaster* Raleigh population (Lobell, et al. 2017). This is unsurprising, given the latter study was focused on intraspecies variation and ovariole numbers in a single *D. melanogaster* population (unfixed variation), while the present study focused on fixed nonsynonymous (to synonymous, dN/dS) and adaptive changes at the interspecies level

###### *SIGNALC genes (table 1)*

*<u>unpaired 2 (unp2)</u>*

The gene *upd2* is a ligand for the JAK/STAT pathway, which interacts with the Hippo pathway during ovariole development (Sarikaya and Extavour 2015). Of the five species in the *melanogaster* subgroup, *upd2* exhibited the highest terminal branch dN/dS value in *D. sechellia* (value of 0.417; table 1) and had signals of branch-site positive selection ($P<0.05$) in this same lineage (and in *D. erecta*, table 1). Thus, the gene protein changes in *D. sechellia* were coupled with the fact that this species has the lowest ovariole number of the species examined (fig. 2). *unp2* also exhibited the highest *tau* value (0.962) of all 27 studied SIGNALC genes, indicating that this gene is narrowly transcribed, which could potentially provide flexibility to adaptively evolve new functions in the ovarioles (table 1; (Otto 2004; Larracuente, et al. 2008; Mank and Ellegren 2009; Whittle, et al. 2021)). Upon inspection, we found that *upd2* was maximally transcribed ($\hat{x}$) in 4-6 hour old *D. melanogaster* embryos, which is also the same stage at which *hpo*, a known effector of ovariole numbers in larval ovaries (Sarikaya and Extavour 2015) is reported as maximally transcribed (Li, et al. 2014). The 4-6 hour old embryos comprise an embryonic stage during which the somatic gonad precursors (SGPs) are being established (Richardson and Lehmann 2010), and thus speculatively these two genes (*upd2* and *hpo*) could potentially influence ovariole numbers by affecting SGP number or behaviour (King, et al. 1968; Sarikaya, et al. 2012; Sarikaya and Extavour 2015). In sum, *upd2* protein sequence changes may contribute to the divergence in ovariole numbers in the *melanogaster* subgroup, particularly the marked decline in the *D. sechellia* branch.

*Zyxin (Zyx)*

For the SIGNALC ovariole-related gene *Zyx*, which modulates activity of the Hippo pathway (table S3; Rauskolb et al. 2011; Gaspar et al. 2014; Ma et al. 2016), each of the five species of the *melanogaster* subgroup had a gene-wide dN/dS value (ranging from 0.267 up to >1, table 1) that was more than threefold higher than the genome-wide median per respective species branch (the genome-wide median dN/dS values for each of the five branches are shown in fig. S1). The patterns suggest potential functional divergence of *Zyx* throughout the clade. *Zyx* exhibited gene-wide dN/dS>1 in *D. sechellia*, suggesting a history of positive selection in that lineage, that co-occurred with its marked decline in ovariole numbers (fig. 2), suggesting the potential for a causative relationship. Upon close examination, we found that *Zyx* in the *D. sechellia* branch was a case when dS approached 0 (the other species branches had dS between 0.006 to 0.010), however the dN value was 0.004 (above the cut-off of 0.001), which was double the dN for *D. simulans* and *D. melanogaster* (even though dS was higher for the latter two species, their dN was lower), together indicating that the elevated dN/dS in *D. sechellia* was due to elevated dN (not low dS, and is conservatively denoted as dN/dS>1). Overall, the patterns suggest positive selection of *Zyx* in *D. sechellia*.

There are plausible cell biological mechanisms whereby adaptive changes in the *Zyx* protein may potentially influence ovariole number divergence. The gene *Zyx* encodes an actin cytoskeleton regulator, with roles in Hippo pathway regulation (table S3, (Rauskolb, et al. 2011)), and evidence suggests it can regulate cell behaviors and movements during development (Amsellem, et al. 2005; Matsui and Lai 2013). Given that actin cytoskeleton activity is a prime candidate for coordinating the cell movements required for the formation of the TFs (Chen, et al. 2001), it may be speculated that Zyx may contribute to this process, and thereby changes in its amino acid sequence may affect TF formation (table 1), and in turn, alter the ovariole numbers (King, et al. 1968; Sarikaya, et al. 2012; Sarikaya and Extavour 2015). This provides a possible mechanism whereby functional changes in the Zyx protein may affect interspecies transitions in ovariole numbers. Further, *Zyx* positively regulates Yki (Rauskolb, et al. 2011), a transcriptional coactivator involved in many processes including cell proliferation (Huang, et al. 2005), and that accumulates in the nucleus in *hpo* RNAi larval ovary somatic cells (Sarikaya and Extavour 2015). We previously showed that elevating Yki levels in the somatic ovary results in an increased number of TFs and ovarioles (Sarikaya and Extavour 2015). Moreover, loss of *Zyx* reduces Yki activity (Rauskolb, et al. 2011). Thus, taken together, it may be hypothesized that the low ovariole numbers in *D. sechellia* as compared to all other *melanogaster* subgroup species (fig. 2) may have arisen from a reduction of Yki activity, that was caused by lowered function/activity of Zyx protein due to its amino acid sequence changes (table 1). Given that Z*yx* exhibited signs of positive selection in the *D. sechellia* branch (gene-wide dN/dS, table 1), this would in theory imply that the reduction in ovariole numbers (rather than an increase) in *D. sechellia* may comprise an adaptive reproductive change.

*Phosphatase and tensin homolog (Pten)*

The SIGNALC ovariole-related gene *Pten* (Kumar, et al. 2020) is a regulator of the mTOR and insulin signalling pathways (table S3). *Pten* exhibited an exceptionally high dN/dS value (0.594) in the *D. erecta* branch, more than five-fold higher than the *D. erecta* genome-wide median (0.104, fig. S1) and had branch-site positive selection (in *D. erecta*, $P<0.05$). This gene also exhibited a markedly elevated dN/dS value in the *D. melanogaster* branch (0.3122, table 1) and showed positive selection using the McDonald and Kreitman (1991) tests that were based on inter- versus intraspecies divergence solely for *D. melanogaster-D. simulans* ($P<0.012$, see Methods, and table 1 Notes (Murga-Moreno, et al. 2019)). *D. erecta* has an intermediate number of ovarioles per female (value of 27, fig. 2) among the five-species of the *melanogaster* subgroup (fig. 2) and forms an outgroup clade with *D. yakuba* (ovariole number 25.8). Similar to numerous other ovariole-related SIGNALC genes in table 1, *Pten* has core functions in regulating the actin cytoskeleton (von Stein, et al. 2005) (table S3), and is involved in cell size, cell growth and tumor suppression (Goberdhan, et al. 1999). Mutations in *Pten* impair cytoskeletal activity in *Drosophila* ovaries, and induce various female reproductive phenotypes including impaired localization of the germ plasm components *oskar* mRNA and Vasa protein in laid eggs and an absence of pole cells (von Stein, et al. 2005). Further, *Pten* RNAi in the *D. melanogaster* larval ovaries increases ovariole number (the *hpo[RNAi]* Ovariole Number phenotype (Kumar, et al. 2020), see table 1). Among all of the 59 tissues/stages used for *tau* analyses (which was a relatively low value of 0.663 in *D melanogaster* for this gene), *Pten* had maximal expression, $\hat{x}$=1, for the adult virgin ovaries (table S3), which is also consistent with an essential female ovarian role. In sum, given its functions in female reproductive system, and the rapid evolution of this signalling gene in four of five of the *Drosophila* lineages (0.128-0.594), and explicit evidence of positive selection in *D. erecta* (table 1), we hypothesize that *Pten* is a putative factor involved in ovariole number transitions in the *melanogaster* subgroup, and that may shape ovariole numbers by adaptive evolution, particularly in the *D. erecta* branch (table 1).

*vein (vn)*

The ovariole-related EGF pathway ligand gene *vn* (table S3) exhibited a dN/dS value that was between 1.6 to 7.2 fold higher than the genome-wide median per respective species branch in three of the *Drosophila* species branches studied (*D. simulans*, *D. sechellia* and *D. yakuba* had values of 0.4712, 0.2069, and 0.2802 respectively; table 1; genome-wide medians in fig. S1), and presented signals of branch-site positive selection in the *D. yakuba* branch ($P<0.05$, table 1). *vn* is involved in growth and tissue patterning, including roles in the *D. melanogaster* adult ovary in somatic escort cells and in the differentiation of female germ stem cells (Slaidina, et al. 2021; Tu, et al. 2021). Further, this gene is expressed in the *D. melanogaster* TFs and SH cells (Slaidina, et al. 2020), which regulate ovariole formation and number (King, et al. 1968; Sarikaya, et al. 2012; Sarikaya and Extavour 2015; Slaidina, et al. 2020), consistent with the finding that *vn* RNAi reduces ovariole number in *D. melanogaster* (the *hpo[RNAi]* Ovariole Number phenotype, table 1 (Kumar, et al. 2020)). The highly elevated dN/dS value for *vn* in both *D. simulans* and *D. sechellia* terminal branches in particular suggest that *vn* protein sequence evolution may contribute to the rapid and two-fold level change in ovariole numbers between these closely related species (fig. 2).

In sum, *vn* has signs of evolvability in its protein sequence, with frequent protein sequence changes and adaptive evolution (in *D. yakuba*) and yet also exhibited relatively slow evolution in the *D. melanogaster* branch (dN/dS=0.0841), perhaps due to high pleiotropy (*tau*=0.741, table S3; note *tau* was measured in *D. melanogaster* as the reference), suggesting the gene has been subjected to strong purifying selection in that lineage. Thus, *vn* appears to have a dynamic evolutionary pattern, suggestive of potential roles in ovariole number divergence in the *D. simulans* and *D. sechellia* branches, and in *D. yakuba*.

***BULKSG genes (table 2)***

<u>*Insulin-like peptide 5 (Ipl5)*</u>

The ovariole-related gene *Ilp5* in table 2 was particularly notable in that it exhibited strong purifying selection in *D. melanogaster* (dN/dS=0.001) and had high dN/dS values in all other branches (values of 0.2932-0.5843 in *D. simulans*, *D. sechellia*, *D. yakuba*, *D. erecta*, each markedly higher than the genome-wide branch median values, which ranged from 0.066 to 0.125, fig. S1). It may be the case that gene redundancy has contributed to the divergence patterns for this gene. The *D. melanogaster Ilp* family has eight genes that have evolved a range of functions and tissue type specializations, including ovarian functions for *Ilp5* (Gronke, et al. 2010). In gene families, one gene member (or another) may pass through various stages of partial redundancy of function while still retaining the original gene function (Ohno 1970; Kirschner and Gerhart 1998), creating a window for relaxed constraint, and evolvability, of protein-coding regions and potentially giving rise to adaptive new functions or subfunctions, including gene-dosage functions (Kirschner and Gerhart 1998; Presgraves 2005; Conant and Wolfe 2008; Kuzmin, et al. 2022). Otherwise, without a function, the genes may accumulate mutations to form a pseudogene or become lost from the genome (Ohno 1970; Presgraves 2005; Conant and Wolfe 2008; Kuzmin, et al. 2022). Given the gene *Ilp5* was retained in the genomes of all five species here (table 2), it may be theorized that its rapid sequence divergence in four of five terminal branches may be associated with the evolution of altered functions that may affect ovariole formation. In fact, *llp5* has been linked to female re-mating frequency in *D. sechellia* (Wigby, et al. 2011), and thus sexual selection pressures could potentially contribute to its very high dN/dS in that branch (0.5843, table 2), and to the evolved decrease in ovariole numbers in this lineage (fig. 2). With respect to the comparatively slow evolution of this gene in *D. melanogaster*, a study of *Ilp5* knockouts found evidence for compensatory expression of *Ilp3* (Gronke et al. 2010). *Ilp5* loss of function conditions cause a decrease in egg laying but do not completely eliminate it (Kumar, et al. 2020), suggesting its redundancy with other *Ilp* family members (and gene-dose compensation) may limit the extent of the negative effects on egg production. Nonetheless, the overall effect on egg yield and thus likely on fitness (Gronke, et al. 2010) may explain the strong purifying selection and low dN/dS value that we observed for the *D. melanogaster* branch, particularly if a weaker effect of *Ilp5* mutations occurred in the other species. In sum, *Ilp5* presents a dynamic history in the *melanogaster* subgroup and may substantively contribute to ovariole number variation in this taxon.

*aquarius* (*aqrs*)

The BULKSG ovariole-related gene *aquarius* (*aqrs*) (table 2) had the highest gene-wide dN/dS in the *D. yakuba* branch (0.3029), followed by *D. erecta* (0.1972), suggesting particularly rapid changes in the outgroup species (fig. 2). Aqrs protein has been associated with the functionality of the male seminal fluid proteins (SFPs) and the sex peptide (SP) in *D. melanogaster* (Findlay, et al. 2014), products that are transferred to the female reproductive tract during copulation and act to increase egg production and decrease female receptivity to courtship (Findlay, et al. 2014). The observed high levels of *aqrs* transcripts in larval somatic ovary cells (table 2) suggests that the gene also plays some role in these female reproductive cells. In this regard, *aqrs* may be a strong target for male-female sexual conflict, which is a factor known to cause rapid evolution of female (and male) proteins and of reproductive traits such as egg production, and may ultimately promote speciation (Rice 1996; Arnqvist, et al. 2000; Swanson and Vacquier 2002; Clark, et al. 2009). Thus, the rapid sequence divergence observed here for *aqrs* may putatively contribute to interspecies ovariole number variation.

***SINGLEC Genes (table S7; fig. 4)***

*Ecdysone-inducible gene E1* (*ImpE1*), *sm*, and *CLIP-190*

In terms of the subset of rapidly evolving genes that we identified that were exclusively upregulated in TFs (TFa and/or TFp, and no other cell types, table S7 Notes), examples included *Ecdysone-inducible gene E1* (*ImpE1*), *smooth (sm*), and *Cytoplasmic linker protein 190* (*CLIP-190*), each of which exhibited positive selection in the *D. sechellia* branch (table S7). *ImpE1* has been linked to cell rearrangements during morphogenesis (Natzle, et al. 1988), *sm* is a nuclear ribonucleoprotein involved in diverse roles including muscle functions (Draper, et al. 2009), neuronal and chemosensory processes (Layalle, et al. 2005), and *CLIP-190* proteins coordinate binding between actins and microtubules (Lantz and Miller 1998), and may affect intracellular transport (Sanghavi, et al. 2012). Taken together, the rapidly evolving genes upregulated in TF cell types are involved in multiple cellular processes likely required for TF morphogenesis, and thus adaptive changes observed in these genes (fig. 4) may contribute to the proximate mechanisms underlying the rapid evolutionary divergence in ovariole numbers in *Drosophila*.

**Genes downregulated in the BULKSG soma cells versus germ cells (table S7)**

In terms of genes upregulated in the germ cells (and thus downregulated in the soma), the top ten genes with the highest degree of upregulation are shown in table S5. The genes exhibited elevated *tau* values ranging from 0.880 to 0.975, indicating a tendency of narrow transcription breadth (and several very narrow, with *tau* values above 0.950) for these genes, potentially reflecting high specialization to germ cell functions. Further, 8 of the 10 genes were largely uncharacterized genes (annotations as "CG number" only in table S5) based on available annotation from DAVID (Huang da, et al. 2009) and FlyBase (Gramates, et al. 2022). The two relatively well characterized genes *no long nerve cord* (*nolo*) and *orientation disruptor* (*ord*) play roles in the nerve cord, extracellular matrix, and meiosis (Huang da, et al. 2009). As the evidence to date suggests that somatic cells, rather than germ cells, are the principal regulators of ovariole number, we did not focus further on this particular genes of very highly and positively upregulated germ cell genes, and instead focused in the main text on the genes upregulated in somatic cells and with variable expression across larval ovary stages in tables 3 and 4.

**BULKSG for three stages of larval ovary development**

We also identified genes from the BULKSG dataset with respect to the data for three stages of larval ovary soma development (early, mid or late). The 30 genes with the highest degree of upregulation ($log_2$ change) in one stage (versus the other two) and that also exhibited rapid evolution (M0 dN /dS>0.20) are provided in table S6 (given the three types of comparisons, we included the top 30 genes, rather than ten as in tables 2, table S5). These genes were utilized in conjunction with sc-RNA (SINGLEC) in table 3.

#### Supplementary Results For The Three-Species Hawaiian *Drosophila* Clade

Table S10 contains the dN/dS results for the Hawaiian *Drosophil*a orthologs of the 27 SIGNALC genes identified in table 1 from the *melanogaster* subgroup (Kumar, et al. 2020), and are described in the main text. Here, we note that six of the 27 studied genes of interest from the *melanogaster* subgroup (from table 1) had high confidence orthologs (reciprocal BLASTX) identified between *D. melanogaster-D. grimshawi* but not among all three species of *D. murphyi-D. sproati-D. grimshawi*. This may reflect rapid evolution of these genes in Hawaiian *Drosophila*, gene loss from the genome, and/or genes missing from the annotation (or lacking a complete CDS that were removed), although the latter has been largely mitigated given the high BUSCO scores from each genome (see the Supplementary Methods section above). Nonetheless, the data in table S10 infers rapid evolution of many of the SIGNALC genes (table S10), including positive selection, in the Hawaiian taxa (for those that had high confidence orthologs).

The ovariole related genes identified from the SINGLEC dataset in fig. 4 (and fig. S4, table S7), are less apt to share functions in Hawaiian *Drosophila* (than SIGNALC) given the fast evolution of gene expression of reproductive genes, including in *Drosophila* (Ranz, et al. 2003; Whittle and Extavour 2019). Thus, we excluded analyses of dN/dS and branch-site positive selection for this group pending future data on sc-RNA sequence or bulk-RNA seq of those cell types in *D. grimshawi* or other closely related Hawaiian taxa. Nonetheless, as cited in the main text, the genes that were highly upregulated in the *D. melanogaster* TF and SH cells (fig. 4, fig. S4, N values therein), were found to have the lowest frequency of high confidence ortholog gene sets identified in Hawaiian *Drosophila* among the nine cell types. The genes without orthologs included those from the *D. grimshawi-D. melanogaster* genome contrast (44.6% of all studied genes in fig. 4without Hawaiian orthologs were from this class) and from the genome contrasts among the three Hawaiian species (55.4% of all studied genes without Hawaiian orthologs were from this class, fig. S6). This pattern suggests that the genes upregulated in the TF and SH cells (in *D. melanogaster*) as a group have evolved rapidly across the studied *Drosophila* species making orthologs unrecognizable, and/or have more commonly exhibited gene gains and losses, than genes from other cell types (Tautz and Domazet-Loso 2011; Tautz, et al. 2013), as described in the main text.
